# Population-level annotation of lncRNA transcription in Arabidopsis reveals extensive variation associated with transposable element-like silencing

**DOI:** 10.1101/2023.03.14.532599

**Authors:** Aleksandra E. Kornienko, Viktoria Nizhynska, Almudena Molla Morales, Rahul Pisupati, Magnus Nordborg

**Affiliations:** Gregor Mendel Institute, Austrian Academy of Sciences, Vienna Biocenter, Dr. Bohr-Gasse 3, 1030, Vienna, Austria

**Keywords:** *Arabidopsis thaliana*, long non-coding RNAs, gene expression, lncRNA annotation, epigenetics, natural variation, transposon silencing

## Abstract

Long non-coding RNAs (lncRNAs) are understudied and underannotated in plants. In mammals, lncRNA loci are nearly as ubiquitous as protein-coding genes, and their expression is highly variable between individuals of the same species. Using *Arabidopsis thaliana* as a model, we aimed to understand the true scope of lncRNA transcription across plants from different regions and study its natural variation. We used transcriptome deep sequencing datasets spanning hundreds of natural accessions and several developmental stages to create a population-wide annotation of lncRNAs, revealing thousands of previously unannotated lncRNA loci. While lncRNA transcription is ubiquitous in the genome, most loci appear to be actively silenced and their expression is extremely variable between natural accessions. This high expression variability is largely caused by the high variability of repressive chromatin levels at lncRNA loci. High variability was particularly common for intergenic lncRNAs (lincRNAs), where pieces of transposable elements (TEs) present in 50% of these lincRNA loci are associated with increased silencing and variation, and such lncRNAs tend to be targeted by the TE silencing machinery. We create a population-wide lncRNA annotation in *A. thaliana* and improve our understanding of plant lncRNA genome biology, raising fundamental questions about what causes transcription and silencing across the genome.

**One-sentence summary:** lncRNA loci are plentiful in the *A. thaliana* genome, but their expression is extremely variable and largely repressed, with TE pieces enriched in intergenic lncRNAs aiding variability and silencing.

## Introduction

Long non-coding RNAs (lncRNAs) are a relatively new and still enigmatic class of genes that are increasingly recognized as important regulators participating in nearly every aspect of biology (Statello et al., 2021). There are more lncRNAs than protein-coding genes in the human (*Homo sapiens*) genome (Volders et al., 2019) and they are apparently abundant in the genomes of all eukaryotes (Mattick and Rinn, 2015; Kapusta and Feschotte, 2014). In human and mouse (*Mus musculus*), lncRNAs have been shown to be involved in various diseases (Wapinski and Chang, 2011; Batista and Chang, 2013), and medical applications have been proposed (Wahlestedt, 2013). Although many lncRNAs have demonstrated functions, most lncRNAs have not been studied (Leone and Santoro, 2016), and many knockouts of seemingly functional candidates showed no phenotypic differences relative to their respective wild types (Sauvageau et al., 2013), leading to continuous debate about the functionality and importance of lncRNAs as a gene class (Mattick et al., 2023). Evolutionary studies of lncRNAs have revealed low sequence conservation and highly divergent expression when compared to protein-coding genes (Necsulea and Kaessmann, 2014; Nelson et al., 2017), yet some signs of conservation and selection have also been found (Johnsson et al., 2014; Mattick et al., 2023). Several studies have looked at how lncRNAs differ between closely related species such as rat (*Rattus norvegicus*) and mouse (Kutter et al., 2012), human and chimp (*Pan troglodytes*) (Necsulea and Kaessmann, 2014) or different plant species (Nelson et al., 2017; Zhu et al., 2022), but few have looked at differences within the same species (Melé et al., 2015). It was recently shown that lncRNAs display salient interindividual expression variation in human (Kornienko et al., 2016) and mouse (Andergassen et al., 2017), much higher than that of protein-coding genes, but the meaning, causes and consequences of this high variability are unknown.

Arabidopsis (*Arabidopsis thaliana*) has higher natural genetic variability than humans (1001 Genomes Consortium, 2016) and represents an interesting and convenient model for studying lncRNA variation. As most research on lncRNAs has been performed in human and mouse (Rinn and Chang, 2020), relatively little is known about lncRNAs in plants (Liu et al., 2015; Budak et al., 2020). Several studies have identified and annotated lncRNAs in plant species such as Arabidopsis (Liu et al., 2012; Palos et al., 2022), wheat (*Triticum aestivum*) (Xin et al., 2011), maize (*Zea mays*) (Li et al., 2014) and strawberry (*Fragaria* × *vesca*) (Kang and Liu, 2015), but, although several databases have been created, the number and comprehensiveness of plant lncRNA annotations are often poorer than those of human and mouse (Zhu et al., 2022; Jin et al., 2021; Di Marsico et al., 2022; Szcześniak et al., 2019). Nonetheless, it is clear that lncRNAs do regulate genes in plants (Liu et al., 2015; Whittaker and Dean, 2017; Chen et al., 2023; Gullotta et al., 2023), and that lncRNA expression is particularly responsive to stress and environmental factors (Wang et al., 2017; Budak et al., 2020). Furthermore, this lncRNA response can be accession-specific to a much greater extent than that of protein-coding genes (Blein et al., 2020). Plant lncRNAs also affect significant crop traits and their relevance for food security has been highlighted (Gullotta et al., 2023). Understanding the real scope of lncRNA transcription in plants could help identify new candidates for functional studies and shed light on the genome biology of lncRNAs in plants and beyond (Palos et al., 2023).

While many lncRNAs have been shown to participate in epigenetic silencing or activation of protein-coding genes (Statello et al., 2021), much less research exists on the epigenetic regulation of lncRNAs themselves (Yang et al., 2023). In Arabidopsis, the epigenetic patterns of some functional lncRNAs have been thoroughly studied (Whittaker and Dean, 2017; Yang et al., 2023), but little is known about the epigenetics of lncRNAs on a genome-wide scale. While high epigenetic variation was reported between accessions (Kawakatsu et al., 2016), it is not clear how this variation affects lncRNAs.

LncRNAs are known to sometimes originate from transposable elements (TEs) (Zhu et al., 2022; Kapusta et al., 2013; Palos et al., 2022), yet what implications this origin has for their expression, epigenetics and variation is not well known. Similarly, while aberrant lncRNA copy number has been connected to disease and other phenotypes in human (*Homo sapiens*) (Xu et al., 2020; Athie et al., 2020), general information about lncRNA copy number and its consequences is missing, in particular in plants.

In this study, we aimed to study the extent and natural variability of lncRNA transcription in Arabidopsis. We annotated lncRNAs using data from 499 accessions, finding thousands of new lncRNA loci and generating an extended lncRNA annotation. We observed high expression and epigenetic variability for lncRNAs among accessions, with lncRNAs being generally silenced in any given accession. Epigenetic variability explains expression variation of many lncRNAs. Long intergenic ncRNAs (lincRNAs) showed particularly high variability and can be divided into protein-coding-like and TE-like loci that show differences in their epigenetic patterns, copy number and—most importantly—the presence of pieces of TE sequences. Indeed, such short pieces of TEs were prevalent in intergenic lncRNAs, likely attracting TE-like silencing to these loci. We provide new insights into the biology of lncRNAs in plants, identify a major role for TE-likeness in lncRNA silencing, and provide an extensive annotation and data resource for the Arabidopsis community.

## Results

### Transcriptome annotation from hundreds of accessions reveals thousands of previously unannotated lncRNAs

To investigate the extent of lncRNA transcription in Arabidopsis, we used newly generated and publicly available (Kawakatsu et al., 2016; Cortijo et al., 2019) polyA^+^ stranded transcriptome deep sequencing (RNA-seq) datasets spanning five different tissues/developmental stages (seedlings, 9-leaf rosettes, leaves from 14-leaf rosettes, flowers, and pollen) and 499 accessions (Figure 1A, see Supplemental Table S1 for accession list, Supplemental Table S2 for RNA-seq samples, and Supplemental Table S3 for RNA-seq mapping statistics). To create a cumulative transcriptome annotation, we mapped the RNA-seq data from all samples onto the TAIR10 genome, assembled transcriptomes from each accession/tissue separately (Supplemental Table S4) and then used a series of merging and filtering steps to generate one cumulative annotation, which we then classified into several gene classes (Figure 1B, Methods, Supplemental Figure S1). We used Araport11 and TAIR10 gene annotations (Cheng et al., 2017; TAIR10 annotation) to guide the classification of transcripts corresponding to protein-coding genes (PC genes), pseudogenes, TE genes and TE fragments, ribosomal RNA (rRNA) and transfer RNA (tRNA) loci, and used an additional protein-coding potential filtering step to identify a set of lncRNAs (Supplemental Figures S1, S2A). Our transcriptome annotation performed well in assembling known PC genes and known lncRNAs (Supplemental Figure S2B).

**Figure 1.**
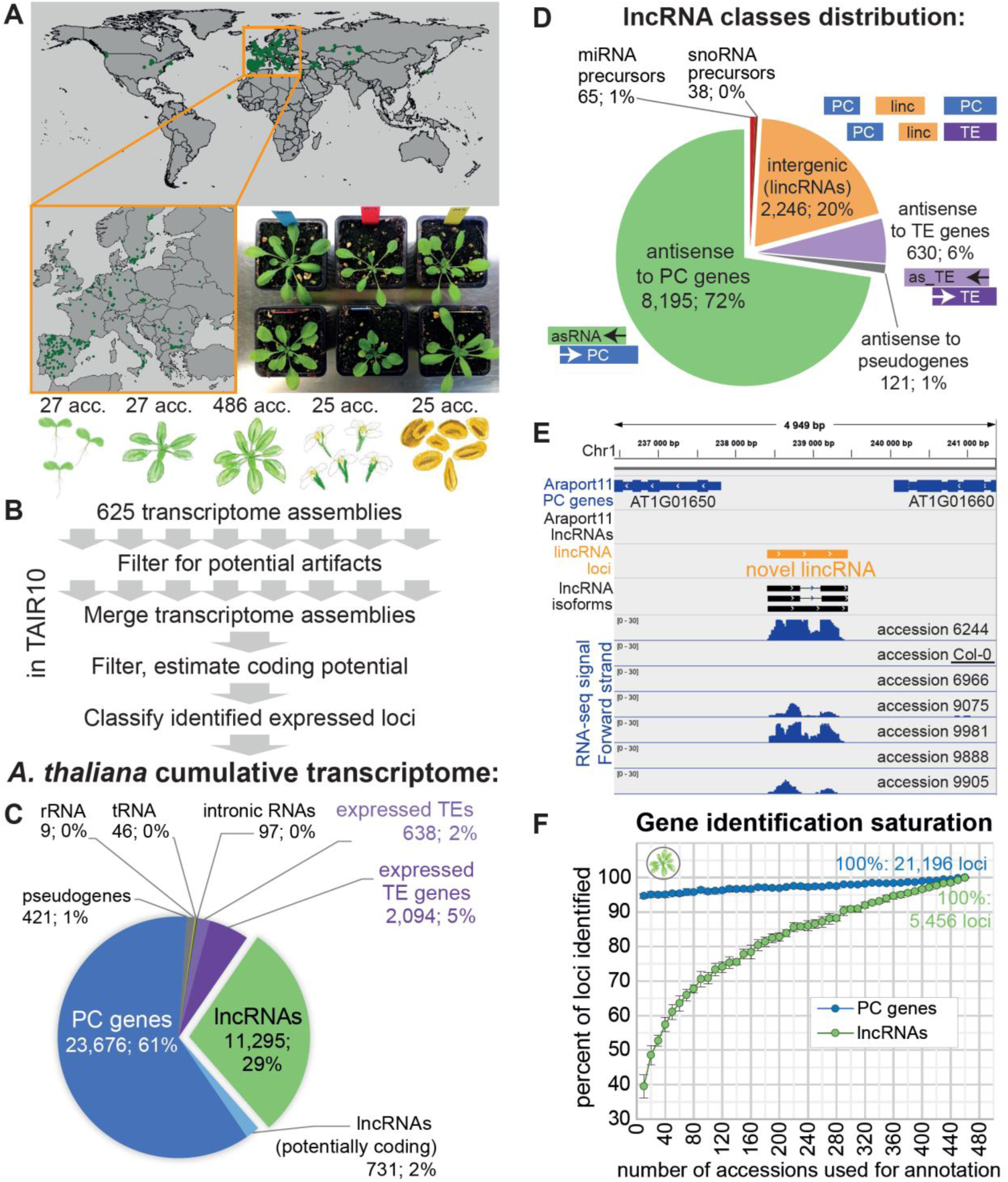
Mapping lncRNA transcription in hundreds of accessions and several tissues reveals thousands of previously unannotated lncRNAs. **A.** Origins of the Arabidopsis accessions used for transcriptome annotation and an example photograph of six different accessions grown in the growth chamber. **B.** Overview of the pipeline used for cumulative transcriptome annotation. Tissues from left to right: 7-d-old seedlings; 9-leaf rosettes; 14-leaf rosettes; flowers; pollen. **C.** Distribution of the types of loci in the cumulative annotation. **D.** Distribution of lncRNA positional classes in the genome. **E.** An example of a previously unannotated intergenic lncRNA on chromosome 1. Expression in seven different Arabidopsis accessions is shown. **F.** Number of lncRNA and PC loci identified as a function of the number of accessions used, relative to the number identified using 460 accessions. Random subsampling of accessions was performed in eight replicates and the error bars indicate the standard deviation across replicates.

In total, we identified 23,676 PC and 11,295 lncRNA loci, the latter thus representing almost one third (29%) of the cumulative transcriptome annotation (Figure 1C). The resulting annotation was highly enriched in lncRNAs (Supplemental Figure S2C) with 10,315 lncRNA loci (91%) being absent from the current public lncRNA annotation by Araport11. Our annotation extended the lncRNA portion of the reference genome from the 2.2% annotated in Araport11 to 10.7%, covering ∼13 Mb in total sequence (Supplemental Figure S2D). Comparison to the recent large-scale lncRNA identification studies in Arabidopsis (Zhao et al., 2018; Ivanov et al., 2021; Kindgren et al., 2020; Palos et al., 2022; Corona-Gomez et al., 2022) that, like Araport 11, were mainly based on the laboratory accession Columbia 0 (Col-0), showed that we identified 5,954 (53%) new lncRNA loci in the TAIR10 genome. We were also able to detect and annotate many TE genes and TE fragments across accessions/tissues (Figure 1C), finding spliced isoforms for 579 TE genes previously annotated as single-exon (Supplemental Figure S2E-H).

We classified lncRNAs based on their genomic position (Figure 1D). The largest group (8,195, or 72%) were antisense (AS) lncRNAs that overlapped annotated PC genes in the antisense direction (Supplemental Figure S3) (Ietswaart et al., 2012). We observed that 8,083 Araport11 PC genes have an antisense RNA partner, over five times more than in the Araport11 reference annotation. Previous studies have reported ubiquitous, unstable antisense transcription (Yuan et al., 2015; Li et al., 2013) as well as the activation of antisense transcription upon stress in Arabidopsis and other plants (Xu et al., 2021; Zhao et al., 2018), however, our data contained exclusively polyA^+^ RNA-seq datasets of seedlings and plants grown under normal conditions, so we can conclude that relatively stable polyadenylated antisense transcripts can be produced over almost a third of PC genes in Arabidopsis across different accessions and tissues.

The second largest class with 2,246 loci (20%) was long intergenic ncRNAs (lincRNAs) (Figure 1D-E). The third largest class (630, or 6%) consisted of lncRNAs that were antisense to TE genes (AS-to-TE lncRNAs). The remaining three classes constituted <3% of all lncRNA loci and we will ignore them below.

The genomic distribution of AS lncRNAs and AS-to-TE lncRNAs mirrored the annotation of PC and TE genes, respectively, with the former being enriched in chromosome arms, and the latter near centromeres. LincRNAs were also enriched near centromeres but were distributed across the genome (Supplemental Figure S4).

### Analyzing more accessions and tissues reveals more lncRNA loci

We hypothesized that a major reason for our discovering so many previously non-annotated lncRNA loci was that our annotation was based on hundreds of accessions, while most previous studies had only used the reference accession Col-0 (Cheng et al., 2017; Yuan et al., 2015, 2016). If lncRNAs are very variably expressed between individuals, as has been shown in humans (Kornienko et al., 2016), data from a single accession would uncover only the subset expressed in that particular accession. To test this idea, we subsampled the unified rosette RNA-seq dataset from the 1001 Genome project (Kawakatsu et al., 2016) and ran our annotation pipeline many times (Methods). This saturation analysis showed that the number of annotated lncRNA loci increases 2.5-fold by raising the number of accessions analyzed from 10 to 460 (Figure 1F). Unlike PC genes, the number of lncRNAs strongly depended on the sample size and showed no sign of saturating even with 460 accessions (Supplemental Figure S5A, B).

To confirm that the observed increase was not simply due to increased sequencing coverage, we compared these results with very high-coverage RNA-seq data from Col-0 only (Cortijo et al., 2019). Applying the same subsampling strategy to these data, we observed a much slower increase that saturated early and could not possibly explain the results in Figure 1F (Supplemental Figure S5C).

LncRNAs can be specific to certain tissues and developmental stages (Cabili et al., 2011; Palos et al., 2022; Liu et al., 2012), so it is likely that our use of multiple tissues helped identify more loci. Our final annotation, which was based on seedlings, rosettes, flowers, and pollen from multiple accessions (Figure 1B) revealed 11,265 lncRNA loci, while the 460-accession rosette analysis above returned only 5,456 lncRNA loci (Figure 1F). To better understand the effect of adding different tissues, we performed another saturation analysis where we varied both the number of tissues and accessions (Supplemental Figure S6). While the number of loci always increased with more accessions, the number of tissues used mattered even more. In particular, adding flowers or pollen to the analysis produced a big jump in the number of genes identified. For example, using data from 20 accessions and four tissues allowed the identification of ∼3 times more lncRNAs than when using just data from seedlings (Supplemental Figure S6). Adding flowers alone nearly doubled the number of lncRNAs identified.

To summarize, by combining RNA-seq data from hundreds of accessions and four developmental stages, we provide a massively expanded lncRNA annotation for Arabidopsis, identifying thousands of previously missed loci. Our extended lncRNA annotation is available in the supplement (Supplemental Data Set S1).

### High lncRNA expression variability between accessions

We showed above that including more accessions allowed the identification of more lncRNA loci (Figure 1F) and hypothesized that the reason was that not every lncRNA was expressed in every accession. Indeed, this appears to be the case: while most PC genes were expressed in nearly all accessions, most lncRNAs, as well as most TE genes and fragments, were expressed in fewer than 5% of all accessions (Figure 2A) (throughout this article we use “expressed” as in “detected” to refer to loci with transcripts per million mapped reads [TPM]>0.5 in a given dataset, aware of our inability to detect unstable or non-polyadenylated transcription using polyA^+^ data). Our analysis of the expression frequency (ON/OFF state) of the four main types of loci showed that while about 50% of PC loci are expressed in every accession, the same was true for no more than 1% of all AS lncRNAs, lincRNAs and TE genes (Supplemental Figure S7).

**Figure 2.**
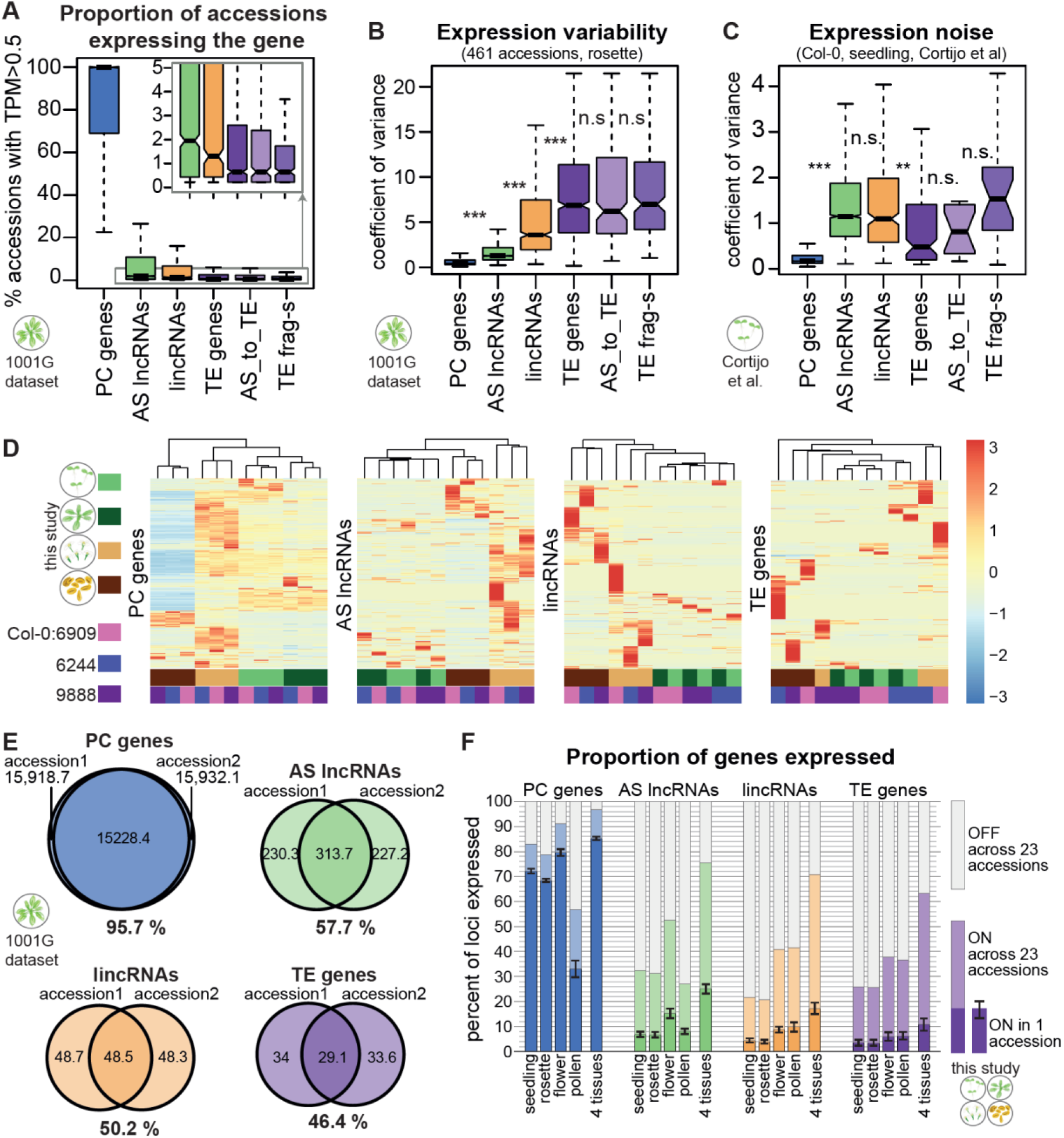
lncRNAs display extensive variability in their expression across accessions and appear to be largely silent. **A.** Proportion of accessions in the 1001 Genomes dataset (Kawakatsu et al., 2016) where the indicated type of gene is expressed (TPM > 0.5). Only genes that are expressed in at least one accession are plotted. **B.** Coefficient of variance of expression in 461 accessions from the 1001 Genomes dataset (Kawakatsu et al., 2016). Only genes with TPM > 1 in at least one accession are plotted. **C.** Expression noise, calculated from 14 technical replicates of Col-0 seedlings; expression noise values averaged across 12 samples are shown (Cortijo et al., 2019). Only genes with TPM > 1 in at least one sample are plotted. Boxplots: Outliers are not shown, and *P*-values were calculated using a Mann-Whitney test on equalized sample sizes: \*\*\**P*<10^−10^, \*\**P*<10^−5^, \**P*<0.01, n.s. *P*>0.01. **D.** Gene expression levels for different types of genes in four tissues for the reference accession Col-0 (accession number 6909) and two randomly picked accessions. Heatmaps were built using “pheatmap” in R with scaling by row. Only genes expressed in at least one sample are plotted. Clustering trees for rows not shown. **E.** Average number of genes expressed in an accession and its randomly selected partner accession from the 1001 Genomes dataset, and the number of genes expressed (TPM > 0.5) in both accessions. Percentages indicate the extent of overlap between accessions. **F.** Proportion of genes expressed in one accession in seedlings, 9-leaf rosettes, flowers, pollen, or all four tissues combined (dark bars). The error bars show standard deviation across the 23 accessions. The light part of the bars indicate the additional proportion of genes that can be detected as expressed when all 23 accessions are considered.

To quantify the natural variability in expression of lncRNAs and other gene types, we calculated the coefficient of variance using rosette RNA-seq data across 461 accessions (Kawakatsu et al., 2016). Similarly to human (Kornienko et al., 2016) and mouse (Andergassen et al., 2017), the expression of both AS lncRNAs and lincRNAs was significantly more variable than that of protein-coding genes (Figure 2B). In particular, lincRNAs showed expression variability almost to the level of TE genes and fragments. The variability in expression of lncRNAs that were antisense to TE genes was similar to that of TE genes and fragments (Figure 2B). Analyzing expression variability in other rosette RNA-seq datasets (Supplemental Figure S8A-B) and other tissues (Supplemental Figure S8C-E) confirmed these results.

Two factors crucially affect expression variability values and must be controlled for when comparing lncRNAs to PC genes: gene length and absolute expression level. lncRNAs are known to be shorter and have lower expression than PC genes (Cabili et al., 2011), which we confirmed in our data (Supplemental Figure S9A,B). Both gene length and absolute expression level were negatively correlated with the coefficient of variance; while this observation held true for all gene types, the anticorrelation slopes were different (Supplemental Figure S9C,D). When we controlled for expression level or gene length alone, the trend shown in Figure 2B was preserved (Supplemental Figure S10A,B). When we controlled for both expression and gene length, the trend was preserved for lincRNAs while AS lncRNAs were similar to PC genes (Supplemental Figure S9E,F), which might be explained by particularly high variability in the expression of short PC genes (Cortijo et al., 2019).

As the 14-leaf rosette dataset produced in this study contained 2–4 repeats for each accession, we could assess the level of intra-accession expression variation. For all classes of genes except AS lncRNAs, the intra-accession expression variation was significantly lower than the inter-accession variation; the difference between the classes of genes mirrored inter-accession variation (Supplemental Figure S10C). This result suggests that compared to PC genes, the expression of lincRNAs and TEs is more unstable and prone to be affected by the precise conditions or noise, while much of the AS lncRNA expression variation between accessions might be defined by generally unstable expression. To estimate the true noise in lncRNA expression, we analyzed the RNA-seq data consisting of Arabidopsis seedlings collected every 2 h over 24 h, with 14 technical replicates per time point (Cortijo et al., 2019). We determined that lncRNAs have significantly noisier expression (Figure 2C) as well as higher circadian expression variability (Supplemental Figure S10D). Interestingly, while TE genes showed higher variability between accessions (Figure 2B), both lincRNAs and AS lncRNAs were noisier than TE genes (Figure 2C).

To illustrate the extent of lncRNA expression variation, we plotted expression across four tissues in three accessions as a heatmap for different types of genes (Figure 2D). While PC gene expression levels clustered the samples according to tissue, lincRNA and TE expression clustered the samples according to accession (AS lncRNA was similar to PC gene but noisier). While pollen samples always clustered separately due to the particular transcriptome of pollen (Slotkin et al., 2009), the expression of lincRNAs and TEs was strikingly different between accessions. It was also notable that particularly many lincRNAs and TE genes were expressed in pollen, while flowers appeared to have higher expression for all four gene types. In general, Figure 2D illustrates how few lncRNAs are expressed in each accession and how striking the inter-accession variation is. Two randomly chosen accessions effectively express the same PC genes, whereas they share only about half of their expressed lncRNAs (Figure 2E). Furthermore, while ∼70% of PC genes were expressed in both seedlings and rosettes of any given accession, only 7% of AS lncRNAs and 4% of lincRNAs were expressed in these tissues (Figure 2F). Strikingly, almost twice as many AS lncRNA loci were expressed in flowers, but not in pollen, while twice as many lincRNAs were expressed in both flowers and pollen (Figure 2F). Like lincRNAs, TE genes showed increased expression in flowers and pollen, in agreement with a previous report (Slotkin et al., 2009). Across all four tissues in 23 accessions, 96% of PC genes, 76% of AS lncRNAs, 71% of lincRNAs, and 63% of TE genes were expressed (Figure 2F), thus covering many more lncRNAs, in line with the saturation analysis (Figure 1H) and the increased individual- and tissue-specificity of lncRNAs (Figure 2B,D).

In summary, lncRNA expression shows high variability between accessions, between tissues, and also between replicates. LincRNAs differ from AS lncRNAs in that they show higher expression variability and increased expression in pollen, but both classes are predominantly silent in any given sample.

### The epigenetic landscape of lncRNA loci suggests ubiquitous silencing

To characterize the epigenetic patterns of lncRNAs in Arabidopsis and investigate their apparently ubiquitous silencing, we performed chromatin immunoprecipitation followed by deep sequencing (ChIP-seq) and bisulfite sequencing in multiple accessions using leaves from rosettes at the 14-leaf stage (Methods, Supplemental Figure S11A, Supplemental Tables S5, S6). For the ChIP experiments, we chose two active marks (H3K4me3 and H3K36me3) and three repressive marks associated with different types of silencing (histone H1, H3K9me2 and H3K27me3) (Supplemental Figure S11B). Histone H1 was shown to be involved in silencing TEs, but also antisense transcription (Choi et al., 2020), H3K9me2 is a common heterochromatin mark and is known to silence TEs (Zemach et al., 2013), while H3K27me3 is commonly associated with polycomb repressive complex 2 (PRC2)-mediated silencing and is mostly deposited on PC genes (Feng and Jacobsen, 2011). H3K27me3 can also present found on some TEs when the normal silencing machinery is inactivated (Déléris et al., 2021; Zhao et al., 2022).

We first analyzed the ChIP-seq data focusing on the reference accession Col-0. The gene-body profiles of the different histone modifications were distinct for the different types of genes (Figure 3A, Supplemental Figure S11C). For example, while AS lncRNAs and PC genes showed similar levels of chromatin modifications—which is expected given their overlapping positions—their profiles differed, with PC genes showing a characteristic drop in H1 and H3K9me2 levels at their transcription start site (TSS) and an increase toward the transcription end site (TES), whereas AS lncRNAs showed an even distribution across the gene body (Figure 3A). LincRNAs and TE genes showed increased levels for the heterochromatic marks H3K9me2 and H1 and lower levels for the active marks H3K36me3 and H3K4me3 (Figure 3A). Calculating normalized and replicate-averaged ChIP-seq coverage over the entire locus (Figure 3B, Supplemental Figure S11D) and the promoter region (Supplemental Figure S11E) confirmed the above observations. Overall, AS lncRNAs were similar to PC genes, while lincRNAs were intermediate between PC genes and TE genes in their heterochromatic marks and the lowest in their active marks (Figure 3B, Supplemental Figure S11D, E).

**Figure 3.**
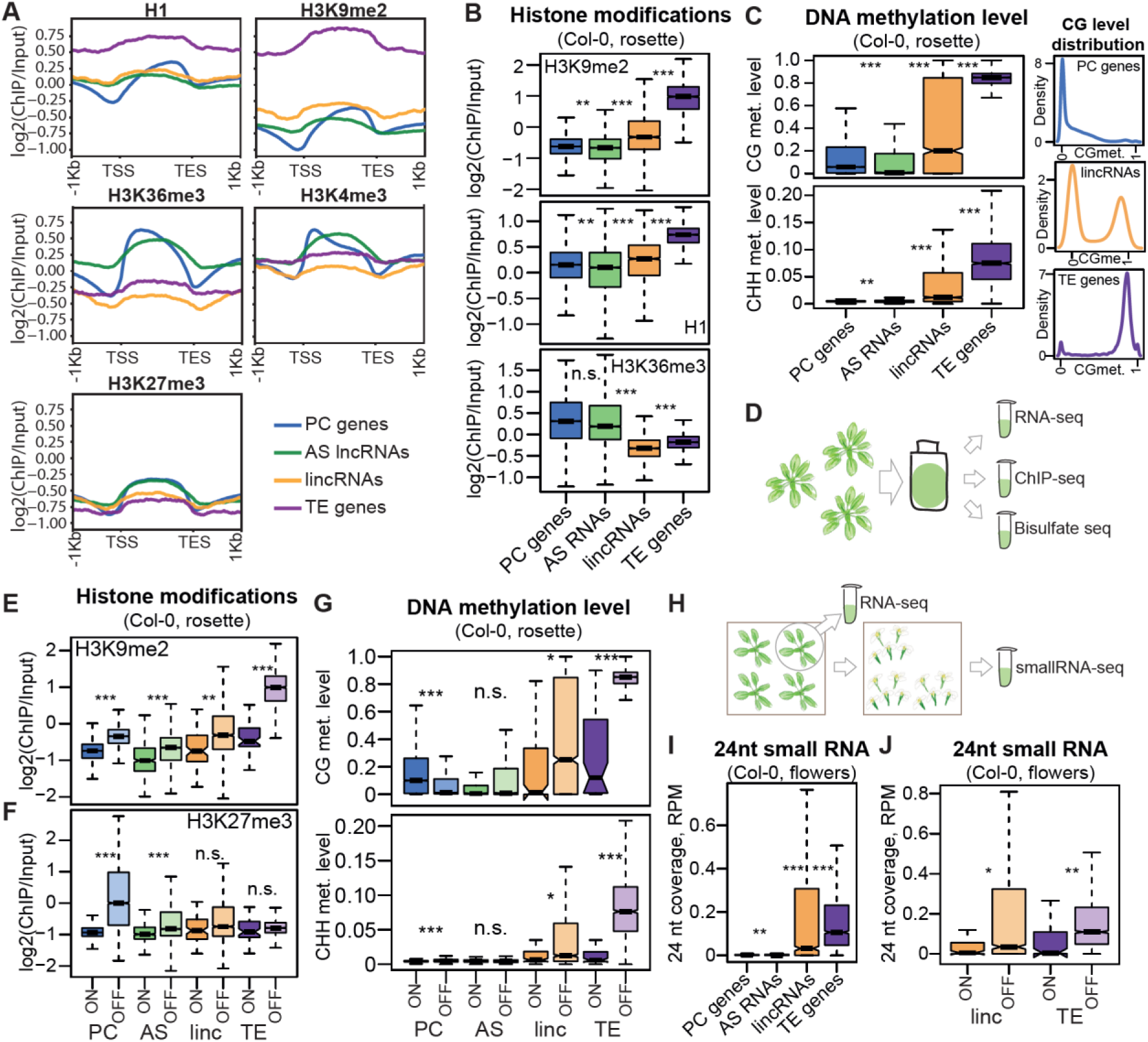
Epigenetic patterns of lncRNAs in Arabidopsis indicate their ubiquitous silencing. **A.** Averaged profiles of the input-normalized ChIP-seq signal for the epigenetic marks histone H1, H3K9me2, H3K36me3, H3K4me3 and H3K27me3 over four gene types from our cumulative transcriptome annotation. The plots show data from Col-0 rosettes, replicate 2. All genes, expressed and silent in Col-0, are used for the analysis. Metaplot profiles were built using plotProfile from deeptools (Ramírez et al., 2016). **B.** H3K9me2, H1 and H3K36me3 histone modifications in Col-0 rosette. The log2 of the gene-body coverage normalized by input and averaged between the two replicates is plotted. **C.** Left, CG and CHH DNA methylation levels in Col-0 rosettes. Right, density of CG methylation levels for PC genes, lincRNA loci and TE genes. Methylation level is calculated as the ratio between the number of methylated and unmethylated reads over all Cs in the respective context (CG, CHH) in the gene body and averaged over four replicates. **D.** Diagram of the experiment: the same tissue from 14-leaf rosettes was used for RNA-seq, ChIP-seq and Bisulfite-seq in this study. **E.** H3K9me2 input-normalized coverage, plotted separately for expressed (ON, TPM>0.5) and silent genes (OFF, TPM<0.5). **F.** H3K27me3 normalized coverage, plotted separately for expressed (ON, TPM>0.5) and silent (OFF, TPM<0.5) genes. **G.** Methylation levels for expressed (ON, TPM>0.5) and silent (OFF, TPM<0.5) genes. Expression was calculated in the corresponding 14-leaf rosette samples. **H.** Diagram of the experiment: one 9-leaf rosette individual from the batch was used for RNA-seq and flowers for small RNA-seq were collected from the remaining individuals at a later point. **I.** Coverage of 24-nt small RNAs in the gene body, calculated as the number of 24-nt reads mapping to the locus and divided by the total number of reads and locus length. **J.** Coverage of 21–22-nt small RNA in Col-0 flowers, plotted separately for expressed (ON, TPM>0.5) and silent (OFF, TPM<0.5) genes. Expression was calculated in the corresponding 9-leaf rosette samples. *P*-values were calculated using a Mann-Whitney test on equalized sample sizes: \*\*\**P*<10^−10^, \*\**P*<10^−5^, \**P*<0.01, n.s. *P*>0.01. Outliers in the boxplots are not shown.

We then analyzed the bisulfite sequencing data, quantifying DNA methylation in three different contexts: CG, CHG, and CHH, where H stands for A, C, or T (Methods). CHG and CHH methylation are common in plants and are involved in TE silencing (Fultz et al., 2015). While PC genes and AS lncRNAs displayed low levels of CG methylation and no CHG or CHH methylation, as expected, lincRNAs exhibited a very significant methylation increase in all three contexts (Figure 3C, Supplemental Figure S12A, B). Interestingly, the distribution of CG-methylation over lincRNAs was bimodal (Figure 3C: right, Supplemental Figure S12C, D), with some loci looking like PC genes, while others looked like TE genes.

As TE genes and lincRNAs are enriched next to the centromeres while PC genes and AS RNAs are not (Supplemental Figure S4), we checked if the observed epigenetic differences held true when controlling for chromosomal position. While pericentromeric genes within 2 Mb of the centromeres all showed more heterochromatic patterns, the observed trends for histone marks and DNA methylation held true, especially for the genes further than 2 Mb from the centromeres (Supplemental Figures S13, S14).

To confirm that the repressive marks at lncRNA loci are associated with silencing, we checked for epigenetic differences between expressed and silent genes (the same samples were used for ChIP-seq, bisulfite sequencing, and RNA-seq, Figure 3D). The repressive marks H3K9me2 (Figure 3E, Supplemental Figure S15A) and H1 (Supplemental Figure S15B, C) were significantly more abundant on silent genes. While this result was true for all gene categories, the H3K9me2 difference for TE genes was particularly high, underscoring the fact that TEs are normally silenced by H3K9me2 deposition (Feng and Jacobsen, 2011). Another repressive mark, H3K27me3, also showed significantly higher levels on silent genes of all categories, but here PC genes showed a striking increase while TE genes were minimally different (Figure 3F, Supplemental Figure S15D). We also found that silent AS lncRNAs show increased CG methylation and that silent lincRNAs have strikingly increased CG and CHH methylation levels, although less so than TE genes (Figure 3.G, Supplemental Figure S16). Expressed PC genes had higher CG gene body methylation levels than silent ones, which is a known phenomenon of as yet unclear function (Bewick and Schmitz, 2017).

Since lincRNAs showed both CHH methylation and H3K9me2, both characteristic of TEs and absent from PC genes (Fultz et al., 2015), we performed small RNA (sRNA) sequencing of flowers (Figure 3H, Supplemental Table S7) to look for evidence of targeting by the 24-nucleotide (nt) small RNAs that are normally involved in TE silencing by the RNA-directed DNA methylation (RdDM) pathway (Matzke and Mosher, 2014). This analysis demonstrated that sRNAs indeed target many lincRNA loci (Figure 3I) (1,131 (50.4%) loci with RPM>0.03) and were associated with silencing for both lincRNAs and TE genes (Figure 3J). Analyzing published sRNA-seq data (Papareddy et al., 2020) from leaves and a very early embryonic stage known as “early heart” where RdDM-mediated silencing is particularly active (Papareddy et al., 2020) confirmed targeting of lincRNAs by 24-nt sRNAs, with samples at the early heart showing very high levels of sRNAs (Supplemental Figure S17A, B). An analysis of shorter 21–22-nt sRNAs, reported to also participate in TE silencing (Pontier et al., 2012), showed that lincRNAs also show higher levels of targeting by this type of siRNAs (Supplemental Figure S17C-E), however there was no clear association between the levels of these shorter siRNAs and the lack of expression of their corresponding lincRNA (Supplemental Figure S17F).

### Epigenetic variation explains expression variation of many lncRNAs

In the previous section, we described the epigenetic patterns only in the reference accession, Col-0. As we produced ChIP-seq, bisulfite-seq, and sRNA-seq data for several accessions (Supplemental Tables S5–S7) we were able to confirm that the epigenetic patterns we observed in Col-0 were similar in other accessions (Supplemental Figure S18). However, while the overall patterns were similar, the variability between accessions at particular loci was very high for both lncRNA (especially lincRNA) and TE genes (Figure 4A, B, Supplemental Figure S19).

**Figure 4.**
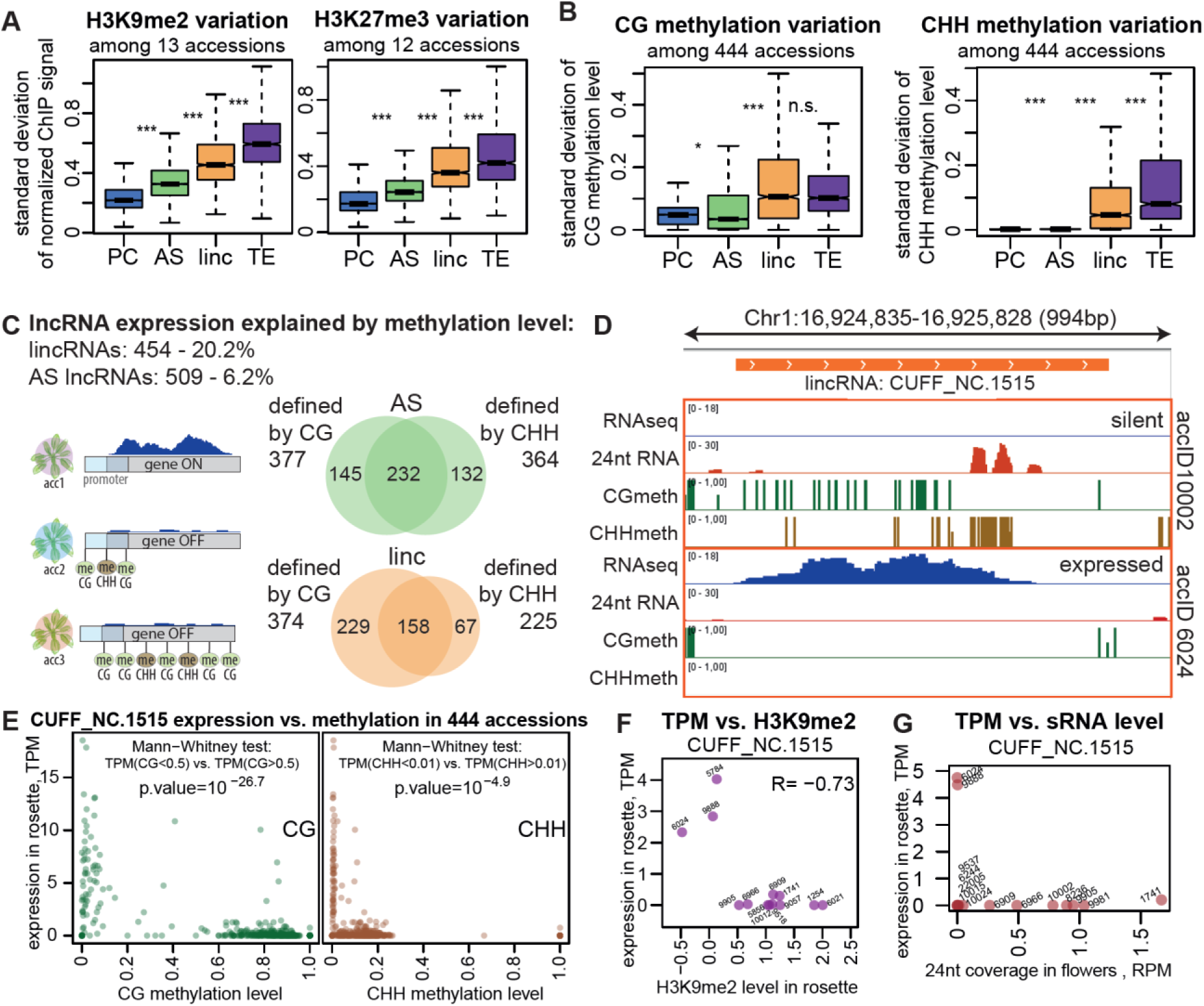
lncRNAs display increased epigenetic variation that explains the variation in expression of many lncRNAs. **A.** Standard deviation of input and quantile-normalized coverage (see Methods) of H3K9me2 (left) and H3K27me3 (right) signals in rosettes across 13 or 12 accessions, respectively. **B.** Standard deviation of CG (left) and CHH (right) methylation levels across 444 accessions (rosettes, 1001G dataset, (Kawakatsu et al., 2016)). *P*-values were calculated using Mann-Whitney test on equalized sample sizes: *** *P*<10^−10^, \*\**P*<10^−5^, \**P*<0.01, n.s. *P*>0.01. Outliers in the boxplots are not shown. **C.** Summary of lncRNAs for which expression can be explained by methylation (Supplemental Figure S20B). The Venn diagrams show the overlap between loci for AS lncRNAs (green) and lincRNAs (orange) that were found to be defined by CG or CHH methylation levels. **D.** An example of a lincRNA defined by CG and CHH methylation, showing RNA-seq signal (forward strand), CG and CHH methylation levels in rosettes, and the 24-nt sRNA signal in flowers in two accessions. **E.** Expression level as a function of CG (left) or CHH (right) methylation for the example lincRNA across 444 accessions (Kawakatsu et al., 2016). The results of the Mann-Whitney tests used for defining the explanatory power of CG/CHH methylation are shown. **F.** Expression in rosettes as a function of H3K9me2 level in rosettes of the example lincRNA in 13 accessions. **G.** Expression in rosettes as a function of 24-nt sRNA coverage in flowers of the example lincRNA in 14 accessions.

To test whether epigenetic variation can explain this variation in expression, we analyzed methylation patterns and expression data from rosettes collected from 444 accessions (Kawakatsu et al., 2016). We determined that for 454 lincRNAs and 509 AS lncRNAs, expression across accessions is indeed explained by the level of CG or CHH methylation at their gene body or promoter (Figure 4C, Supplemental Figure S20A, Methods). While these numbers correspond to only 20.2% and 6.2% of all lincRNAs and AS lncRNAs respectively, this analysis could only be performed on a limited number of informative loci with sufficiently high methylation variation and expression frequency (Supplemental Figure S20B, Methods). Among these informative loci, we could explain expression variation by variation in DNA methylation for 50.7% of lincRNAs and 21.5% of AS lncRNAs.

An example of such a lncRNA with high variation between accessions is displayed in Figure 4D: the accession that expresses the lincRNA lacks CG and CHH methylation in the locus as well as 24-nt sRNAs while the accession where the lincRNA is silent has both CG and CHH methylation, and 24-nt sRNAs in flowers. The epigenetic variation at this locus was extensive and quite binary with very strong association between the presence of methylation and the lack of expression and vice versa (Figure 4E). For the accessions with available data, the repressive histone modifications H1 (Supplemental Figure S20C) and H3K9me2 (Figure 4F), as well as 24-nt sRNA coverage (Figure 4G), were also anticorrelated with expression across accessions.

In summary, we establish that lncRNAs display distinctive epigenetic patterns, consistent with the above observation of the lack of expression and suggesting ubiquitous silencing. Compared to PC genes, lncRNAs display increased epigenetic variation between accessions that explained the variation in expression of ∼20% of lincRNAs and ∼6% of AS lncRNAs. Many lincRNAs show TE-like epigenetic status that is associated with silencing, as well as being targeted by 24-nt siRNAs, which are characteristic of the RdDM pathway for silencing TEs. Because of the interesting patterns and the outstanding variation observed for lincRNAs, we focused on them below.

### lincRNAs are enriched for TE pieces

Several similarities between lincRNAs and TE genes were apparent. First, both showed increased expression in flowers and pollen (Figure 2F). Second, lincRNA expression was dramatically more variable than that of PC genes and antisense lncRNAs — almost at the level of TE genes (Figure 2A). Third, our survey of the epigenetic landscape showed that lincRNAs display TE-like characteristics, although to a lesser extent (Figure 3A, B, F). Fourth, similarly to expression variation, levels of repressive chromatin at lincRNA loci were more variable than at PC genes and AS lncRNAs, trending toward the pattern seen in TE genes (Figure 4A, B). Furthermore, lncRNAs can originate from TEs and contain parts of their sequences, both in plants and animals (Kapusta et al., 2013; Palos et al., 2022; Corona-Gomez et al., 2022); moreover, TE domains within lncRNAs can play significant roles in lncRNA biology (Johnson and Guigó, 2014), such as their nuclear export or retention (Lubelsky and Ulitsky, 2018), or even have a crucial role in their function (Colognori et al., 2020). Accordingly, we asked if TE sequences contributed to lincRNA loci in Arabidopsis and affected their expression, epigenetics, and variability.

To this end, we used a BLAST-based analysis to identify sequences similar to TAIR10-annotated TEs inside loci and their borders (Figure 5A, Methods). We called each match a “TE piece” and merged overlapping same-direction TE pieces. We further refer to each TE-like region of a locus as a “TE patch”, no matter whether it was constituted by a single TE piece or several merged ones (Figure 5A). LincRNA loci were clearly enriched in TE patches compared to AS lncRNAs and PC genes, as well as randomly picked intergenic regions of corresponding length (Figure 5B, Methods). We determined that 52% of lincRNAs but only 27% of matching intergenic controls contain a TE patch. We also observed an enrichment for TE patches in upstream and downstream lincRNA border regions compared to matching controls (Figure 5B). On a per kb basis, lincRNA borders showed the highest density of TE patches, even higher than for lincRNA loci, and the difference between lincRNAs and other gene types became even more prominent (Figure 5C). TE genes had fewer TE patches per 1 kb than lincRNAs, presumably because TE genes usually contain one large TE patch corresponding to the full TE, while lincRNAs contained several smaller patches (Supplemental Figure S21A).

**Figure 5.**
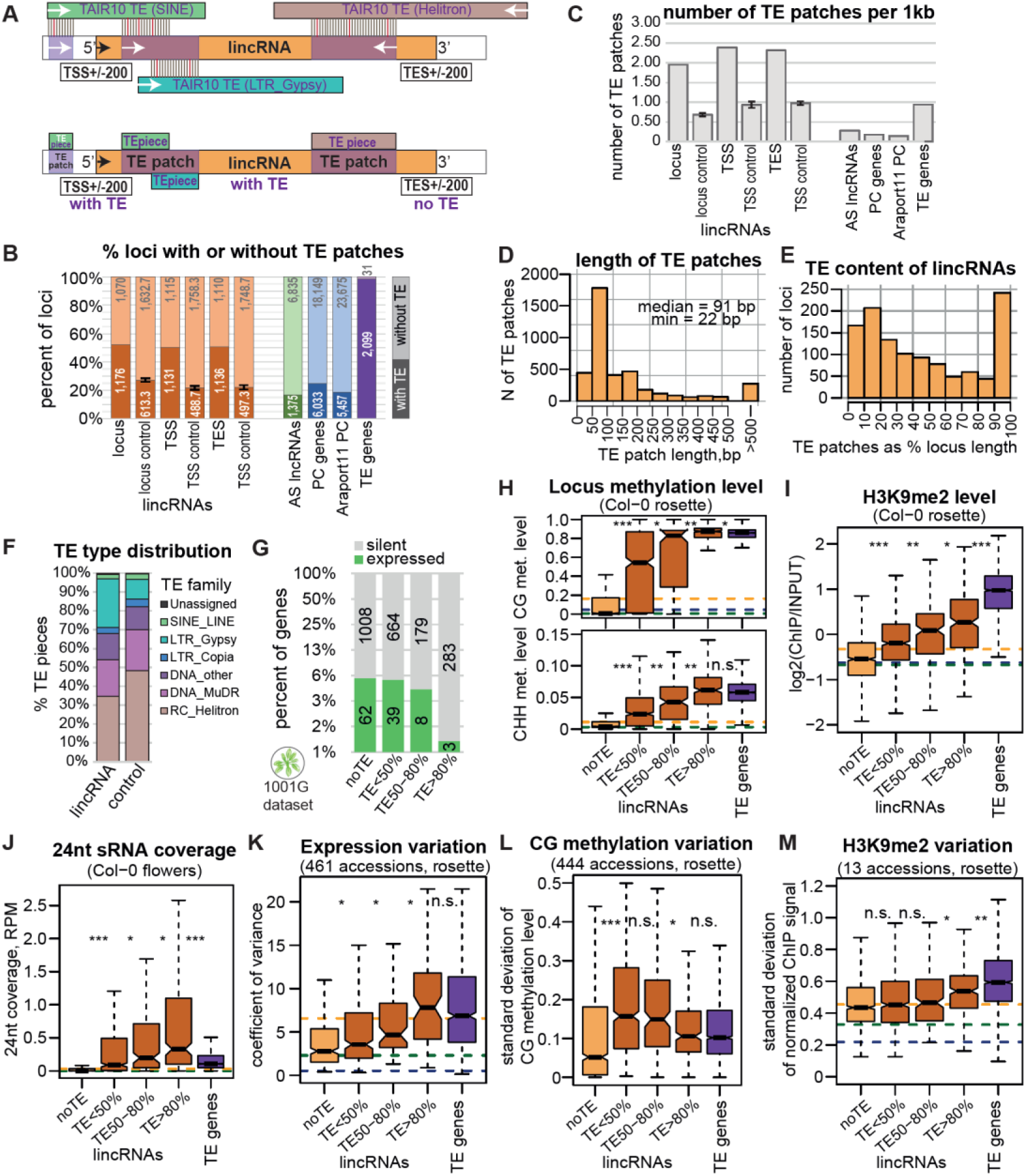
Many lincRNAs contain pieces of TEs that affect their silencing and expression variation. **A.** Outline of TE content analysis. Top, TAIR10-annotated TEs were compared to the sequences of lincRNAs (and other loci) by BLAST. Bottom, The mapped pieces of different TEs overlapping in the same direction were merged into “TE patches’’. The upstream and downstream “borders” of genes were analyzed in the same way. **B.** Proportion of loci containing a TE piece. The intergenic controls for lincRNAs, lincRNA transcription start site (TSS) ± 200 bp and transcription end site (TES) ± 200 bp were obtained by shuffling the corresponding loci within intergenic regions (lincRNAs excluded) three times and averaging the results. The error bars on controls represent the standard deviation between the three shuffling replicates. **C.** Number of TE patches per 1 kb. **D.** Distribution of the length of TE patches (any relative direction) within lincRNA loci. **E.** TE content distribution among lincRNAs. TE patches were considered in any relative direction. The loci with large TE content are those where the TE patches map antisense to the lincRNA locus. **F.** Proportion of TE pieces of different TE families within different types of loci. **G.** Proportion of expressed lincRNAs as a function of their TE content. The y-axis is displayed in log2 scale. **H–J.** Levels of methylation (**H**), H3K9me2 (**I**), and 24-nt sRNAs (**J**) for lincRNA loci as a function of their TE content, with TE genes for comparison. CG and CHH methylation data displayed are from Col-0 rosettes (Kawakatsu et al., 2016). **K–M.** Expression variability between 461 accessions (Kawakatsu et al., 2016) (**K**), standard deviation of CG methylation levels across 444 accessions (Kawakatsu et al., 2016) (**L**) and standard deviation of quantile- and input-normalized H3K9me2 levels in rosettes across 13 accessions (**M**) of lincRNA loci as a function of their TE content, with TE genes for comparison. *P*-values were calculated using Mann-Whitney tests: *** *P*<10^−10^, \*\**P*<10^−5^, \**P*<0.01, n.s. *P*>0.01. Outliers in the boxplots are not shown.

It is important to note that lincRNAs are not simply expressed TEs. Our lncRNA annotation pipeline required that a lncRNA did not overlap with any TE genes and allowed for a maximum of 60% same-strand exonic overlap with annotated TE fragments (Supplemental Figure S1). While only 486 (21.6%) of all lincRNA loci had a same-strand exonic overlap with a TAIR10-annotated TE fragment, 1176 (52.4%) contained a TE patch (Figure 5B), both sense and antisense to the lincRNA direction (Figure 5A, Supplemental Figure S21B). In addition, 136 (12%) of TE-containing lincRNA loci fully (>90%) overlapped with annotated TE fragments but were transcribed in the direction antisense to those (Supplemental Figure S21C,D). While 893 (76%) of the TE-patch-containing lincRNA loci overlapped with an annotated TE fragment (Supplemental Figure S21E,F), 472 in the sense and 605 in the antisense direction, 283 (24%) did not overlap with any annotated TEs but did contain a TE patch (Supplemental Figure S21G,H), that is contained pieces of sequences bearing resemblance (see Methods) to TE sequences. TE patches within lincRNAs (and other genes) were generally short with a median length of 91 bp and a minimal length of 22 bp (Figure 5D), thus much shorter than TAIR10-annotated TE fragments or the TE patches that our analysis identified in TE genes (Supplemental Figure S21I). Moreover, 74% of lincRNAs with multiple TE pieces contained pieces of TEs from different families, and 24% pieces of TEs from both Class I and Class II (Supplemental Figure S21F,G,H). The protein-coding potentials of lincRNAs and TE genes were also very different (Supplemental Figure S2A). Collectively, these results suggest that lincRNAs can be considered as a gene category separate from expressed TE fragments.

The relative TE content of TE-containing lincRNA loci differed greatly, with a few lincRNAs fully covered by TE patches, corresponding to lincRNAs that are antisense to a TE fragment (Figure 5E, Supplemental Figure S22A). On average, lincRNAs contained 309 bp of same-strand TE-like sequences and 436 bp of antisense TE-like sequences per kb. LincRNA loci were particularly enriched in TE sequences from the long terminal repeat (LTR) type TE *Gypsy* compared to matching intergenic controls and other gene types (Figure 5F, Supplemental Figure S22B), and this enrichment was particularly pronounced in lincRNAs with antisense TE patches (Supplemental Figure S22C-D). We did not, however, observe any particular localization for TE pieces from different families within the lincRNA loci (Supplemental Figure S22E-F).

### The TE content of lincRNAs affects their expression and epigenetics

We asked if the TE content of a lincRNA affected its expression, epigenetic characteristics, and their variability. Indeed, when binned based on relative TE sequence content, lincRNAs with higher TE content were less often expressed (Figure 5G) and showed higher levels of CG and CHH methylation (Figure 5H), H3K9me2 (Figure 5I) and 24-nt siRNAs (Figure 5J). The expression variation (Figure 5K, Supplemental Figure S23) and epigenetic variation (Figure 5L, M, Supplemental Figure S24) of lincRNAs also depended on their TE content, but not as strongly.

LincRNAs are enriched in the pericentromeric regions (Supplemental Figure S4) that are naturally enriched in TEs and heterochromatin, which might confound our TE piece (Figure 5B, C) and epigenetic analyses (Figure 5H-J). Controlling for the proximity to centromeres, we first discovered that while all gene types have higher TE content closer to centromeres, the increased TE content of lincRNAs observed in Figure 5B was preserved (Supplemental Figure S25A). Second, while all pericentromeric lincRNAs, even those without TE patches, showed high repressive chromatin, the level of heterochromatic marks at lincRNA loci further from centromeres strongly depended on their TE content (Supplemental Figure S25B-D). Furthermore, while 24-nt sRNA coverage was generally low near centromeres (consistent with previous findings, see (Sigman and Slotkin, 2016)), it strongly depended on the TE content in chromosome arms (Supplemental Figure S25E). Thus, the presence and the relative size of TE sequences inside lincRNA loci are indeed associated with a more repressive chromatin state irrespective of chromosomal location.

In summary, we showed that intergenic lncRNAs are highly enriched for short pieces of TEs. About half of all lincRNAs have a TE sequence within them, and higher TE content is associated with more repressive epigenetic marks when comparing different lincRNAs in the genome.

### Copy number of lincRNAs affects their expression variability and epigenetic patterns

Apart from expression variability, epigenetic patterns and TE sequence content, another classical TE feature was evident for lincRNAs: lincRNAs were often present in multiple copies (Supplemental Figure S26A). We decided to investigate this pattern further and see whether it affects their epigenetic patterns and expression.

We used a BLAST-based approach to look for multiple gene copies in TAIR10 (Methods) and found that lincRNAs are much more commonly multiplicated than PC genes and AS lncRNAs, with 28% being present in more than one copy and 8% in more than 10 copies (Figure 6A). Again, lincRNAs were intermediate between PC genes and TE genes. We split lincRNAs into four categories: single- or multi-copy lincRNAs, with or without TE patches (Figure 6B, top). Similarly to the overall lincRNA distribution (Figure 5B), about half of all single-copy lincRNAs contained a TE patch, while most multi-copy lincRNAs did (Figure 6B). LincRNAs with higher copy numbers also showed higher TE sequence content (Supplemental Figure S26B). Although *Helitron*s have the highest copy number in the Arabidopsis genome (Quesneville, 2020), lincRNA with pieces of *Gypsy* elements showed the highest copy number (Supplemental Figure S26C), even when the TE sequence content was no more than 20% of the locus (Supplemental Figure S26D).

**Figure 6.**
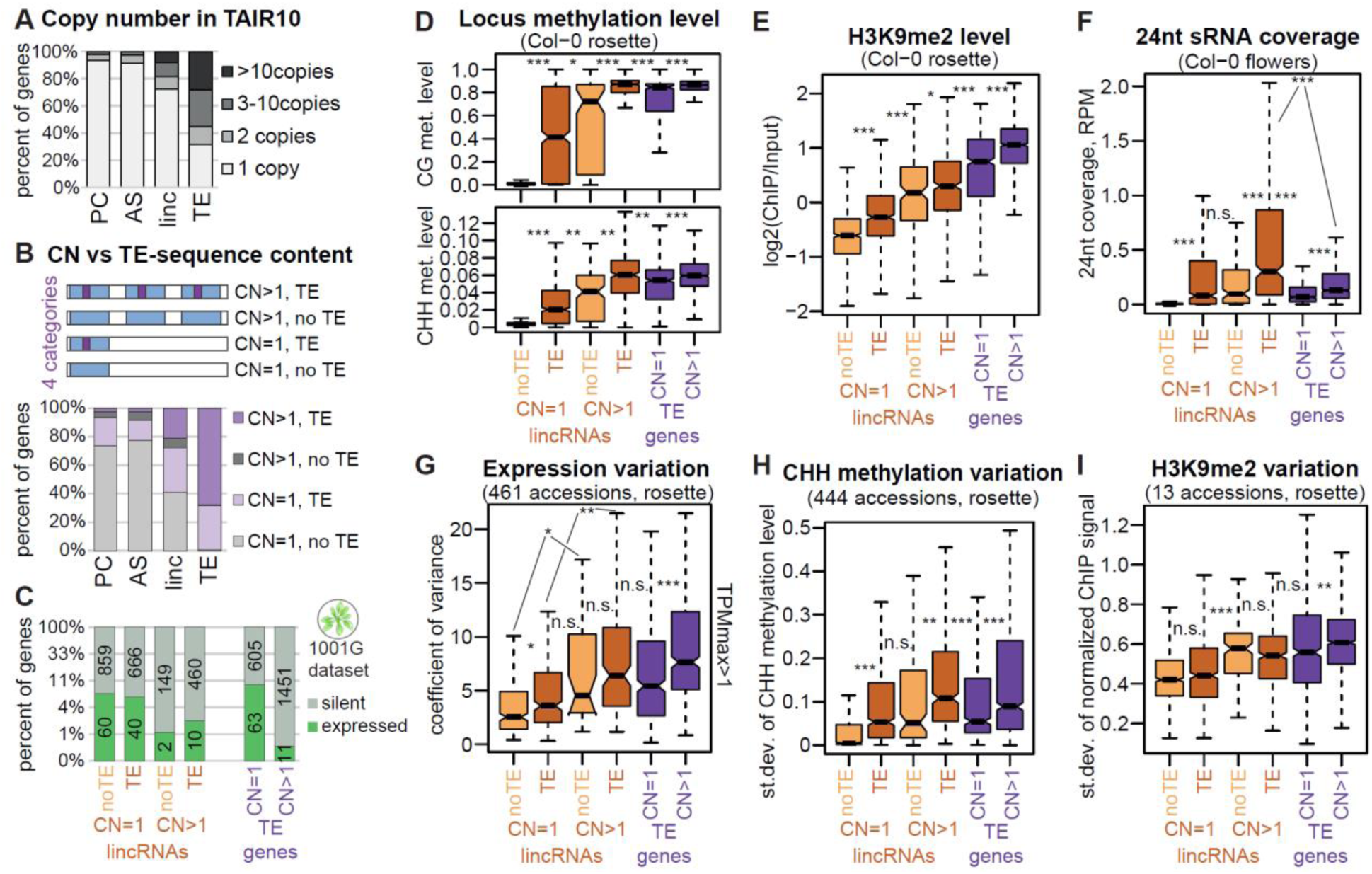
Copy number of lincRNAs affects their epigenetic patterns and variability. **A.** Distribution in copy number for PC genes, AS lncRNAs, lincRNAs and TE genes from the cumulative transcriptome annotation in the TAIR10 genome. **B.** Top, diagram of the four types of loci; bottom, the distribution of copy number of the four types of loci: 1 copy with no TE patch, 1 copy with a TE patch, multiple copies with no TE patch in the original locus, and multiple copies with a TE patch in the original locus. **C.** Proportion of the four types of lincRNAs, and two types of TE genes: expressed (TPM>0.5, green) or silent (TPM<0.5, gray) in Col-0 rosettes. The y-axis is displayed in log3 scale. **D–F.** CG and CHH methylation levels (**D**), H3K9me2 levels (**E**) in Col-0 rosettes, and 24-nt sRNA coverage in Col-0 flowers (**F**) for the four types of lincRNAs and two types of TE genes. CG and CHH methylation data displayed are from Col-0 rosettes (Kawakatsu et al., 2016). **G–I.** Boxplots showing expression (**G**), CG methylation (**H**) and H3K9me2 (**I**) variability for the four types of lincRNAs and the two types of TE genes. *P*-values in the boxplots are calculated using a Mann-Whitney test: *** *P*<10^−10^, \*\**P*<10^−5^, \**P*<0.01, n.s. *P*>0.01. Outliers in the boxplots are not plotted.

We analyzed the features of all four categories of lincRNAs and observed that increased copy number is associated with lower expression (Figure 6C) and increased repressive chromatin marks (Figure 6D-F, Supplemental Figure S27A-F), as well as expression and epigenetic variability (Figure 6G-I, Supplemental Figure S27G-I). The presence of a TE patch within multi-copy lincRNA loci was associated with strikingly increased CG and CHH methylation levels (Figure 6D) and targeting by 24-nt siRNAs (Figure 6F) but did not appear to affect the level of H3K9me2 or H1 or their variability (Figure 6E, I, Supplemental Figure S27D, I).

In summary, we show that many lincRNAs are present in multiple copies and that increased copy number is associated with increased silencing and variability in expression and epigenetic marks. This effect comes in addition to the effect of TE patches, in that lincRNAs with both multiple copies and TE patches (i.e. most TE-like) show the highest level of silencing.

### lincRNAs are silenced by TE-like and PC-like mechanisms

We saw that lincRNAs are ubiquitously silenced, with very few lincRNAs being expressed in any particular accession (Figure 2F) and with very few accessions expressing any particular lincRNA (Figure 2A). We also observed that TE pieces within lincRNAs were associated with heterochromatin and siRNA targeting (Figure 4F-H), at least when comparing lincRNA loci within a single genome. We investigated these patterns in greater detail, connecting them to known silencing pathways.

First, we observed that silent lincRNAs show a binary behavior when it comes to which silencing mark – H3K9me2 or H3K27me3 –covers the locus (Figure 7A). We observed the same pattern across all gene types, but whereas almost all TE genes showed H3K9me2 silencing and most PC genes and AS lncRNAs were covered with H3K27me3, lincRNAs were split into two large categories (Figure 7B, Supplemental Figure S28). We thus defined two non-overlapping classes of lincRNAs based on which epigenetic mark they presented: H3K9me2 lincRNAs and H3K27me3 lincRNAs, or K9 lincRNAs and K27 lincRNAs for short (Figure 7A). K27 lincRNAs were almost free of TE patches, were present as one copy, and showed low DNA methylation and targeting by 24-nt siRNAs, while K9 lincRNAs tended to have higher TE content, were present as multiple copies, and showed strikingly more DNA methylation and targeting by sRNA (Figure 7C-G). We concluded that K27 lincRNAs and K9 lincRNAs are PC-like and TE-like, respectively. The bimodality in the epigenetic features we observed before (Figure 3) can thus be explained by lincRNAs being a heterogeneous group of PC-like and TE-like lincRNAs with different features.

**Figure 7.**
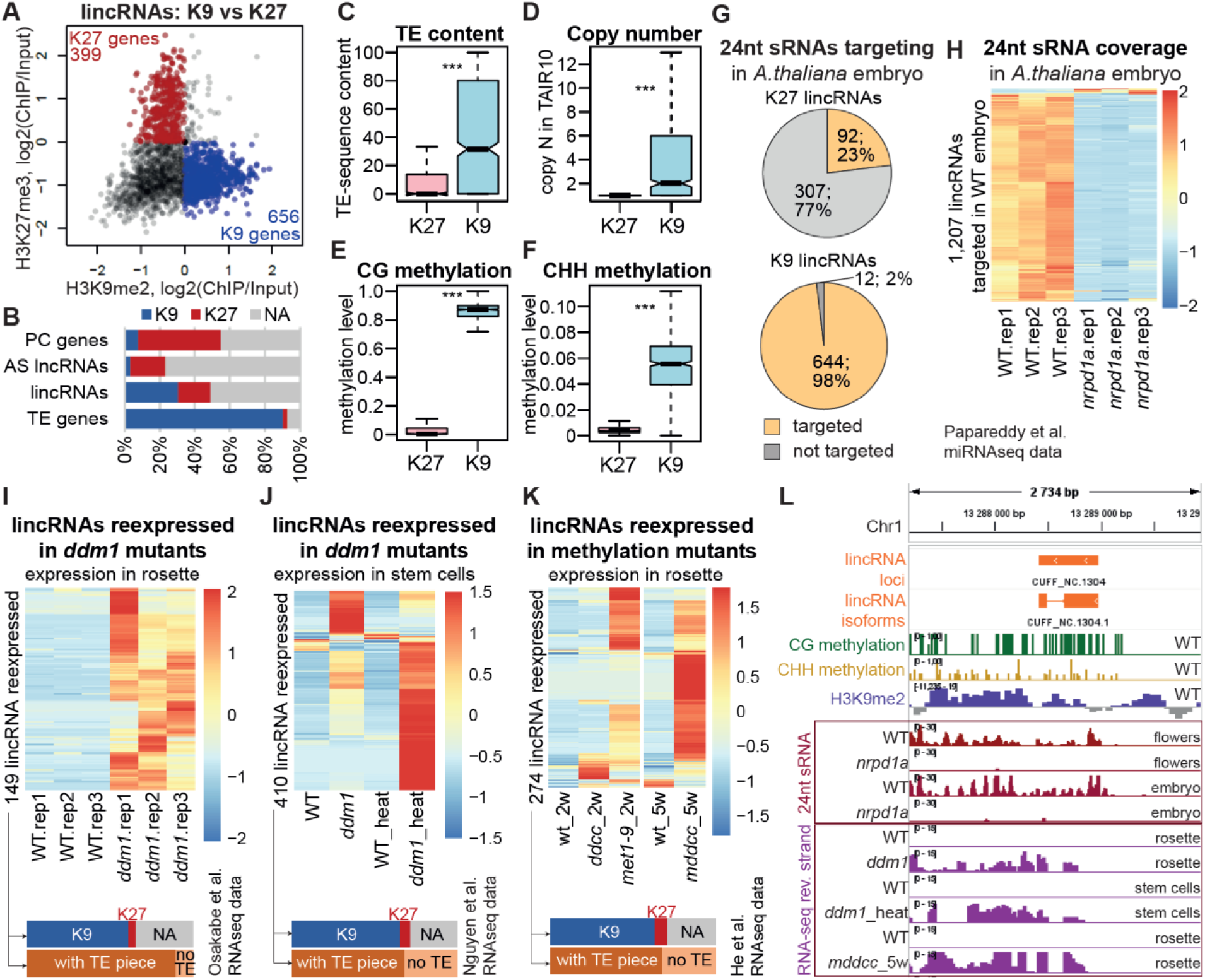
lincRNAs are silenced by PC-like and TE-like mechanisms. **A.** H3K27me3 levels as a function of H3K9me2 levels over lincRNA loci in Col-0 14-leaf rosettes (average of two replicates). K27 genes, red, K27 signal>0, K9 signal<0; K9 genes, blue, K27 signal<0, K9 signal>0. **B.** Proportion of K9 (blue) and K27 (red) genes among the four gene types. NA, genes with neither mark (gray, K27 signal<0, K9 signal<0). **C–F.** Boxplots showing the relative TE sequence content (**C**), copy number (**D**), CG methylation level (**E**) and CHH methylation level (**F**) of lincRNA loci classified as K27 or K9 genes. Outliers not plotted. *P*-values were calculated using Mann-Whitney tests: \*\*\**P*<10^−10^. CG and CHH methylation data displayed are from Col-0 rosettes (Kawakatsu et al., 2016). **G.** Distribution of K27 and K9 lincRNAs targeted by 24-nt sRNAs (reads per million [RPM]>0.03) in Arabidopsis embryos (“early heart” stage) (Papareddy et al., 2020). The sRNA coverage was averaged across three replicates. **H.** 24-nt sRNA coverage in Arabidopsis embryos (“early heart” stage) in the wild type (WT, Col-0) and in Pol IV-deficient mutants (*nrpd1a*, Col-0 background) (Papareddy et al., 2020). The 1,207 lincRNAs that are targeted (RPM>0.03, average of three replicates) by 24-nt sRNAs in the WT are plotted. **I.** Expression level of the 149 lincRNAs re-expressed in rosettes of the *ddm1* mutant in the Col-0 background (Osakabe et al., 2021). The bars at the bottom show the distribution of K9 (blue), K27 (red), and TE-containing (dark orange) or TE-free (light orange) loci among the re-expressed lincRNAs (same for **J** and **K**). **J.** Expression level of the 410 lincRNAs re-expressed in shoot stem cells of the *ddm1* mutant in the Col-0 background under mock conditions or with heat stress treatment (Nguyen et al., 2023). **K.** Expression level of lincRNAs re-expressed in the rosettes of DNA methylase mutants *met1-9*, *ddcc*, and *mddcc* (all in the Col-0 background) (He et al., 2022) (see Methods). Heatmaps were built using “pheatmap” in R with scaling by row. No column clustering, row clustering dendrograms not displayed. **L.** An example of a lincRNA epigenetically silenced in Col-0 WT but expressed in the silencing mutants.

PC-like lincRNAs are likely silenced by PRC2, which establishes the H3K27me3 repressive mark (Hansen et al., 2008); however, we did not try to confirm this hypothesis. Instead, we focused on the TE-like lincRNAs and hypothesized that the TE-like epigenetic patterns we observed were due to TE silencing pathways.

TEs in plants are thought to be silenced by two main mechanisms. First, in the RNA-directed DNA methylation mechanism known as RdDM (Onodera et al., 2005), RNA polymerase IV (Pol IV)-transcribed RNA from TE loci is turned into 24-nt sRNAs that guide the DNA methylation machinery to the locus being transcribed as well as to all homologous loci, allowing this mechanism to recognize and silence newly inserted TEs as well (Fultz et al., 2015). The second mechanism known to maintain TE silencing involves DECREASED DNA METHYLATION 1 (DDM1), METHYLTRANSFERASE 1 (MET1), CHROMOMETHYLASE 2 (CMT2), and CMT3, working together to establish the repressive H3K9me2 histone mark and DNA methylation at TE loci (Osakabe et al., 2021; Sigman and Slotkin, 2016). To test whether lincRNAs are also actively silenced by these mechanisms, we made use of publicly available RNA-seq data from mutants in components of the TE silencing machinery in Arabidopsis.

First, we analyzed the effect of inactivating the RdDM pathway. We observed above that 24-nt siRNA targeted ∼50% of all lincRNA loci in flowers. Analysis of the sRNA data from Papareddy et al (Papareddy et al., 2020) showed that 54% of lincRNA loci are targeted in early embryos; this targeting was highly specific to K9 lincRNAs (Figure 7G). Knocking out *NUCLEAR RNA POLYMERASE D1* (*NRPD1*), encoding the largest subunit of PolIV, caused a dramatic loss of 24-nt sRNA coverage over 98% of those lincRNAs in embryos (Figure 7H) as well as flowers (Supplemental Figure S29A) (Papareddy et al., 2020). As 21–22-nt sRNA were also shown to trigger RdDM-mediated TE silencing (Nuthikattu et al., 2013) and be produced by Pol IV (Panda et al., 2020; Pontier et al., 2012), we analyzed the 21–22-nt sRNA levels at lincRNA loci. We determined that, similarly to 24-nt sRNAs, increased levels of 21–22-nt sRNAs in early embryos were associated with silencing in TE-containing lincRNAs, but not in TE-free lincRNAs (Supplemental Figure S29B). Moreover, targeting by 21–22-nt sRNAs was specific to K9 lincRNAs (Supplemental Figure S29C) and knocking out *NRPD1* sharply reduced the level of 21– 22-nt small RNAs at lincRNA loci that are normally targeted in the wild type (Supplemental Figure S29 D, E).

Next, we checked for the effect of removing DDM1, a key factor in TE silencing (Osakabe et al., 2021). Using the *ddm1* mutant in the Col-0 background, we observed that 149 of our lincRNAs become re-expressed in rosette leaves in this mutant compared to Col-0 (Figure 7I), and 410 lincRNAs were re-expressed in *ddm1* stem cells (Figure 7J). Heat stress combined with knocking out *ddm1* was particularly beneficial for the reactivation of lincRNA, which is similar to TE behavior (Nguyen et al., 2023) (Supplemental Figure S30). The removal of CG and non-CG DNA methylation in Arabidopsis also allowed re-expression of many lincRNAs in rosette leaves (Figure 7K). The re-expressed lincRNAs were again predominantly K9 lincRNAs and thus mostly TE-containing (Figure 7I-K, bottom, Supplemental Table S8).

The re-expression of lincRNAs in the *nrpd1* and *ddm1* mutants underscores two important points. First, that lincRNAs, predominantly the TE-like lincRNAs, are indeed silenced by the TE-silencing machinery. Second, while we see most lincRNAs as being silent in any given accession, many retain the potential to be expressed and must therefore be actively silenced rather than having been inactivated by mutations. In fact, an analysis spanning across the different tissues and mutants with deactivated TE silencing pathways in Col-0 showed that over 50% of our annotated lincRNAs can be expressed in Col-0 (Supplemental Figure S31), in contrast to 4–10% normally expressed in one sample (Figure 2F). Thus, it appears that any genome is capable of expressing a large fraction of the numerous lincRNAs it harbors, but they are actively silenced, presumably largely via TE silencing pathways.

### TE pieces within lincRNA loci appear to attract silencing to them

The presence of TE pieces within lincRNA loci was associated with increased epigenetic silencing (Figure 5); in addition, TE silencing pathways predominantly affected lincRNAs with TE pieces (Figure 7I-K). We hypothesized that TE pieces might be decisive for TE-like silencing of lincRNAs by attracting the silencing machinery to the locus. To investigate this idea, we made use of the fact that different TE types show different silencing patterns. In particular, the RdDM pathway is more prevalent for DNA elements (Class II TEs), which are heavily targeted by 24-nt siRNAs, while retrotransposons (Class I TEs) such as LTR elements are more affected by the DDM1/CMT2 pathway showing heterochromatic patterns with high H3K9me2 levels (Sasaki et al., 2019; Sigman and Slotkin, 2016). If TE pieces inside lincRNA loci are decisive for their silencing, we would expect that lincRNAs with pieces of different types of TEs would show silencing patterns resembling that of the corresponding TEs. Our analysis confirmed this hypothesis: lincRNA loci with pieces of DNA TEs, especially those derived from *MuDR* (*Mutator Don Robertson*) elements, showed significantly increased levels of 24-nt sRNAs (Figure 8A), and lincRNAs with pieces of LTRs, especially those of *Gypsy* elements, showed significantly increased H3K9me2 levels (Figure 8B). Class I TEs are more prevalent in the chromosome arms and LTRs are enriched closer to the centromeres (Quesneville, 2020); these trends were preserved when controlling for chromosomal position (Supplemental Figure S32).

**Figure 8.**
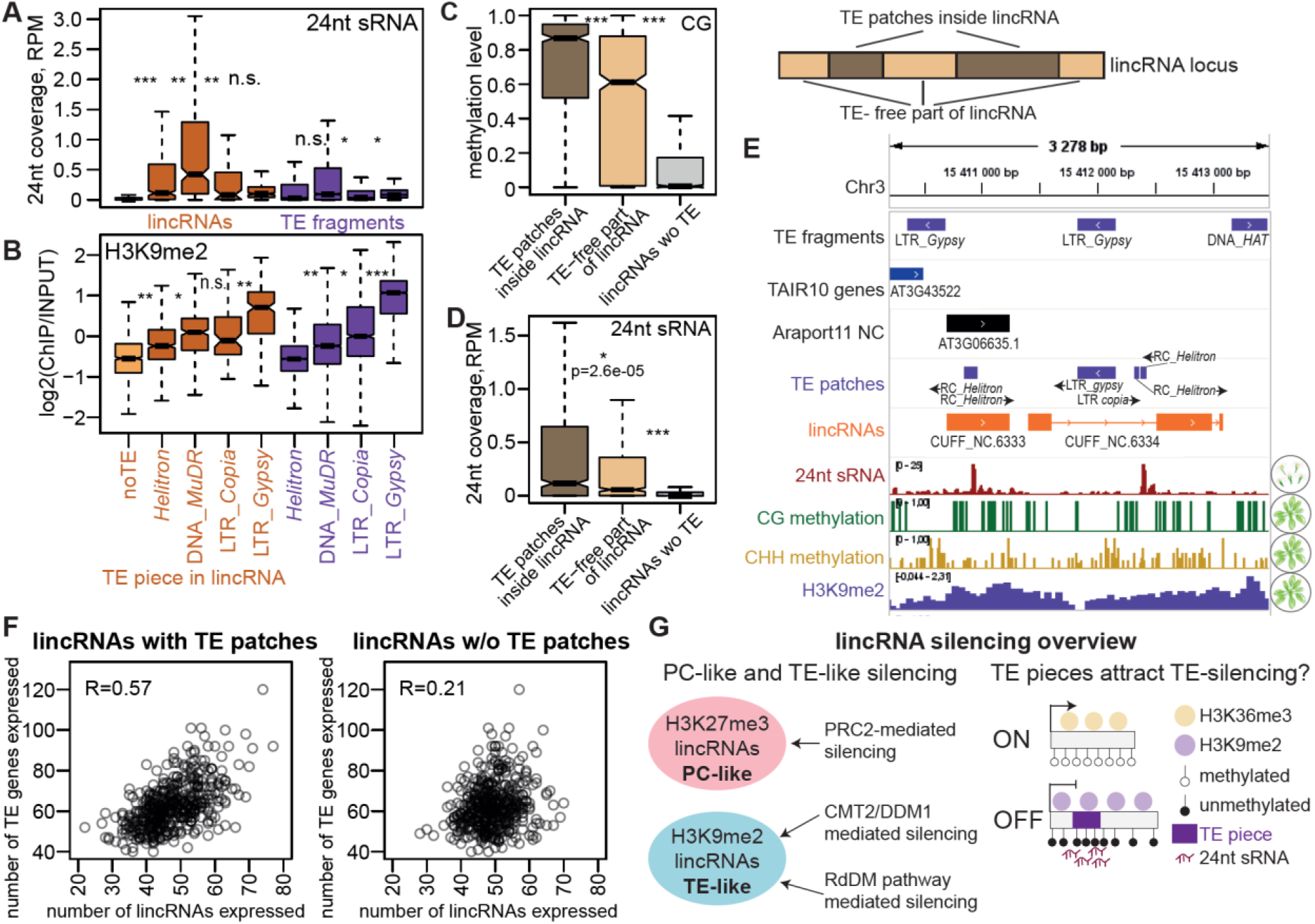
TE pieces appear to attract silencing to lincRNA loci. **A-B.** 24-nt sRNA levels in Col-0 flowers (**A**) and H3K9me2 levels in rosettes (**B**) for lincRNAs with pieces of TEs from four superfamilies and TAIR10 TE fragments from the same superfamilies. Only lincRNAs with TE pieces from one superfamily are plotted. The light-orange boxplot indicates lincRNAs without TE pieces (noTE). **C,D.** Boxplots showing CG methylation level (**C**) and 24-nt sRNA coverage (**D**) for TE patches inside lincRNAs, TE-patch-free parts of TE-containing lincRNA loci and lincRNA loci without TE patches. Outliers not plotted. *P*-values were calculated using Mann-Whitney tests: \*\*\**P*<10^−10^, \**P*< 0.01. **E.** Integrative Genomics Viewer (IGV) screenshot showing an example of lincRNAs with TE patches that have higher level of CG methylation and 24-nt sRNA coverage over TE patches than over the rest of the locus. **F.** Scatterplot showing the number of TE genes expressed in rosettes of 460 different accessions (Kawakatsu et al., 2016) as a function of the number of lincRNAs with TE pieces (left) and without TE pieces (right) expressed in the same accession. Pearson’s correlation coefficient is displayed. **G.** Summary of lincRNA silencing pathways. PC-like lincRNAs that show H3K27me3 repressive histone marks are likely silenced by PRC2, while TE-like lincRNAs that display H3K9me2 are silenced by CMT2/DDM1 and RdDM pathways. TE piece presence likely attracts TE silencing and repressive chromatin to the lincRNA locus.

Although TE patches usually constitute only a portion of a lincRNA locus (Figure 5E), they are associated with silencing over the full-length of the locus (Figure 5G). We analyzed repressive chromatin marks on the TE patch and TE-patch-free parts of lincRNA loci and determined that while TE patches show higher repressive chromatin epigenetic modification, there was a very significant increase in repressive chromatin also outside of TE patches (Figure 8C,D; Supplemental Figure S33), consistent with spreading of silencing (Sigman and Slotkin, 2016). While H3K9me2 generally covered the whole TE-containing locus, 24-nt sRNAs and DNA methylation were more restricted to the TE pieces inside loci (Figure 8C-E, Supplemental Figure S33).

Finally, we noticed that the numbers of lincRNAs and TE genes expressed in a given accession were quite well correlated (Supplemental Figure S34A). TE silencing can vary across accessions and indeed, we observed that the number of TE genes expressed across accessions varies nearly three-fold. The correlation was much stronger for lincRNAs that contained TE pieces (Figure 7N) supporting our hypothesis of shared silencing mechanisms. The number of TEs and lincRNAs expressed was correlated in every tissue (Supplemental Figure S34B), indicating the organism-wide success or failure of silencing. Interestingly, while the number of expressed loci correlated well between accessions, the correlation between the mean expression levels across expressed lincRNAs and TE genes was much lower (Supplemental Figure S34C, D) indicating that the two types of loci might share the same silencing machinery, but likely not the general transcription apparatus and factors. We tried to identify genetic factors associated with the number of TE genes and lincRNAs expressed using genome-wide association study (GWAS), but could not see any clear association, except for one nearly significant peak on chromosome 2 near the *XERICO* gene (At2g04240), encoding a protein with a zinc finger domain, (Supplemental Figure S35), which is interesting as proteins with such domains are thought to participate in TE silencing (Yang et al., 2017).

In sum, lincRNAs display two distinct silencing mechanisms (Figure 8G): PC-like silencing via H3K27me3 that is normally deposited by PRC2 (Hansen et al., 2008) and TE-like silencing, achieved via DDM1–CMT2 and RdDM silencing pathways (Fultz et al., 2015). The presence of TE pieces within lincRNAs appears to induce their TE-like silencing (Figure 8G).

## Discussion

### An extended Arabidopsis lncRNA annotation

Unlike annotations based only on the reference accession Col-0, we used almost 500 Arabidopsis accessions and several developmental stages and identified several thousand previously unannotated lncRNA loci in the TAIR 10 reference genome. We conclude that over 10% of the genome can express lncRNAs, but that most are not expressed in any particular accession or tissue, preventing a comprehensive lncRNA identification from few accessions or tissues. Analyzing more accessions allows identification of more lncRNA loci, with little evidence of saturation even when using data from several hundred accessions (Figure 1F, Supplemental Figure S5A). We provide an extended lncRNA annotation (Supplemental Data Set S1) as a resource for the Arabidopsis research community. Our results also suggest that lncRNA annotations in other plant species could similarly be extended by population-wide studies.

In our study, we annotated lncRNAs using polyA^+^ RNA-seq data. While affordable and less prone to transcriptome assembly artifacts, polyA^+^ RNA-seq can miss non-polyadenylated and/or unstable lncRNAs. Other methods, such as Global run-on sequencing (GRO-seq) (Hetzel et al., 2016), plant native elongating transcripts sequencing (plaNET-seq) (Kindgren et al., 2020), or exosome depletion (Thomas et al., 2020), can successfully detect an extended set of transcripts in Arabidopsis. Characterizing nascent and non-polyA transcription in multiple accessions and tissues would help capture the truly full scope of possible transcription and extend our understanding of transcription variation, allowing the distinction between variability in stability and in transcription initiation.

The largest part of our lncRNA transcriptome annotation consists of lncRNAs that are antisense to PC genes. Apart from the general problem of natural variation impeding lncRNA identification described above, identifying antisense lncRNAs crucially depends on having high-quality stranded RNA-seq data and a careful analysis to avoids artifacts (Supplemental Figure S1). We were able to annotate almost 9,000 antisense lncRNAs with nearly 30% of all PC genes having an antisense partner, which greatly extends the scope of antisense transcription. This is an important finding, since most functional lncRNAs reported in Arabidopsis, such as *COOLAIR* (Csorba et al., 2014), *antisense DELAY OF GERMINATION 1* (*asDOG1*) (Fedak et al., 2016), *SVALKA* (Kindgren et al., 2018), and recently *SERRATE antisense intragenic RNA* A (*SEAIRa*)(Chen et al., 2023), are antisense lncRNAs, and the massive extension of AS lncRNA annotation reported here thus opens a broad field for functional studies. A deeper investigation into antisense lncRNAs and their function is beyond this study, but we provide a list of 14 AS lncRNAs that show striking negative correlation in expression with their partner PC gene (Supplemental Table S9, Supplemental Figure S36 A, B) and thus are excellent candidates for being regulatory.

The second largest class of lncRNAs was intergenic lncRNAs that do not overlap with any PC genes, and these are the main focus of this article. This type of lncRNAs is very actively studied in mammals, with many functional examples reported (Rinn and Chang, 2020). Arabidopsis lincRNA loci we annotate in this study are enriched for previously reported interesting genetic associations (Supplemental Figure S36C) (Togninalli et al., 2020) as 157 lincRNA loci contained top GWAS hits associated with 65 different, mostly climate-related, phenotypes in Arabidopsis (Supplemental Table S10) (Togninalli et al., 2020). In this study, we focused on lincRNAs because they showed extreme expression variability and an interesting position intermediate between PC genes and TE genes in terms of expression, epigenetic features, and variation (Figures 2,3). Their bimodal distribution in CG methylation levels was particularly striking (Figure 3C). We also observed a clear dichotomy between H3K27me3 and H3K9me2 silencing (Figure 5A) that was recently also reported by Zhao et al (Zhao et al., 2022). K27-silenced and K9-silenced lincRNAs were distinct in many features, most strikingly TE piece content, which made them similar to PC genes and TE genes, respectively, thus allowing us to distinguish two lincRNA subclasses: PC-like and TE-like lincRNAs. TE-likeness was conferred by the presence of TE pieces within the lincRNA locus.

### TE pieces in lincRNAs

We showed that about half of all Arabidopsis lincRNA loci contained sequences similar to TAIR10 annotated TEs, which we refer to as TE pieces, or patches when they held similarity to more than one TE superfamily. Strikingly, TE pieces were nearly 20 times more common within lincRNA loci than within PC genes and about three times more common than in random intergenic regions (Figure 5C). It is unclear why lincRNA loci are so dramatically enriched in TE pieces. While this enrichment over PC genes is understandable, as TE insertions can be more deleterious for PC genes than for lincRNAs, the enrichment over random intergenic regions is very interesting. As lincRNAs are simply expressed intergenic regions of the genome without protein-coding capacity, the enrichment suggests that having a TE piece within the locus increases the probability of transcription. While our analyses suggest that TE pieces are associated with silencing, they might also provide the ability to be expressed when silencing fails. TEs are known to be the source of novel promoters in various organisms (Sundaram and Wysocka, 2020). Thus, we can hypothesize that for many lincRNAs, TE pieces within the locus provide the potential for being transcribed, as well as contribute to it being silenced, albeit imperfectly, leading to our ability to detect these loci in our population-wide annotation. This hypothesis would go along with the extreme expression variability of TE-containing lincRNAs (Figure 5K) and the very high variability in the overall level of TE gene and lincRNA expression (Figure 7O) that indicates high TE-silencing variability. Alternatively (and arguably more obscurely), the enrichment of TE pieces within lincRNAs may be caused by their transcriptional activity, if actively transcribed loci are more attractive for insertions compared to non-transcribed intergenic regions.

One very interesting group of TE-containing lincRNAs are lincRNAs with antisense *Gypsy* elements. LincRNAs showed significant enrichment in *Gypsy* pieces or often full elements in the antisense direction (Supplemental Figure S22C). Why *Gypsy* elements show antisense transcription more commonly than other elements remains to be investigated. We speculate that these elements might increase their mobilization chances by being transcribed from another strand, as strandedness does not matter for the transposition of retroelements, since it involves a double-stranded DNA step.

### The nature of TE pieces

Another major topic raised by our results is the nature and origin of the TE pieces we identified in lincRNA loci. Some of these TE pieces are simply parts of intact TE fragments that are overlapped by the lincRNA locus (Supplemental Figure S21E,F), and some are full TE fragments in the direction antisense to the direction of lincRNA transcription (Supplemental Figure S21C,D). In these cases, the nature of the TE sequence inside lincRNA is clear but the question of what came first — the expression or the TE — remains. Most intriguing are the many cases of short, and sometimes very short, independent pieces of TEs within lincRNA loci, the nature of which is puzzling. First, these TE pieces might represent insertions into the loci. However, their small size (Figure 5D) raises the question of how they were able to mobilize and get inserted into the lincRNA loci. Non-autonomous TEs (Quesneville, 2020), in particular, DNA-TE derived *MITE*s (miniature inverted-repeats transposable elements) (Oki et al., 2008) and LTR-TE derived *SMART*s (small LTR-retrotransposons) have been studied (Mhiri et al., 2022), yet those still have a length of a few hundred bp, while our pieces are often around 100 bp or shorter. It has also been suggested that small non-autonomous TEs can transpose with a piece of a nearby genomic sequence, thus shuffling it around, but there is little understanding of how this might work (Quesneville, 2020).

As many lincRNAs are known to originate from TEs (Kapusta et al., 2013), such as the famous *X-inactive specific transcript* (*XIST*) lncRNA (Colognori et al., 2020), it is also possible that the TE pieces we find within lincRNAs are not insertions but rather remnants of decaying TEs. One approach to distinguishing between the two possibilities would be to study the structural variation of TE pieces: variability of the presence of that precise piece would clearly indicate insertion/excision rather than the decay of a larger TE. What we could assess within the scope of this study is whether multiple TE pieces within one lincRNA locus resemble one or multiple TE families. If a locus contains pieces of different TEs, this would be evidence against the TE decay hypothesis. Among lincRNAs with more than one TE piece in TAIR10, 74% have TE pieces from different superfamilies and 24% from both Class I and Class II TEs. Further research and an analysis of full genomes from multiple accessions are crucial for understanding the nature, evolutionary history, and population dynamics of TE pieces inside lincRNAs.

### Silencing

We discovered that the Arabidopsis genome has a large potential for lincRNA expression that is massively repressed by silencing. While many lincRNAs are repressed by PC-like H3K27me3-mediated silencing, about as many are repressed by TE silencing, which is associated with having a TE piece within the locus. The presence of a TE piece was correlated with repressive chromatin marks and silencing, with higher TE contents in a locus being correlated with stronger silencing (Figure 5E-H). We also showed that inactivating TE silencing pathways in the reference accession Col-0 allowed expression of many TE-like lincRNAs that are normally completely silent in this accession. TE pieces appear to attract silencing to the locus, as we observed that lincRNAs seem to be preferentially silenced by RdDM- or CMT2-silencing pathways depending on which TE family the TE piece within the lincRNA locus came from (Figure 7E). Interestingly, TE pieces and multiple copy number were associated with the same patterns of silencing in both AS lncRNAs and PC genes (Supplemental Figure S37), although the relative number of such TE-like genes was much smaller (Figure 4B). This observation suggests that a genome-wide mechanism for suppression of TE-like loci exists (Sigman and Slotkin, 2016).

The mechanism by which short TE pieces attract TE-like silencing to a lincRNA locus is unclear. It is known that full-length TEs can induce the silencing of nearby genes by the spreading of repressive chromatin (Sigman and Slotkin, 2016) and we hypothesize that TE pieces are capable of this as well. However, how they themselves obtain repressive chromatin is unclear. One possibility is that 24-nt siRNAs produced at TE loci find the TE pieces by homology and initiate silencing at this “TE-like” locus (Fultz et al., 2015). They likely initially target only the TE piece and not the full locus, and we do see that the 24-nt siRNA and CG/CHH methylation signal is highest at TE patches (Figure 8C,D, Supplemental Figure S33). However, we also observed a significant increase of 24-nt siRNA as well as CG and CHH methylation levels outside of TE patches, which may suggest that the spreading might include sRNAs starting to be produced at the locus. It is also possible that many lincRNA loci with TE patches initiate their own silencing through the RdDM pathway, thus producing their own Pol IV-dependent sRNAs.

It is also unclear what causes the failure of silencing of certain lincRNAs in certain accessions. It is possible that the silencing machinery varies in efficiency, and we see some evidence for this in the three-fold range of variation in the number of TE genes and TE-containing lincRNAs expressed across accessions (Figure 7N). However, we could not find any gene expression level or single nucleotide polymorphisms (SNPs) that was clearly associated with the overall extent of lincRNA or TE transcription. It is also unclear how variation in silencing efficiency could account for such a strong lincRNA landscape variability across accessions with similar overall lincRNA transcription (Supplemental Figure S38). This variation may reflect the presence of particular TE loci producing the appropriate siRNAs for TE pieces within particular lincRNA loci.

Further studies are clearly needed. In this study, we focused on the reference genome, demonstrating that TE pieces within lincRNA loci are important for silencing. Direct experiments, like inserting a TE piece into a TE-free lincRNA locus and assessing the resulting expression change, are outside the scope for this study. Similarly, an analysis of the full genomes of multiple accessions, including variation for TE and TE-fragment content, would be informative, and such an analysis is underway.

### The distinction between lincRNAs and TEs

As lincRNAs with TE pieces showed many similarities to TEs, including similar epigenetic patterns, silencing pathways and increased copy number, the question might arise as to whether these TE-containing lincRNAs are distinct from TEs. By definition, a lncRNA is a transcript longer than 200 nt without protein-coding potential and thus technically any non-protein coding transcript arising from a TE can be considered a lncRNA. However, our annotation pipeline did distinguish lncRNA loci from expressed TE fragments by a <60% sense exonic overlap with annotated TE fragments, as well as by applying a protein-coding capacity cutoff to lncRNAs but not TEs. The 60% cutoff we applied is admittedly arbitrary, although common in the lncRNA field, and strongly depends on the TE annotation used. The TE annotation in Arabidopsis is far from complete and studies analyzing recent genetic mobility in Arabidopsis and annotation of new TEs are underway. A largely extended TE annotation could affect the TE-overlap filtering step we used in our annotation pipeline classifying some of our lincRNAs as “expressed TEs” (see Supplemental Figure S1). However, many of the TE patch-containing lincRNA loci showed only a minor overlap with annotated TE fragments, while quite a few (24%) had no overlap at all. TEs furthermore have a direction and many lincRNAs were transcribed antisense to the TE fragment they overlapped with, indicating that these are separate transcriptional units, even though the inherently non-strand-specific epigenetic marks were shared between the two. Moreover, most of the lincRNAs contained a mix of TE pieces from different families, which is a strong indication that these lincRNAs are unlikely to be intact TEs. Thus, we think that most of our lincRNAs are distinct from intact TEs and that the effect TE-patches have on the expression and silencing of lincRNAs, and other loci they occur in, is a very interesting phenomenon deserving future research.

Nevertheless, some of the lincRNAs we detected might actually be previously unannotated active or recently active TEs. The presence of a patch similar to annotated TEs might resemble the occasional sequence likeness between annotated TEs of different families. We identified 58 lincRNA loci with a sense TE patch that were present in more than 10 copies; they represent the most likely candidates for unannotated TEs. We also found 39 lincRNAs that, while having no TE patches, were also present in more than 10 copies, highlighting that all TE sequences are unlikely to have been annotated. However, to definitively conclude that a lincRNA locus is in fact a TE, we would need evidence of mobilization between accessions or species, and evidence of the TE piece within the lincRNA locus being an integral part of it rather than an insertion – thus not showing variability between accessions. These analyses represent future directions and are outside the scope of this study.

### lincRNA expression variation and future directions

Our study initially had two major goals: to create a population-wide map of lncRNA transcription in Arabidopsis and characterize its natural variation. We discovered that the extent of lncRNA transcription in Arabidopsis is much larger than previously thought and that lncRNA expression patterns are largely variable between accessions with half of all lncRNAs being expressed in one accession while being off in another (Figure 2E). In this study, we characterized the expression variability of lncRNAs in Arabidopsis, but we only accessed the epigenetic patterns among the factors that could explain the expression variation across accessions. We showed that lncRNAs display extensive epigenetic variation (Figure 4A,B) and this variation can explain the expression of ∼50% of informative lincRNAs and ∼20% of informative AS lncRNAs (Supplemental Figure S20B). While purely epigenetic variation is well-known (Rajpal et al., 2022; Xu et al., 2019), our analysis did not distinguish between this and when the epigenetic variation that defines expression variation is itself defined by an underlying genetic or structural variation. We showed that two structural features of lincRNA loci – their TE content and their copy number – are associated with silencing and increased expression and epigenetic variation (Figures 5, 6), and it is clear that variation in these two features might be responsible for the variation in expression that we observed between accessions. In this study, we constrained our analysis to the reference genome, because an analysis of structural variation in copy number or TE-piece-presence requires full-genome assemblies of non-reference accessions. We will perform these analyses in an upcoming study that investigates the determinants of lincRNA expression across accessions in greater depth.

In conclusion, analyzing transcriptomes from multiple accessions and tissues of Arabidopsis accessions allowed us to drastically extend its lncRNA annotation and study the natural variation of lncRNA expression. We established that 10% of the Arabidopsis genome is covered with almost 12,000 lncRNA loci, however, most of them are silent in any given sample. LncRNAs, particularly long intergenic ncRNAs, show very high expression and epigenetic variation. The silencing of lincRNAs is achieved via PC-like and TE-like mechanisms, with the latter being defined by the pieces of TEs present in about half of all lincRNAs. We produced a multi-accession transcriptome and epigenetic resource, as well as an extended lncRNA annotation useful for the Arabidopsis community and provide new insights into the genome biology and composition of lncRNAs.

## Materials and methods

### Sample collection

Seeds were surface sterilized with chlorine gas for ∼1 h, stratified at 4°C for ∼5 d to induce germination before being sown onto soil (three parts Peat moss Gramoflor professional mixture (Gramoflor GmbH) mixed with one part Gramoflor Premium Perlite 2-6 (Gramoflor GmbH)). Plants were grown in growth chambers at 21°C under long-day conditions (16-h light/ 8-h dark) with a light intensity of 130 to 150 μmol/m^2^/s (HLG-240H-30A LED lamps, Mean Well Enterprises Co., LTD). For each tissue type, all accessions were grown and processed in parallel at all stages to avoid non-accession-related variation. The “14-leaf rosette” (or mature leaves) samples were collected at the 13–16-leaf stage before plants started to bolt. Approximately 8 leaves (avoiding the oldest and the youngest leaves) were collected from 2–3 individuals of the same accession into 20ml Polyvial bottles (Zinsser Analytic) with metal beads inside, snap-frozen in liquid nitrogen, and stored at –70°C. Tissue was ground while frozen, producing 1–2 mL of tissue powder that was used for preparation of RNA-seq, bisulfate-seq, and ChIP-seq libraries. The 14-leaf rosette samples had 2–4 replicates per accession (Supplemental Table S2): accessions were grown in the growth chamber, followed by tissue harvesting two to four times, with a gap of several weeks between each batch. For each accession, the samples collected in each batch are referred to as replicates. For the “9-leaf rosette” samples, the full rosette at the 9-leaf stage was collected, with one plant harvested per sample. Seedlings were collected at 7 d post sowing (∼5 d post germination). Full seedlings with the root were harvested with approximately 10 seedlings harvested per sample. For the “flower” samples, flowers and flower buds were collected from approximately five individuals per sample. Accessions 1741, 6024, 6244, 9075, 9543, 9638, 9728, 9764, 9888, 9905, 22003, 22004, 22005, 22006, 22007 were vernalized by taking them out from the 21°C growth chamber at the age of ∼ three weeks and placing them into 10°C growth chambers (under long-day conditions) for ∼ four weeks to induce flowering. Accessions 6069 and 6124 did not flower even after the cold treatment. Pollen was collected using the method described in (Johnson-Brousseau and McCormick, 2004) that uses vacuum suction and a series of filters to harvest dry pollen from flowering plants. Polyester mesh filters of three sizes were used (150 mm, 60 mm and 10 mm) for sample collection. Collected pollen was snap-frozen in liquid nitrogen, stored at –70°C, and ground for total RNA ahead of RNA-seq using ∼0.5 mL of 0.5-mm diameter glass beads (Scientific Industries, Inc.). All tissue grinding was performed using the Retsch Oscillating Mill MM400 (Retsch GmbH) with 1/30 frequency for 90 seconds using custom made metal adapters that were pre-cooled in liquid nitrogen to keep the samples frozen while being ground.

### RNA sequencing and analysis

Total RNA was isolated and treated with DNAse I (NEB) using a KingFisher Robot with an in-house magnetic RNA isolation kit. Total extracted RNA was diluted in nuclease-free water (Ambion) and stored at –80°C. Libraries for RNA-seq were prepared using a TruSeq Stranded mRNA kit (Illumina) following the manufacturer’s protocol with a 4-min RNA fragmentation time and 12 PCR cycles for library amplification. RNA-seq was performed at the Vienna Bio Center (VBC) NGS facility on an Illumina HiSeq 2500 machine in paired-end read mode of 150 bp and 125 bp. Raw RNA-seq data were aligned to the TAIR10 genome using STAR (Dobin et al., 2013) with the following options: --alignIntronMax 6000 --alignMatesGapMax 6000 --outFilterIntronMotifs RemoveNoncanonical --outFilterMismatchNoverReadLmax 0.1 --outFilterMismatchNoverLmax 0.3 --outFilterMultimapNmax 10 --alignSJoverhangMin 8 --outSAMattributes NH HI AS nM NM MD jM jI XS. Gene expression levels were calculated using featurecounts from the Subread package with -t exon option and an exonic SAF file as an annotation (Liao et al., 2014).

### Transcriptome assembly and lncRNA annotation

Transcriptome assembly was performed in several steps as described in Supplemental Figure S1; the scripts are provided at https://github.com/aleksandrakornienko/Kornienko_et_al_lncRNA_expression_variation_and_silencing. In brief, the following RNA-seq datasets were used for transcriptome assembly: 14-leaf-rosette data from 461 accessions (100-bp single-end reads) (Kawakatsu et al., 2016), seedling data from Cortijo et al (Cortijo et al., 2019) (75-bp paired-end, 14 replicates for each of the 12 samples were pooled), our 14-leaf-rosette data from 28 accessions (2–4 replicates each) (125-bp paired-end), our seedling and 9-leaf-rosette data from 27 accessions (150-bp paired-end), and our flower and pollen data from 25 accessions (150-bp paired-end). First, the transcriptomes of each sample were assembled separately using Stringtie v.2.1.5 (Pertea et al., 2015) with options: -c 2 -m 150 -j 2.5 -a 15 guided by the TAIR10 gene annotation (-G). The transcriptome assemblies of the same tissue and data types were then merged using Cuffmerge (Cufflinks v.2.2.1) (Trapnell et al., 2010) with --min-isoform-fraction 0 before performing a second merging of the resulting seven transcriptomes to obtain the cumulative transcriptome annotation. A series of filtering steps were applied, including a transcript-length cutoff of 200 nt for multiexon genes and 400 nt for single-exon genes, and then the genes were split into 1) PC genes by exonic overlap with TAIR10 or Araport11 annotated PC genes; 2) TE genes based on exonic overlap with Araport11 annotated TE genes; 3) TE fragments with >60% same strand exonic overlap with TE fragments annotated in Araport11; 4) Pseudogenes by exonic overlap with Araport11 annotated pseudogenes; 5) Initial lncRNAs showing no overlap with PC genes, TE genes, or pseudogenes, and with <60% same strand exonic overlap with TE fragments annotated in Araport11. Then lncRNA transcripts (and the corresponding loci containing those transcripts) with protein-coding capacity as tested by CPC2 (Kang et al., 2017) were removed; rRNA, tRNA, sn/snoRNA and miRNA precursor lncRNAs were classified based on overlap with the appropriate annotations. The remaining lncRNAs were classified into 1) Antisense lncRNAs by antisense overlap with TAIR10 or Araport11-annotated PC genes; 2) lncRNAs antisense to pseudogenes; 3) lncRNAs antisense to Araport11 TE genes (AS_to_TE); 4) intergenic lncRNAs (lincRNAs) with no overlap with PC genes, TE genes or pseudogenes. LincRNAs were additionally filtered against loci that started <100 bp downstream from annotated genes to avoid read-through transcripts. The number of Araport11 PC genes with an antisense transcript was calculated using Araport11 non-coding and novel_transcribed_region annotations filtered for genes longer than 200 bp.

### Gene saturation curve

To create the gene saturation curves for accession and tissue number, the annotation pipeline was automated and run many times with different numbers of accessions and tissues. The accession saturation curve was generated by inputting 10 to 460 transcriptome assemblies (one assembly being one accession) obtained from the 1001 Arabidopsis genome dataset (Kawakatsu et al., 2016) into the same annotation pipeline used for the main gene annotation defined above. Subsampling of accessions was done randomly with eight iterations for each number of accessions. The curve fitting and prediction of the saturation curve behavior with up to 1000 accessions was done by fitting a linear model using the lm function in R with the command line: model <– lm(y ∼ x + I(log2(x))) (Supplemental Figure S5A,B). The control for the accession saturation curve was done using the data from Cortijo et al. (Cortijo et al., 2019), from which 1 to 12 transcriptome assemblies (corresponding to 12 samples with 14 replicates per sample pooled into one BAM file pre-assembly) were randomly chosen and fed into our standard annotation pipeline, counting the number of loci identified as an output. The procedure was performed eight times for each assembly number. Then the number of reads was calculated and juxtaposed to the number of reads in the multi-accession saturation curve (Supplemental Figure S5C). As different datasets had different read modes, the results were aligned by calculating the total read length and multiplying it by the total read number. The tissue saturation curve analysis was performed on 23 accessions that had data from all four tissues. Random sampling of accessions was performed with eight iterations as replicates for each number of accessions. Tissues were assessed in this particular order: 1) seedling; 2) rosette; 3) flowers; 4) pollen, without random sampling (Supplemental Figure S6).

### Expression variation analysis

Inter-accession variability was calculated as coefficient of variance (standard deviation divided by mean) of TPM of the locus across accessions in the dataset. When multiple replicates were available, the average between the replicates was taken for the coefficient of variation calculation. Intra-accession variability was calculated using our 14-leaf multi-replicate dataset as follows: for each accession, the coefficient of variance for TPMs across replicates was calculated, and then the coefficients of variance for each accession were averaged. Expression noise was calculated using the seedling dataset from (Cortijo et al., 2019) that contained 14 replicates for 12 time points: coefficient of variance across 14 replicates was calculated for each of the 12 time points and then these 12 coefficients of variance were averaged to produce the resulting “noise” level. Circadian expression variation was also calculated using the seedling dataset from Cortijo et al: for each time point TPMs from 14 replicates were averaged and then coefficient of variance was calculated across the 12 time points. The 12 time points represent the samples collected every 2 hours within a 24-hour period of time (12 hours light, 12 hours dark) and by “circadian expression variation” we mean - expression variation during a 24-hour period (across the 12 time points).

### ChIP sequencing

Chromatin immunoprecipitation was performed with a protocol adapted from (Yelagandula et al., 2014) (full protocol is available at https://github.com/aleksandrakornienko/Kornienko_et_al_lncRNA_expression_variation_and_silencing). Briefly, 1–2 g of ground frozen leaf tissue was fixed with 1% formaldehyde at 4°C for 5 min, then nuclei were isolated and lysed using a series of lysis and centrifugation steps; chromatin was fragmented in 1 ml Covaris milliTUBEs using a Covaris E220 Focused-ultrasonicator for 15 min at 4 °C with the following settings: duty factor of 5.0, peak incident power of 140; 200 cycles per burst. An aliquot was then taken out as input sample and frozen at –20°C; the remaining supernatant was split in five tubes, one for each antibody, and were processed together. The antibodies used were against: H1 (Agrisera) - 3 micrograms per reaction, H3K4me3 (Abcam), 3 µg per reaction; H3K9me2 (Abcam), 4 µg per reaction; H3K27me3 (Millipore), 4 µg per reaction; H3K36me3 (Abcam), 4 µg per reaction. The immunoprecipitation was performed using pre-washed Dynabeads Protein A magnetic beads (Invitrogen) and incubated at 4°C overnight. Afterwards the samples were washed, followed by elution, overnight reverse-crosslinking, treated with RNAse A (Fermentas) for 30 min at room temperature, and the DNA isolated using a Qiaquick PCR purification kit (Qiagen) with 0.3 M sodium acetate. Next, ChIP-seq libraries were prepared from half of the resulting sample (due to very low amounts, we did not measure the DNA concentrations) with a NEBNext Ultra II DNA kit (New England Biolabs) according to the manufacturer’s protocol and sequenced as 100-bp single-end reads on an Illumina NovaSeq 6000 instrument.

### ChIP-seq analysis

Raw ChIP-seq reads were mapped using STAR (Dobin et al., 2013) adjusted for ChIP-seq with the following options --alignIntronMax 5 --outFilterMismatchNmax 10 --outFilterMultimapNmax 1 --alignEndsType EndToEnd. Only samples with > 1 million unique non-duplicated reads were used for analysis. Aligned BAM files from each ChIP-sample were then normalized by the corresponding input samples using bamCompare from deeptools (Ramírez et al., 2016) with the following options: --operation log2 --ignoreDuplicates --effectiveGenomeSize 119481543; bigwig and bedgraph files were created. Read coverage over loci and promoters was estimated using bedtools map on the bedgraph files with the “mean” operation. To estimate the variation in histone modification levels, the ChIP-seq coverage values were normalized again to achieve the same range of values across accessions, applying quantile normalization, setting the 20% and 80% quantile values for each sample to the same value across samples with the function in R: quantile_minmax <– function(x) {(x-quantile(x,.20)) / (quantile(x,.80) - quantile(x,.20))}. Histone modification variation was then calculated as the standard deviation of quantile-normalized levels averaged across replicates for each accession.

### Bisulfite sequencing

Bisulfite sequencing was performed as described in (Pisupati et al., 2022). Briefly, DNA was extracted from frozen leaf tissue (14-leaf rosettes) using a Nuclear Mag Plant kit (Machery-Nagel) and the bisulfate sequencing libraries were prepared using a tagmentation method described in (Wang et al., 2013) using an in-house Tn5 transposase (IMBA-IMP-GMI Molecular Biology Services) and an EZ-96 DNA Methylation-Gold Mag Prep kit (Zymo Research) for bisulfite conversion. Bisulfate sequencing libraries were sequenced on thane Illumina NovaSeq 6000 instrument in the 100-bp paired-end mode.

### DNA methylation analysis

Bisulfite sequencing data were used to call methylation in three contexts (CG, CHG and CHH where H stands for A, C or T) using the method described in (Pisupati et al., 2022). The methylation level per locus for each context was determined by dividing the number of methylated reads by the total number of reads covering the cytosines in the CG, CHG or CHH context. Thus, the values of methylation of each locus range from 0 to 1 and roughly correspond to the ratio of methylated to total cytosines in the locus (we did not take the average of the ratios for each cytosine to avoid high error rates caused by low read coverage).

### Small RNA sequencing and analysis

Small RNA was isolated from frozen and ground flower samples using a NucleoSpin miRNA kit (Macherey-Nagel) following the manufacturer’s protocol. Small RNA-seq libraries were prepared using a QIAseq miRNA Library Kit (Qiagen). Raw fastq files were trimmed with cutadapt (v.1.18) (Martin, 2011) using -a AACTGTAGGCACCATCAAT and --minimum-length 18 options. Trimmed reads were aligned to the TAIR10 genome using STAR (v.2.9.6) adjusted for sRNA-seq, allowing 10 multimappers and 2 mismatches. Reads that were 24-nt-long, and likewise 21– 22nt-long reads, were extracted and read coverage for each bp of the genome was calculated using genomeCoverageBed (bedtools v.2.27.1) and normalized by dividing by the unique number of reads in the sample. Final sRNA coverage was calculated by mapping the normalized read coverage per bp over the loci of interest and calculating the average coverage across all bp of each locus. Raw sRNA-seq data from (Papareddy et al., 2020) were processed using the same pipeline. The cutoff for calling a locus as being targeted by 24-nt or 21–22-nt sRNAs was set to RPM=0.03.

### Explaining expression variation by DNA methylation variation levels

To determine if DNA methylation can explain expression variation, matching RNA-seq and DNA methylation data from the 1001 Genomes project was used (Kawakatsu et al., 2016). Out of 461 RNA-seq samples, 444 had matching bisulfite sequencing data, thus data for these 444 accessions were used for this analysis. For each lncRNA in our annotation, we asked if accessions where a lncRNA was highly methylated showed significantly lower expression (Mann-Whitney test, *P*<0.01) than accessions with low methylation levels at this lncRNA locus (Supplemental Figure S20A-B). This analysis was performed for exonic gene-body TPM calculated as described above using four estimates of methylation level: CG and CHH methylation level of gene body; CG and CHH methylation level of promoters (TSS ± 200 bp). We set “low” CG methylation level as “<0.5”, and “high” CG as “≥0.5”, and “low” CHH methylation level as “<0.01”, and “high” CHH as “≥0.01”. These cutoffs were defined based on the distribution of CG and CHH methylation levels of PC genes (average gene body methylation level across 444 accessions): 90% of PC genes in our transcriptome annotation had accession-wide-mean CG methylation levels below 0.37 and CHH methylation levels below 0.01.

### TE-piece analysis

To find TE pieces in various loci, 31,189 annotated TE sequences from TAIR10 were compared to the sequence of each locus using BLASTN (BLAST+ v2.8.1) with options -word_size 10 - strand both -evalue 1e-7. We required >80% sequence identity and did not restrict the length. We then merged same-strand overlapping TE pieces into TE patches. For all of our analyses we grouped TE families into 7 superfamilies: DNA_other, DNA_MuDR, SINE_LINE, RC_Helitron, LTR_Gypsy, LTR_Copia, Unassigned_NA.

### Copy number analysis

Copy number was estimated by extracting the sequence of the locus from the TAIR10 genome and using it as a query against the TAIR10 genome using BLASTN (BLAST+ v2.8.1) with options - word_size 10 -strand both -outfmt 7 -evalue 1e^−7^. We allowed for copies to be disrupted by insertions of no more than 1.5 kb and applied a cutoff of >80% on sequence identity and >80% on length to all regions identified by BLASTN.

### lincRNA re-expression analysis

Raw RNA-seq data from the silencing mutants from (Osakabe et al., 2021; He et al., 2022; Nguyen et al., 2023) were processed using the same RNA-seq pipeline as described above. The re-expression in the mutants was generally defined as the lack of expression in the wild type (TPM<0.5, averaged from all available replicates), the presence of expression in the mutant (TPM>0.5), and additionally a three-fold difference between the expression in the mutant and the wild type (WT) (MUT>3*WT). For the *ddm1* knockout in stem cells, the mock *ddm1* mutant sample was matched with the mock WT and the heat-treated *ddm1* mutant was matched with the heat-treated WT sample (heat treatment as described in (Nguyen et al., 2023)). For the methylation mutants, *ddcc* and *met1-9* mutants were 2-week-old seedlings and matched with the 2-week-old WT control, while the *mddcc* mutant was matched ith the 5-week-old WT control. The *met1-9* mutant corresponds is a *MET1* knockout with loss of CG methylation; the *ddcc* mutant is the quadruple mutant for *DOMAINS REARRANGED METHYLASE 1* (*DRM1*), *DRM2*, *CMT2*, and *CMT3* with a loss of CHG and CHH methylation; the *mddcc* mutant is a quintuple mutant for MET1, *DRM1*, *DRM2*, *CMT2*, and *CMT3* with a nearly full loss of all methylation (He et al., 2022).

### Use of public datasets

The summary of the public datasets used in our study and the corresponding mapping statistics are available in Supplemental Tables S2, S3 and S11. The public datasets were downloaded from NCBI GEO using the specified GEO accession numbers below:

1. RNA-seq and bisulfite-seq from mature leaves of 14-leaf rosettes from the 1001 Genomes project (Kawakatsu et al., 2016): **GSE80744** and **GSE43857**.
2. RNA-seq data from Col-0 seedlings from 12 time points with 14 technical replicates each (Cortijo et al., 2019): GEO accession number **GSE115583.**
3. Early embryo and flower bud sRNA-seq data from *nrpd1* knockouts (Papareddy et al., 2020): **GSE152971**.
4. Rosette RNA-seq data from *ddm1* knockouts (Osakabe et al., 2021): **GSE150436**.
5. Stem cell RNA-seq data from *ddm1* knockouts with and without heat stress (Nguyen et al., 2023): **GSE223915**
6. Rosette RNA-seq data from DNA methylation-free mutants (He et al., 2022): **GSE169497**.

### Availability of data and materials

The sequencing data produced in this study are available at the NCBI Gene Expression Omnibus (https://www.ncbi.nlm.nih.gov/geo/) as SuperSeries GSE224761. The gene annotations and the 14-leaf rosette RNA-seq dataset from 28 Arabidopsis accessions are available under the accession number GSE224760 (note that gene annotations are also available in the supplement as Supplemental Data Set S1); the corresponding bisulfite sequencing data are under the GEO accession number GSE226560. The 14-leaf rosette ChIP-seq data from 14 accessions are available under accession number GSE226682. The RNA-seq data from seedlings, 9-leaf rosettes, flowers, and pollen from 25 to 27 accessions are available under the GEO accession number GSE226691. The flower sRNA-seq data from 14 Arabidopsis accessions are available under the GEO accession number GSE224571. The code used for the analyses as well as the full ChIP protocol are available at our GitHub repository (https://github.com/aleksandrakornienko/Kornienko_et_al_lncRNA_expression_variation_and_silencing).

## Supporting information

Gene annotations

## Acknowledgements

We would like to thank the Vienna Biocenter Core Facilities GmbH (VBCF) Next Generation Sequencing for NGS services, VBCF Plant Sciences for the excellent growth chambers, IMBA-IMP-GMI Molecular Biology Services for providing the access to instruments, molecular biology reagents and support, and the VBC Ethics, Health and Safety team for their support during the COVID pandemic. We would like to thank Mirjam Bissmeier for help with preliminary ChIP-seq analyses, and Bhagyshree Jamge, Vu Nguyen, Ramesh Yelagandula, Zdravko Lorkovic, Nathalie Durut, and Ortrun Mittelsten Scheid for their advice with experiments and data, and fruitful discussions. We would like to greatly thank Detlef Weigel for helping to secure funding for the “1001 Genomes Plus” project and for his helpful comments on the manuscript.

## Author contributions

A.E.K. and M.N. designed the study. A.E.K. performed most of the experiments and data analyses. A.M.M. helped with sample collection. R.P. performed the bisulfite sequencing data processing. V.N. prepared all libraries for sequencing. A.E.K. and M.N. wrote the paper. The authors declare they have no competing interests.

## Supplemental files

Supplemental Figure S1. Cumulative transcriptome annotation pipeline.

Supplemental Figure S2. Transcriptome annotation supplement.

Supplemental Figure S3. Antisense lncRNAs supplement.

Supplemental Figure S4. Genomic distribution of annotated loci.

Supplemental Figure S5. Gene identification saturation analysis.

Supplemental Figure S6. Gene identification saturation analysis: tissues and accessions.

Supplemental Figure S7. Expression frequency.

Supplemental Figure S8. Inter-accession expression variation in different tissues.

Supplemental Figure S9. Expression variability controls: absolute expression level and gene length.

Supplemental Figure S10. Inter- and intra-accession expression variability with replicates.

Supplemental Figure S11. Histone mark profiling.

Supplemental Figure S12. DNA methylation levels supplement.

Supplemental Figure S13. DNA methylation vs distance to the centromere.

Supplemental Figure S14. Heterochromatic histone marks vs distance to the centromere.

Supplemental Figure S15. Histone modifications of silent and expressed genes: supplement.

Supplemental Figure S16. DNA methylation of silent and expressed genes: supplement.

Supplemental Figure S17. Coverage of 24nt and 21-22 sRNA in early embryo and leaves.

Supplemental Figure S18. Epigenetic patterns in non-reference accessions.

Supplemental Figure S19. Epigenetic variation supplement 1.

Supplemental Figure S20. Epigenetic variation supplement 2.

Supplemental Figure S21. TE patches in lincRNAs.

Supplemental Figure S22. Sense and antisense TE pieces in lincRNAs: content, family, position.

Supplemental Figure S23. Expression variation vs TE content: expression control.

Supplemental Figure S24. lincRNA methylation variation vs. TE content: supplement.

Supplemental Figure S25. TE pieces affect epigenetics when controlled for chromosomal location.

Supplemental Figure S26. Copy number supplement.

Supplemental Figure S27. Copy number affects lincRNA epigenetic pattern: supplement.

Supplemental Figure S28. H3K27me3 and H3K9me2 dichotomy.

Supplemental Figure S29. Loss of 24-nt targeting in *nrpd1a* mutants.

Supplemental Figure S30. TE silencing mutants supplement.

Supplemental Figure S31. The scope of lincRNA expression potential in Col-0.

Supplemental Figure S32. Genomic position and epigenetics of lincRNAs with pieces of Class I and II TEs.

Supplemental Figure S33. Spreading of silencing from TE patches.

Supplemental Figure S34. Variability of the number of TE genes and lincRNAs expressed.

Supplemental Figure S35. GWAS on expressed TE gene number.

Supplemental Figure S36. lncRNA candidates.

Supplemental Figure S37. TE pieces affect expression and epigenetic patterns of AS lncRNAs and PC genes.

Supplemental Figure S38. High expression variability despite similar expressed gene numbers.

Supplemental Table S1. Accessions overview.

Supplemental Table S2. RNA-seq data overview.

Supplemental Table S3. RNA-seq data statistics.

Supplemental Table S4. Overview of transcriptome assemblies.

Supplemental Table S5. ChIP-seq summary and statistics.

Supplemental Table S6. Summary of bisulfite-seq samples and read number.

Supplemental Table S7. Small RNA-seq summary and statistics.

Supplemental Table S8. LincRNAs re-expressed in TE silencing mutants.

Supplemental Table S9. AS lncRNA candidates with anticorrelated expression from their partner PC gene.

Supplemental Table S10. lincRNAs with AraGWAS hits.

Supplemental Table S11. Public RNA-seq datasets summary and statistics.

Supplemental Data Set S1. Compiled annotations for all types of loci produced in this study.

Supplemental Data Set S2. Summary of statistical analyses.

