## Supplemental Figures for "Population-level annotation of lncRNA transcription in Arabidopsis reveals extensive variation associated with transposable element-like silencing"

### Table of Contents

Supplemental Figure S1. Cumulative transcriptome annotation pipeline ... 2
Supplemental Figure S2. Transcriptome annotation supplement ... 4
Supplemental Figure S3. Antisense lncRNAs supplement ... 5
Supplemental Figure S4. Genomic distribution of annotated loci ... 6
Supplemental Figure S5. Gene identification saturation analysis ... 7
Supplemental Figure S6. Gene identification saturation analysis: tissues and accessions ... 8
Supplemental Figure S7. Expression frequency ... 9
Supplemental Figure S8. Inter-accession expression variation in different tissues ... 10
Supplemental Figure S9. Expression variability controls: absolute expression level and gene length ... 11
Supplemental Figure S10. Inter- and intra-accession expression variability with replicates ... 13
Supplemental Figure S11. Histone mark profiling ... 14
Supplemental Figure S12. DNA methylation levels supplement ... 15
Supplemental Figure S13. DNA methylation vs distance to the centromere ... 16
Supplemental Figure S14. Heterochromatic histone marks vs distance to the centromere ... 17
Supplemental Figure S15. Histone modifications of silent and expressed genes: supplement ... 18
Supplemental Figure S16. DNA methylation of silent and expressed genes: supplement ... 19
Supplemental Figure S17. Coverage of 24-nt and 21–22-nt sRNA in early embryo and leaves ... 20
Supplemental Figure S18. Epigenetic patterns in non-reference accessions ... 21
Supplemental Figure S19. Epigenetic variation supplement 1 ... 22
Supplemental Figure S20. Epigenetic variation supplement 2 ... 23
Supplemental Figure S21. TE patches in lincRNAs ... 24
Supplemental Figure S22. Sense and antisense TE pieces in lincRNAs: content, family, position ... 26
Supplemental Figure S23. Expression variation vs TE content: expression control ... 27
Supplemental Figure S24. lincRNA methylation variation vs. TE content: supplement ... 28
Supplemental Figure S25. TE pieces affect epigenetics when controlled for chromosomal location ... 29
Supplemental Figure S26. Copy number supplement ... 30
Supplemental Figure S27. Copy number affects lincRNA epigenetic pattern: supplement ... 31
Supplemental Figure S28. H3K27me3 and H3K9me2 dichotomy ... 32
Supplemental Figure S29. Loss of sRNA targeting *nrpd1a* mutants ... 33
Supplemental Figure S30. TE silencing mutants supplement ... 34
Supplemental Figure S31. The scope of lincRNA expression potential in Col-0 ... 35
Supplemental Figure S32. Genomic position and epigenetic pattern of lincRNAs with pieces of Class I and II TEs ... 36
Supplemental Figure S33. Spreading of silencing from TE patches ... 37
Supplemental Figure S34. Variability of the number of TE genes and lincRNAs expressed ... 38
Supplemental Figure S35. GWAS on expressed TE gene number ... 39
Supplemental Figure S36. lncRNA candidates ... 40
Supplemental Figure S37. TE pieces affect expression as epigenetic patterns of AS lncRNAs and PC genes ... 41
Supplemental Figure S38. High expression variability despite similar expressed gene numbers ... 42
References ... 43

**Supplemental Figure S1** (Supports Figure 1)

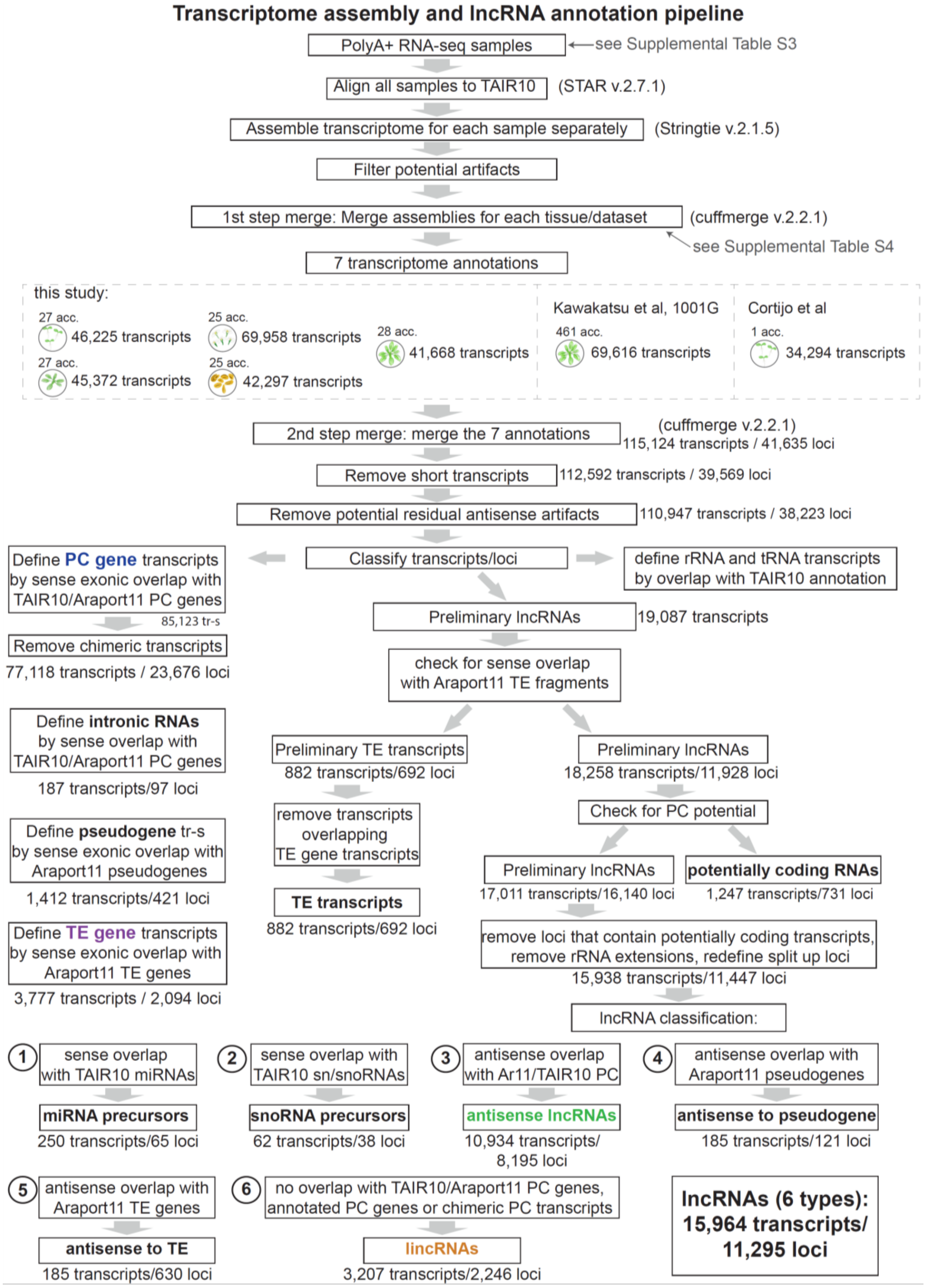


### Supplemental Figure S1. Cumulative transcriptome annotation pipeline

Algorithm for de novo lncRNA and mRNA identification, showing the number of transcripts and loci identified at different steps. First, we used polyA$^+$ RNA-seq data (Supplemental Table S3) to create

625 transcriptome assemblies using Stringtie with options: -A $abundance_file -p 4 -c 2 -m 150 -j 2.5 -a 15 -x chloroplast,mitochondria --rf G $ref_annotation. We then filtered each transcriptome assembly using these consecutive steps:

**1.** Remove all transcripts shorter than 200 nt,

**2.** Remove all transcripts with TPM < 0.5,

**3.** Remove all single-exon transcripts shorter than 400 nt with TPM<5.0,

**4.** Remove potential "mirror" transcripts (suspected strand-leakage): transcripts with >30% exonic overlap of the transcript on the opposite strand must be well-expressed above our strand-leakage cut-off: TPM_antisense_transcript must be > 0.06+ 0.012*TPM_sense_transcript (model obtained by preliminary testing).

**5.** Remove chimeric transcripts with >30% same-strand overlap with two Araport11 PC genes.

Pre-filtered transcriptome assemblies from separate samples were merged using Cuffmerge (Cufflinks v. 2.2.1 (Trapnell et al. 2012)) with --min-isoform-fraction 0 option. Merging was performed in 2 steps. First, we merged assemblies from similar tissues and read-modes (Supplemental Table S4):

**1.** seedlings (28 samples from 27 accessions),

**2.** 9-leaf rosettes (28 samples from 27 accessions),

**3.** leaves from 14-leaf rosettes (28 samples from 28 accessions),

**4.** pollen (28 samples from 25 accessions),

**5.** flowers (40 samples from 25 accessions),

**6.** 14-leaf rosettes from the 1001 Genomes (Kawakatsu et al. 2016) (461 samples from 461 accessions),

**7.** seedlings from (Cortijo et al. 2019) (12 samples from one accession).

We thus obtained seven transcriptome assemblies, which we then merged using Cuffmerge again. After the second step of merging (115,124 transcripts: 41,635 nonoverlapping loci), additional filtering against potential artefacts was performed.

**1.** Remove transcripts shorter than 200 nt and single-exon transcripts shorter than 400 nt (retained 112,592 transcripts: 39,569 non-overlapping loci).

**2.** Remove potential residual antisense leakage artefacts:

**a.** transcripts that are single-exon, overlap an exon of a multiexon transcript with >98% of their length) and are shorter than 600 nt,

**b.** transcripts that have very similar introns (>95% antisense-strand overlap) on the antisense strand to PC genes. (Retained 110,947 transcripts: 38,223 non-overlapping loci)

Then PC genes, pseudogenes and TE genes were identified by sense exonic overlap with Araport11 or TAIR10 annotated respective gene categories. PC genes underwent an additional chimeric transcript removal step. Preliminary lncRNAs were defined by no overlap with Araport11/TAIR10-annotated PC genes, TE genes and pseudogenes, no exonic overlap with non-chimera-filtered de novo annotated PC genes (to avoid calling fragments of possible PC gene extensions as lncRNAs), and no more than 60% sense exonic overlap with Araport11 annotated TEs.

The preliminary lncRNA transcripts were then filtered by calculating their protein-coding potential using CPC2 (Kang et al. 2017). All lncRNA transcripts that were part of the locus that was defined as containing TE gene, PC gene or pseudogene transcripts were removed. Then lncRNA transcripts were classified into different categories:

**1.** miRNA precursors by sense overlap with TAIR10-annotated miRNAs,

**2.** sno/snRNA precursors by sense overlap with TAIR10-annotated sno/snRNAs,

**3.** Antisense lncRNAs by antisense overlap with Araport11/TAIR10-annotated PC genes (not *de novo* annotated PC genes).

**4.** lncRNAs antisense to pseudogenes by antisense overlap with Araport11-annotated pseudogenes,

**5.** lncRNAs antisense to TE genes (AS_to_TE) by antisense overlap with Araport11 annotated TE genes (not *de novo* annotated TE genes),

**6.** lincRNAs by no overlap with Araport11/TAIR10 annotated PC, de novo PC genes (not filtered for chimeric transcripts), pseudogenes. Additionally, all lincRNAs starting closer than 100 bp downstream from PC genes or pseudogenes were removed to avoid calling fragments of read-through transcripts as lncRNAs.

**Supplemental Figure S2** (Supports Figure 1)

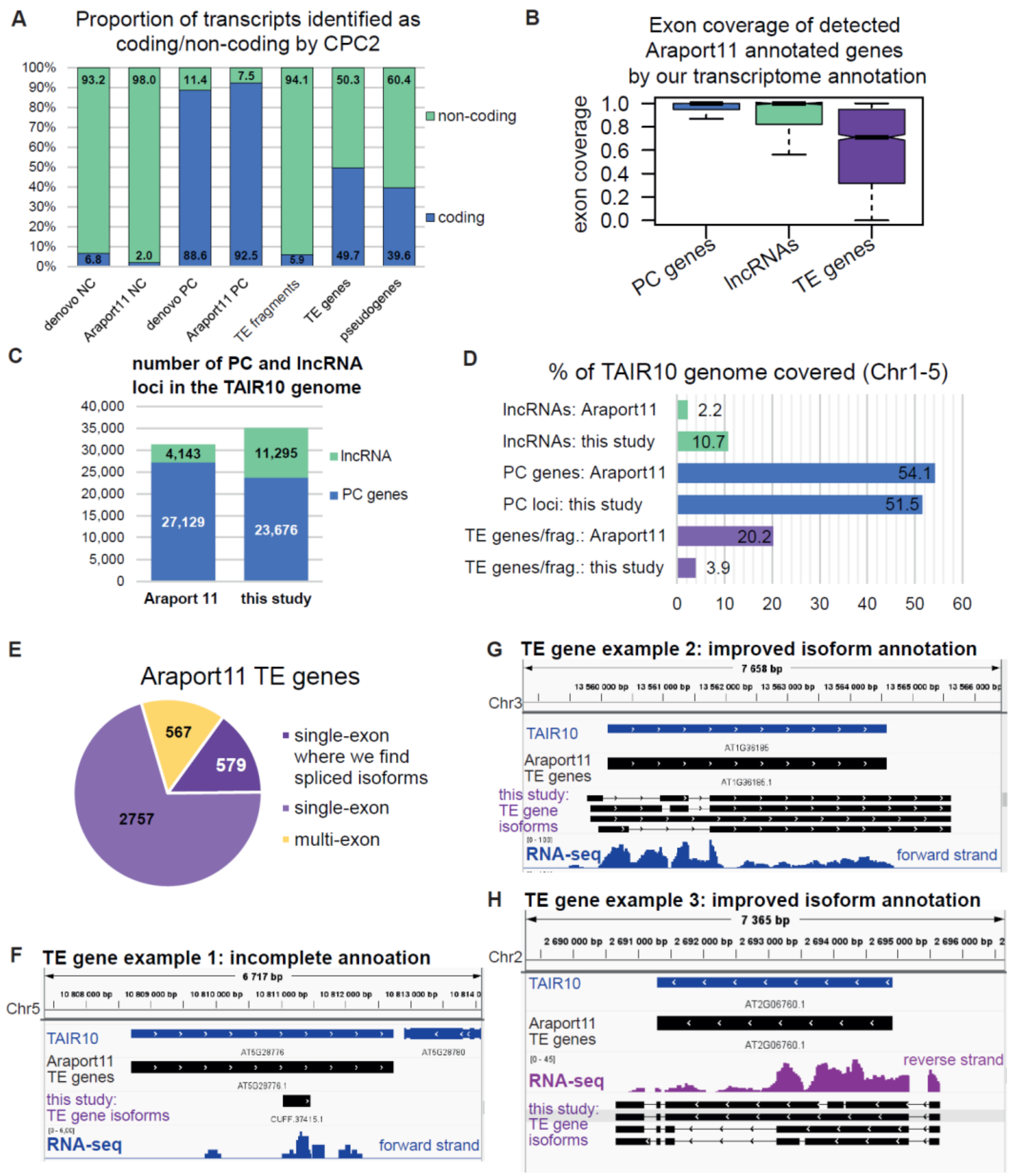


### Supplemental Figure S2. Transcriptome annotation supplement

**A.** Transcripts of different gene types identified as “coding” or “non-coding” by CPC2 (Kang et al. 2017). **B**. Same-strand exonic coverage of Araport11-annotated genes by our transcriptome annotation. Only genes that were detected are displayed. Araport11-annotated genes were filtered by length to match the length-filtering within our annotation pipeline: only genes with transcripts longer than 200 nt; single-exon genes must be longer than 400 nt. Araport11 lncRNA annotation was combined from non-coding RNA and novel transcribed regions annotation. Outliers are not plotted. The low coverage of TE genes can be explained by many incompletely annotated TE genes (as shown in **F**) and many TE genes with improved spliced annotation (as shown in **G** and **H**) that reduces the exonic coverage of the single exon annotated by Araport11 by introducing introns. **C.** Number of PC and lncRNA loci. **D.** Genome coverage. **E.** Distribution of Araport11-annotated TE genes. We defined “single-exon genes where we find spliced isoforms” as Araport11 TE genes with 1 exon for which our transcriptome annotation (Figure 1) finds multi-exon isoforms that covered > 50% of the gene length. **F.** An example of an Araport11 TE gene for which we produce an incomplete annotation, likely because of its low expression in all samples. **G, H.** Example of Araport11-annotated “single-exon” TE genes for which we improved the annotation finding extended and spliced isoforms.

**Supplemental Figure S3** (Supports Figure 1)

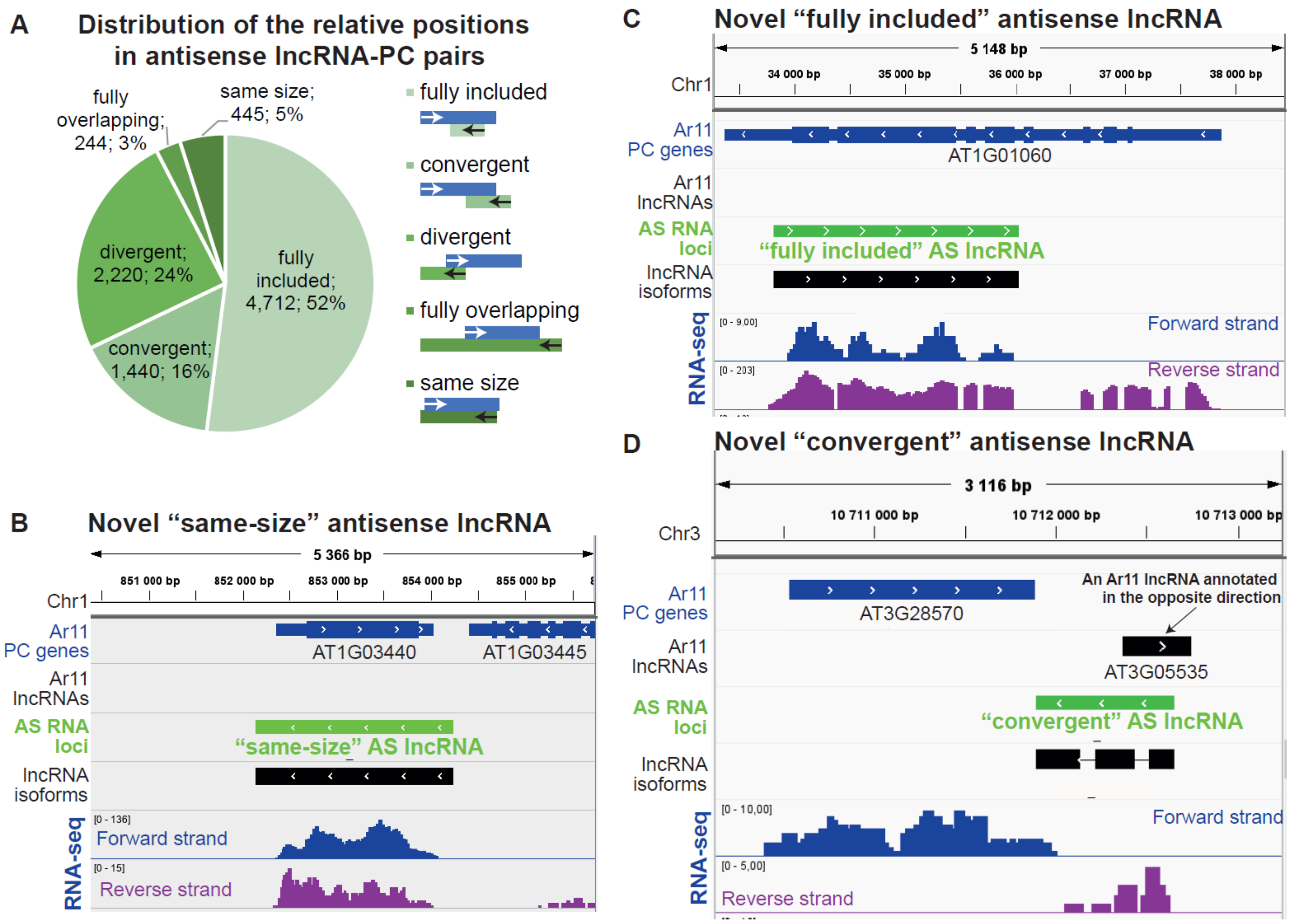


### Supplemental Figure S3. Antisense lncRNAs supplement

**A.** Distribution of AS lncRNA types in the cumulative transcriptome annotation. In the legend on the right, blue represents PC genes and green represents AS lncRNAs. **B.** An example of a previously unannotated "same-size" AS lncRNA. **C.** An example of a previously unannotated "fully-included" AS lncRNA. **D.** An example of a previously unannotated "convergent" AS lncRNA

**Supplemental Figure S4** (Supports Figure 1)

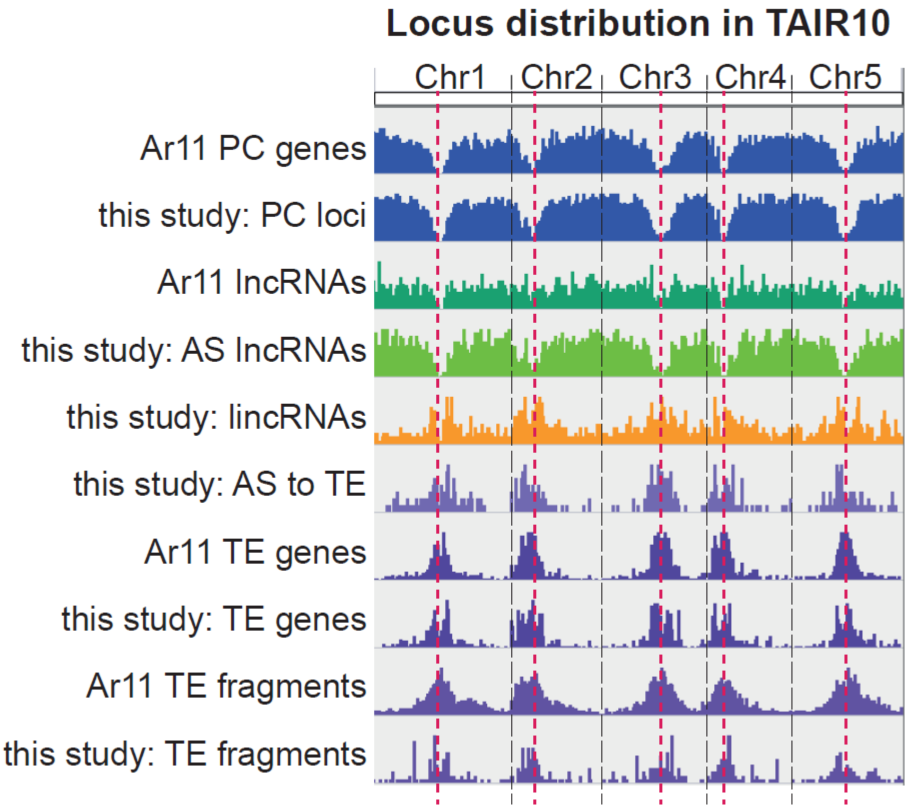


### Supplemental Figure S4. Genomic distribution of annotated loci

Genomic distribution of different gene types. The plot shows the IGV browser view of gene annotations on the 5 *A. thaliana* chromosomes. Ar11: Araport 11 annotations. Dashed vertical red lines indicate centromeres.

### **Supplemental Figure S5** (Supports Figure 1)

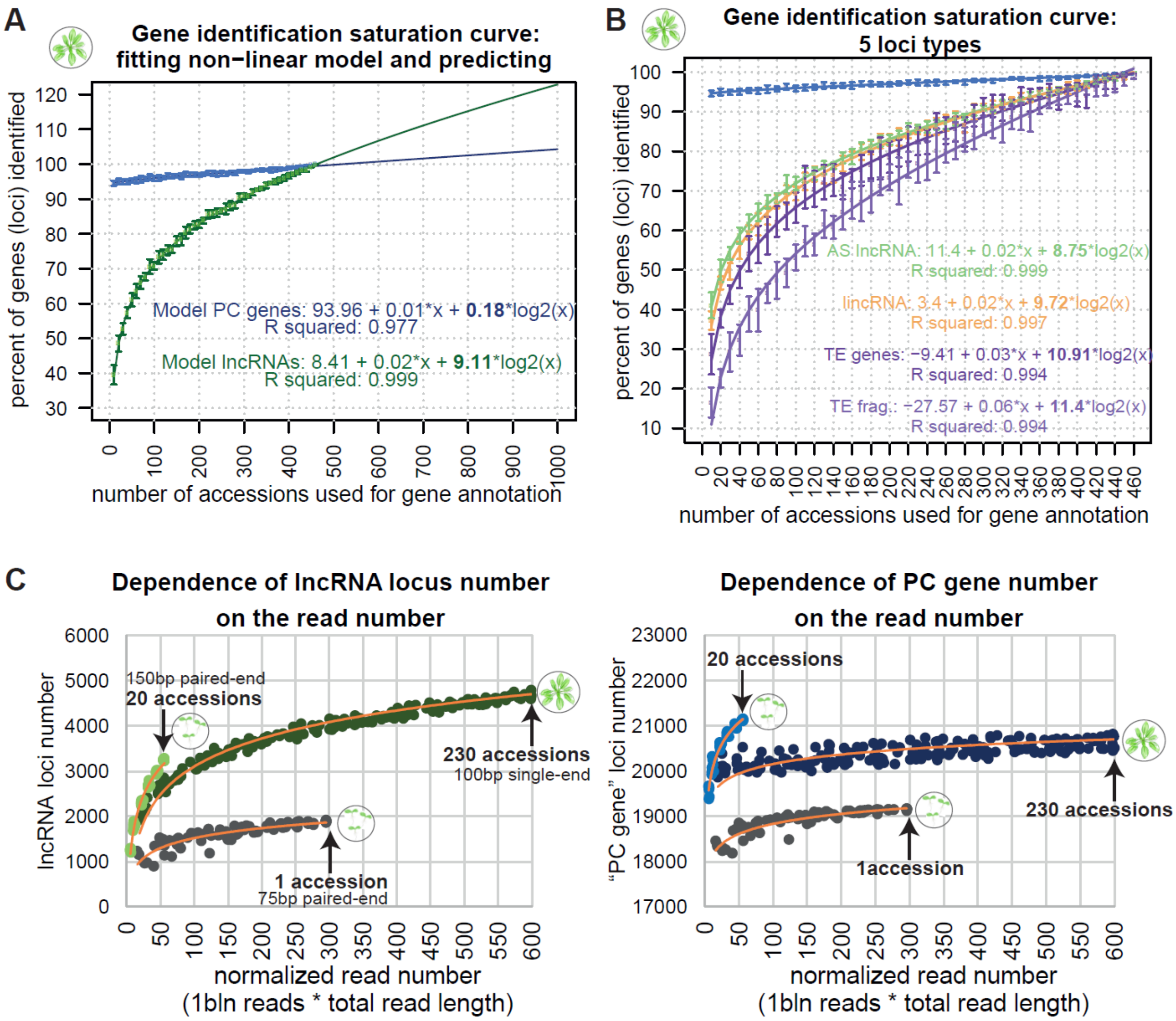


### **Supplemental Figure S5. Gene identification saturation analysis**

**A.** Data displayed in Figure 1F with fitted linear models for all lncRNAs in green (six lncRNA types as shown in Figure 1D) and PC genes in blue. The curve fitting and prediction of the saturation curve behavior with up to 1000 accessions was done by fitting a linear model using the lm function in R: model <– lm(y ~ x + I(log2(x))). **B.** Data displayed in Figure 1F with fitted linear models for AS lncRNAs (green), lincRNAs (orange), TE genes (purple) and TE fragments (light purple). Model fitting as in (**A**). **C.** Controlling for the role of the increase of the number of total reads used for the gene identification by comparing the gene identification dynamics from multi-accession datasets vs. a large single-accession dataset from (Cortijo et al. 2019). Plots show number of lncRNA loci (left) and PC gene loci (right) identified using our annotation pipeline. The gene numbers from the seedling data from Cortijo et al are displayed as dark-grey circles. The gene numbers from the rosette dataset of the 1001 Genomes Project with up to 460 accessions (as shown in (A)) is shown with dark green/blue circles and the plot is cropped at 230 accessions. The gene numbers from our seedling dataset with up to 20 accessions (as shown in Supplemental Figure S6) is shown with light green/blue circles. As every dataset had different read modes, we could not directly compare the read number between the datasets, so we normalized the read number of each annotation (one annotation is one data point on the plot) by multiplying the total read length and the total read number used for transcriptome assemblies. For paired-end reads total read number was double the read length.

**Supplemental Figure S6** (Supports Figure 1)

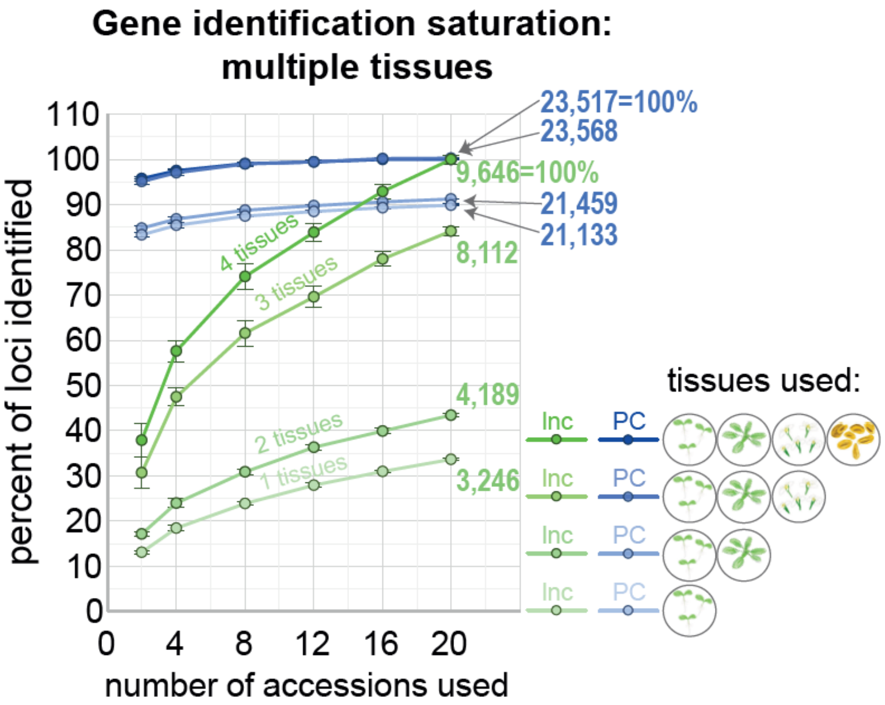


**Supplemental Figure S6. Gene identification saturation analysis: tissues and accessions**

Relative number of lncRNA (shades of green) and PC gene (shades of blue) loci (y-axis) identified using the annotation pipeline when using a varying number of accessions (x-axis) and tissues (from paler to brighter: 1, 2, 3 and 4 tissues as displayed on the bottom right of the plot). The lines and points corresponding to the PC gene numbers from 3 or 4 tissues are almost indistinguishable. The error bars represent standard deviation between the 8 replicates of random accession-picking for each accession number. The 100% is set to the loci number annotated using 20 accessions and 4 tissues. The numbers of the right show the absolute number of loci identified using 20 accessions. Tissues used: 7-day-old seedlings, 9-leaf rosettes, flowers, and flower buds, pollen.

**Supplemental Figure S7** (Supports Figure 2)

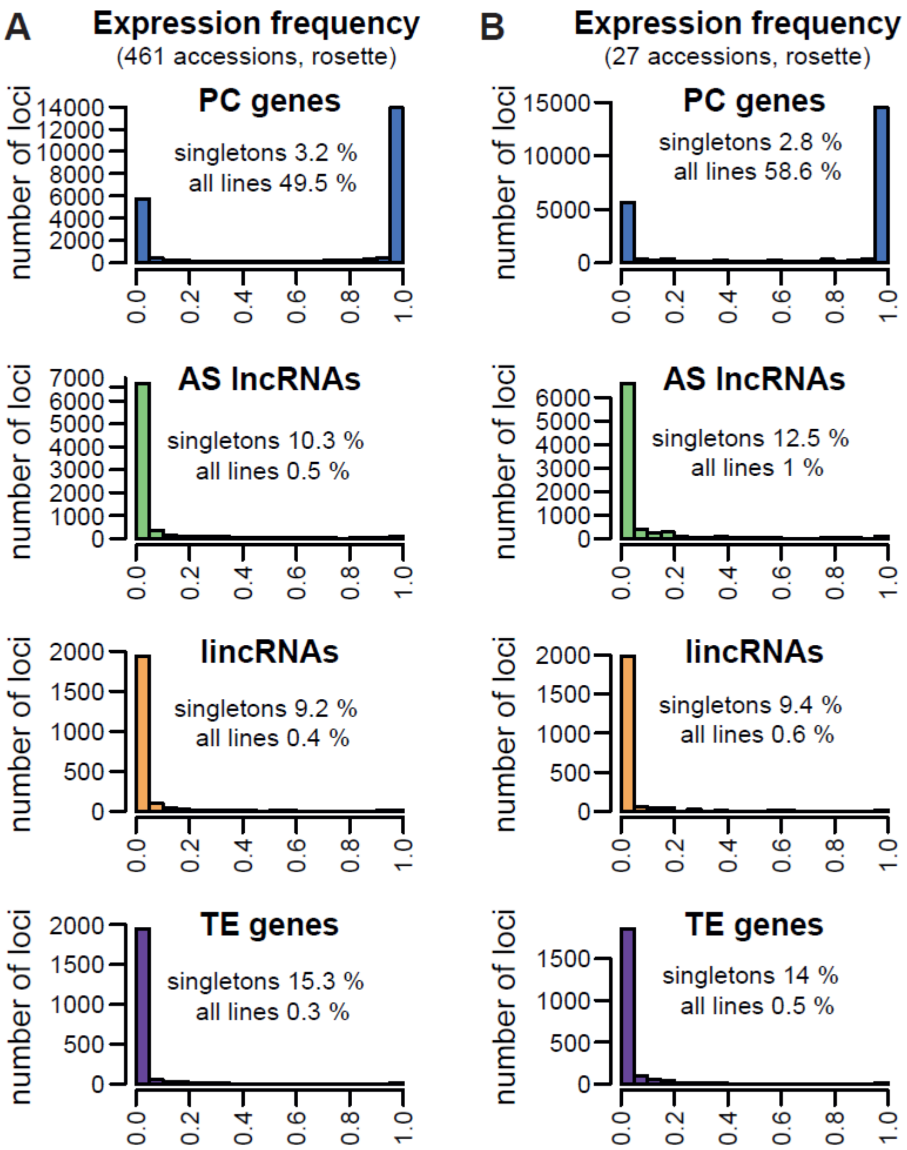


**Supplemental Figure S7. Expression frequency**

Frequency of the expression of the four types of loci across (**A**) 461 accessions ((Kawakatsu et al. 2016), rosette) and (**B**) 27 accessions (this study, 9-leaf rosette). The histograms show in how many accessions the locus is expressed (TPM>0.5), as a proportion of total accessions in each sample. The percentages of loci that are only expressed in one accession ("singletons") and that are expressed in every accession ("all lines") are listed on each histogram.

**Supplemental Figure S8** (Supports Figure 2)

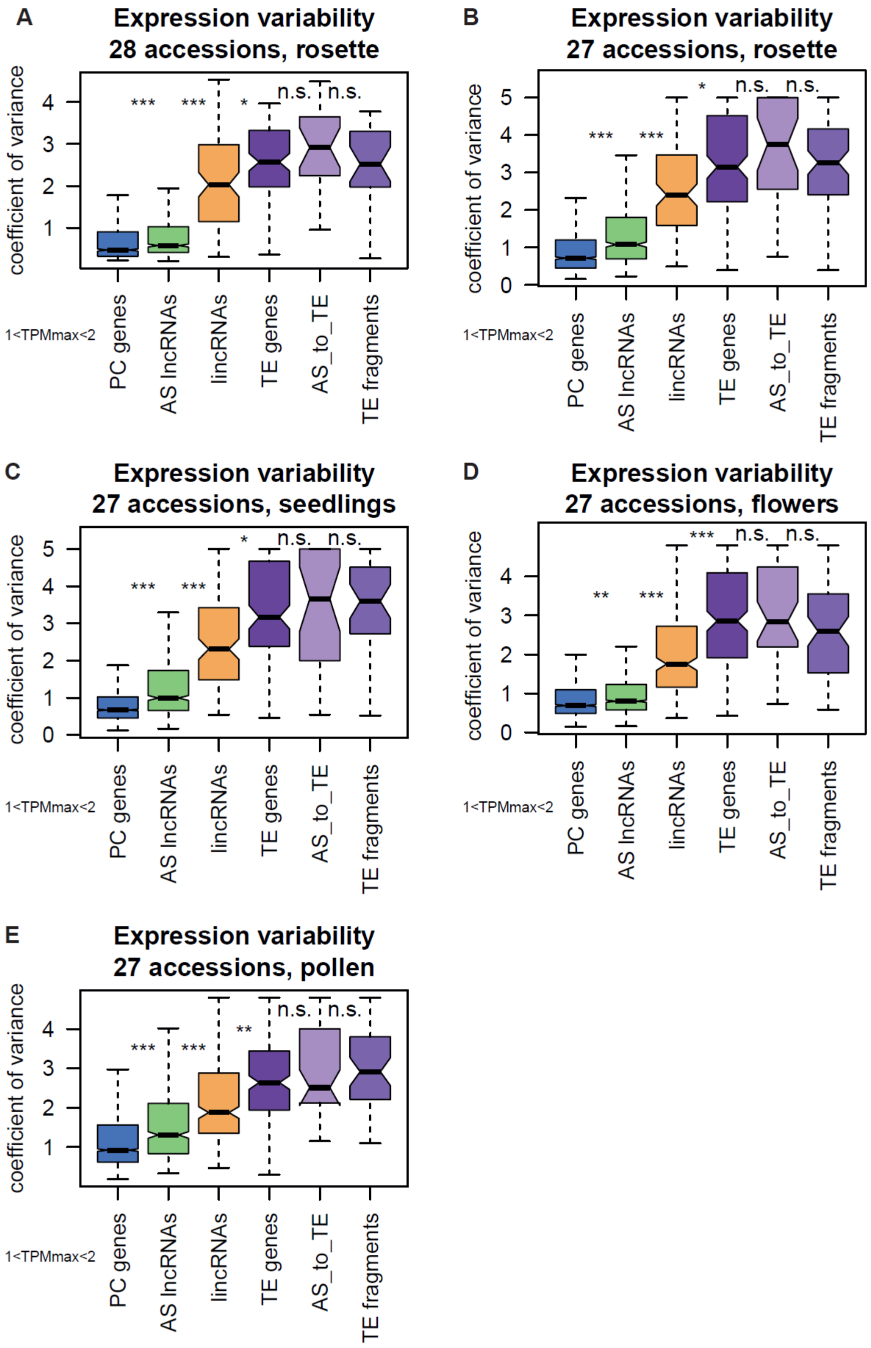


### Supplemental Figure S8. Inter-accession expression variation in different tissues

Expression variability across natural accessions of different types of loci in 14-leaf-rosette (**A**), 9-leaf rosette (**B**), seedlings (**C**), flowers (**D**) and pollen (**E**). Only loci for which their maximal expression (TPM) across the analyzed accessions is in the range from 1 to 2 are plotted. Outliers are not plotted. *P*-values were calculated using Mann-Whitney tests on equalized sample sizes: *** $P < 10^{-10}$, ** $P < 10^{-5}$, * $P < 0.01$, n.s.: $P > 0.01$.

**Supplemental Figure S9** (Supports Figure 2)

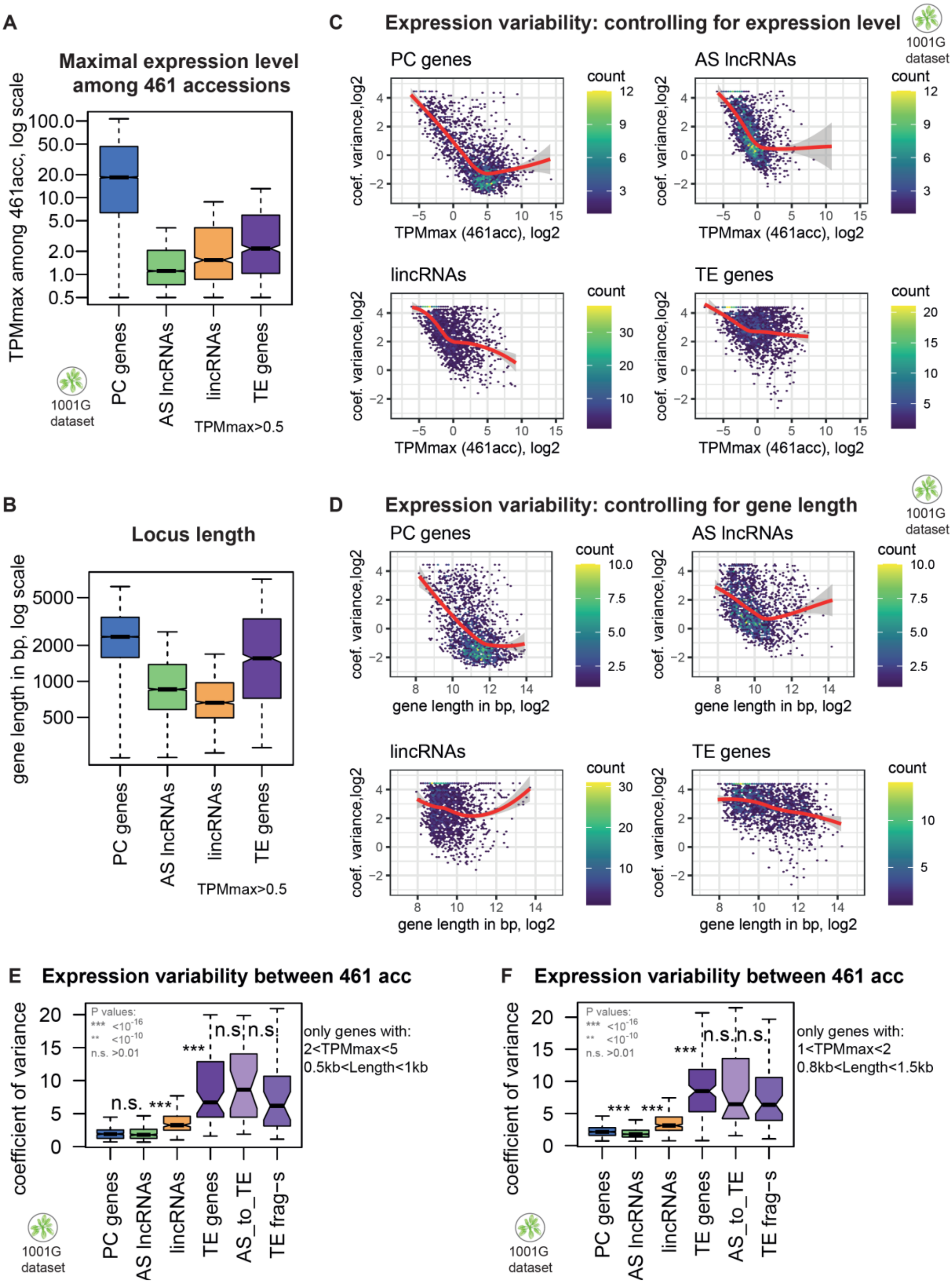


**Supplemental Figure S9. Expression variability controls: absolute expression level and gene length**

**A.** Maximal expression levels of PC genes, AS lncRNAs, lincRNAs and TE genes in rosettes across 461 accessions (Kawakatsu et al. 2016). Only genes that are expressed (TPM>0.5) in at least one accession are plotted. **B.** Length of PC, AS lncRNA, lincRNA and TE loci in our transcriptome annotation. **C-D.** Coefficient of variance as a function of the maximal expression level across accessions (**C**) and locus length (**D**). Data from 461 accessions, rosettes, (Kawakatsu et al. 2016). Both x- and y-axes values are log2. The scatterplot was built in R using geom_hex(bins = 70) + scale_fill_continuous(type = "viridis"). The trendlines were fitted using geom_smooth(method='loess', formula= y~x,col="red") in ggplot. **E-F.** Expression variability of different types of loci across 461 accessions, rosette (Kawakatsu et al. 2016), for genes with expression and length conditions specified on the right from the boxplot. Outliers are not plotted. *P*-values were calculated using Mann-Whitney tests on equalized sample sizes: *** $P < 10^{-10}$, ** $P < 10^{-5}$, * $P < 0.01$, n.s.: $P > 0.01$.

**Supplemental Figure S10** (Supports Figure 2)

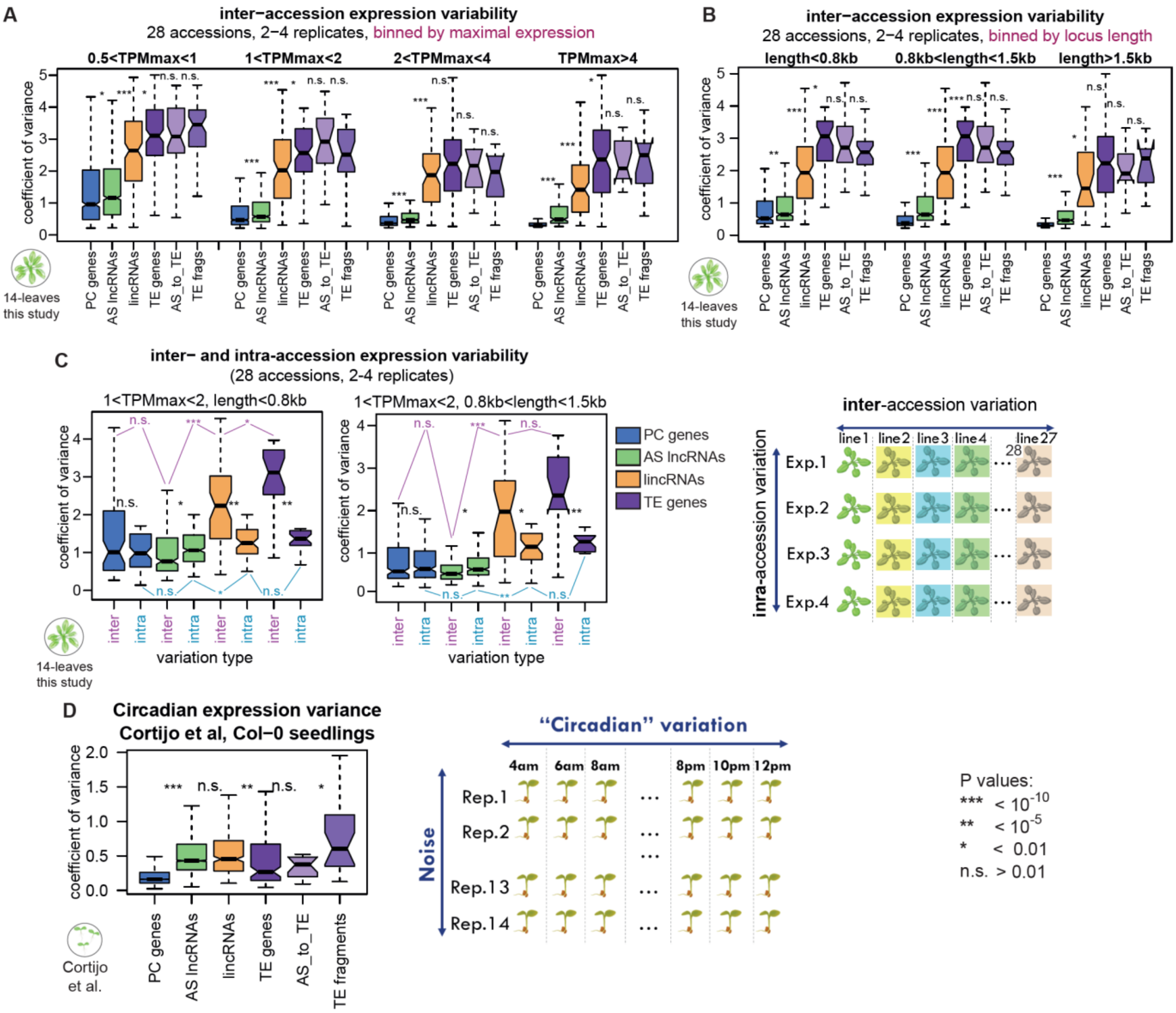


### Supplemental Figure S10. Inter- and intra-accession expression variability with replicates

**A. Inter**-accession expression variability for genes binned according to their maximal TPM across accessions (replicated RNA-seq from 28 accessions from 14-leaf rosettes). **B. Inter**-accession expression variability for genes binned according to their length (replicated RNA-seq from 28 accessions from 14-leaf rosettes). **C. Inter**- vs **intra**-accession variability of expression in 14-leaf rosettes in the 28-accession dataset for two expression-length bins. **Inter**-accession variability: coefficient of variance of expression across 28 accessions with the expression value for each accession averaged from 2–4 replicates. **Intra**-accession variability: mean coefficient of variance of expression across 2–4 replicates for each accession (see Methods). Blue: significance of the difference between the **intra**-accession variability for different gene categories. Purple: significance of the difference between the **inter**-accession variability for different gene categories. **D.** Circadian (diurnal) expression variation calculated from Col-0 seedlings sampled every 2 hours during the day (Cortijo et al. 2019). The expression values are averaged from 14 technical replicates for each of the 12 time points. Only genes with TPM > 1 in at least one sample are plotted (see Methods).
Boxplots: Outliers are not plotted. *P*-values were calculated using Mann-Whitney tests on equalized sample sizes: *** $P < 10^{-10}$, ** $P < 10^{-5}$, * $P < 0.01$, n.s.: $P > 0.01$.

**Supplemental Figure S11** (Supports Figure 3)

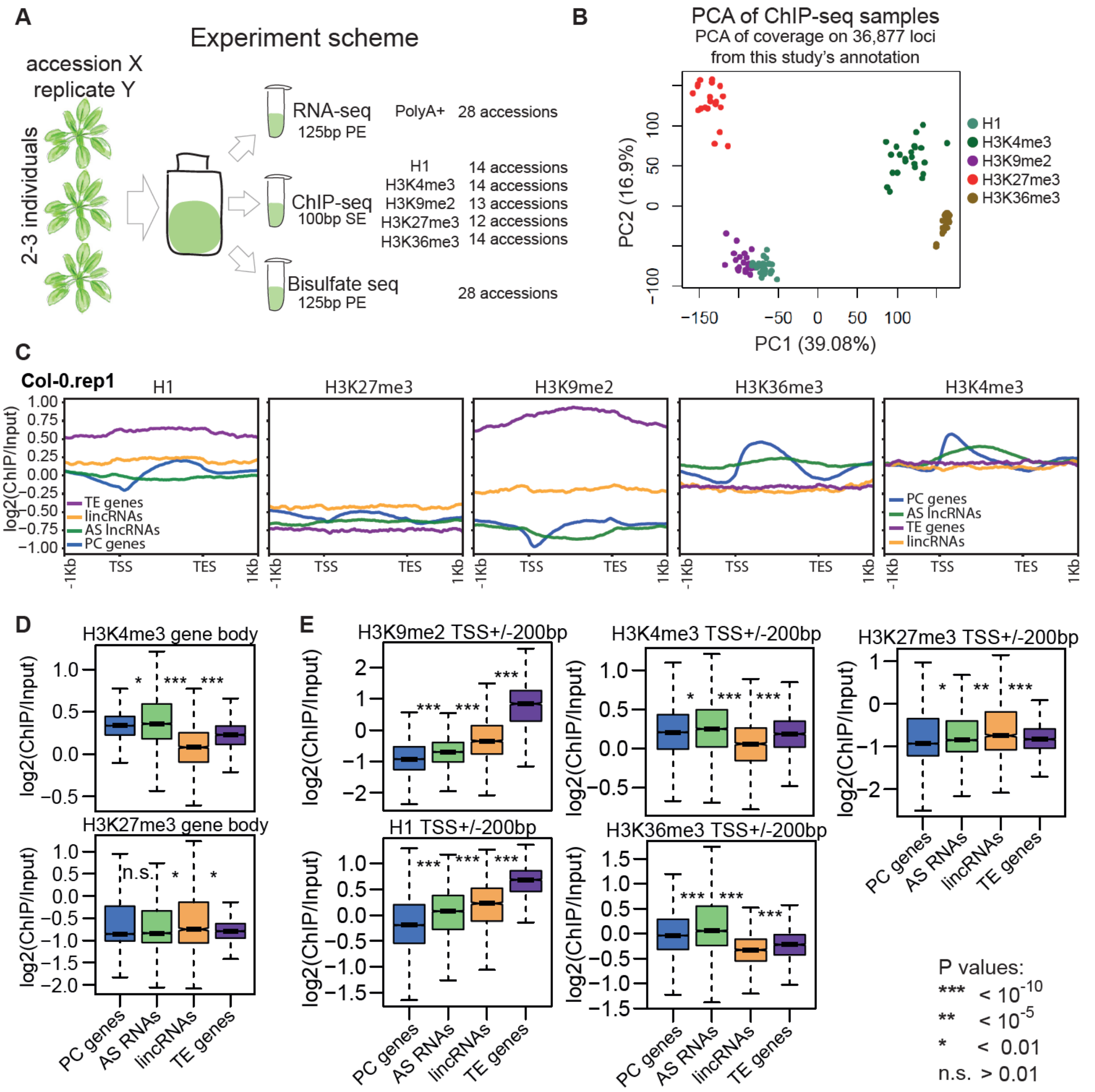


### Supplemental Figure S11. Histone mark profiling

**A.** Diagram of the samples collected in this study. Each replicate comes from independent experiments (plant growing) performed with a gap of few months. The leaf tissue from 2–3 individuals was frozen and ground and then parts of that were used for RNA-seq, ChIP-seq and Bisulfite-seq. **B.** Principal component analysis (PCA) results for ChIP-seq samples produced in this study. PCA was performed on coverage normalized by input. Coverage was calculated for PC gene, AS lncRNAs lincRNA, TE gene, TE fragment and AS_to_TE lncRNA loci from our cumulative annotation. For each sample, the average value between replicates was used for the PCA. **C.** Averaged profiles of the input-normalized ChIP-seq signal for histone H1, H3K9me2, H3K36me3, H3K4me3 and H3K27me3 levels over four gene types. Data from Col-0, mature leaves from 14-leaf rosette, replicate 1, are plotted. Profiles were built using plotProfile from deeptools. **D.** H3K4me3 (top) and H3K27me3 (bottom) histone modifications over the entire gene body in Col-0 rosettes. The log2 of the coverage normalized by input and averaged between the two replicates is plotted. **E.** H3K9me2, H3K4me3, H3K27me3, H1 and H3K36me3 histone modifications over promoters (TSS ± 200 bp) in Col-0 rosettes. The log2 of the coverage normalized by input and averaged between the two replicates is plotted. Boxplots: Outliers are not plotted. *P*-values were calculated using Mann-Whitney tests on equalized sample sizes: *** $P < 10^{-10}$, ** $P < 10^{-5}$, * $P < 0.01$, n.s.: $P > 0.01$.

### **Supplemental Figure S12** (Supports Figure 3)

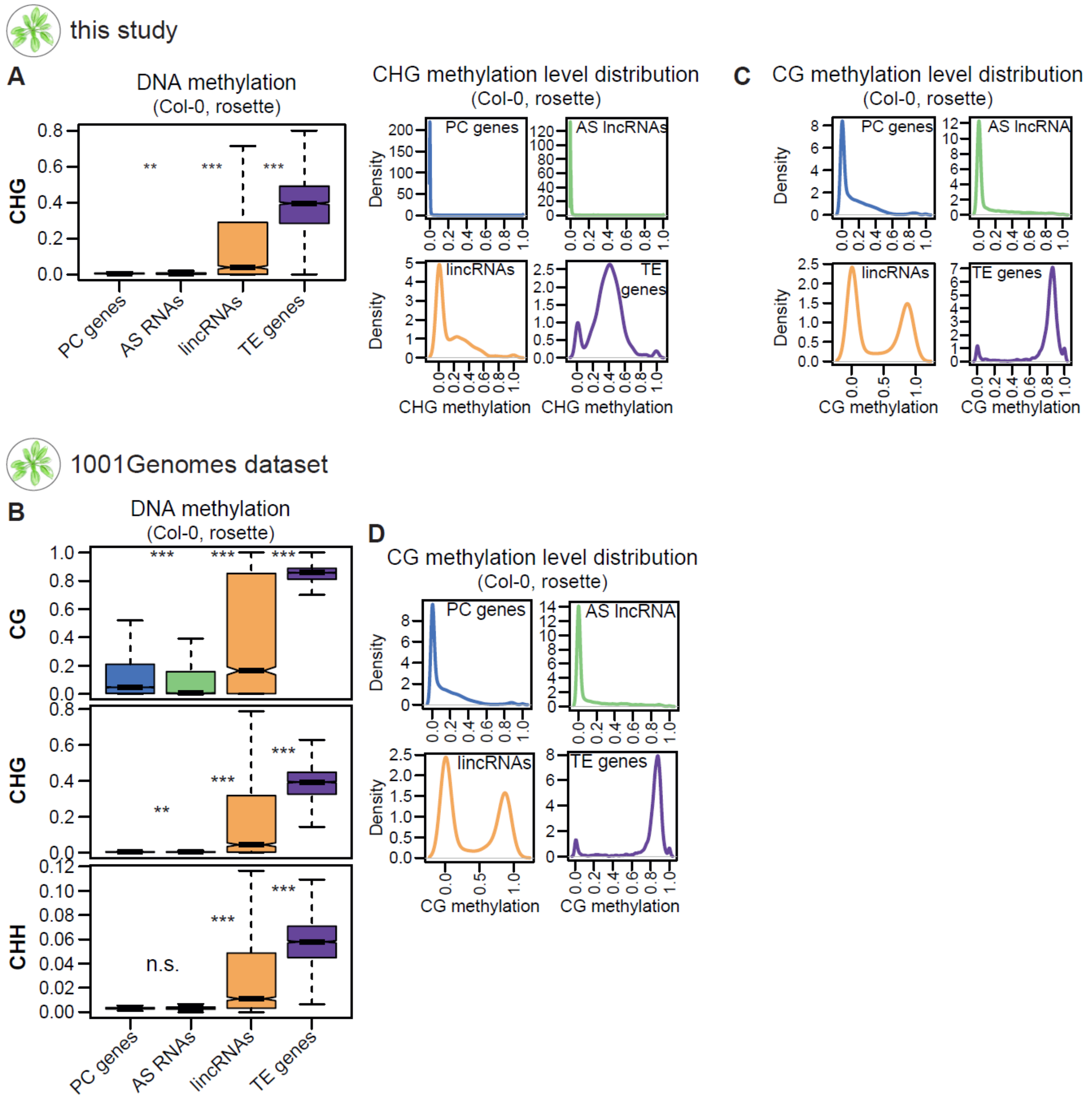


#### **Supplemental Figure S12. DNA methylation levels supplement**

**A.** Left, CHG DNA methylation level in Col-0 rosette (mature leaves from 14-leaf rosettes). Right, density of the CHG methylation level for PC gene, AS lncRNA, lincRNA and TE gene loci. Methylation levels were averaged from four replicates. **B.** CG, CHG and CHH DNA methylation level in Col-0 rosettes (mature leaves from 14-leaf rosettes), methylation data from the 1001 Genomes Project dataset (Kawakatsu et al. 2016). **C.** Density of the CG methylation level (Figure 3C) for PC gene, AS lncRNA, lincRNA and TE loci. **D.** Density of the CG methylation level for PC gene, AS lncRNA, lincRNA and TE loci. Methylation data from the 1001 Genomes Project dataset (Kawakatsu et al. 2016). Methylation level was calculated as the ratio between the number of methylated and unmethylated reads over all Cs in the respective context (CG or CHH) in the gene body.

### **Supplemental Figure S13** (Supports Figure 3)

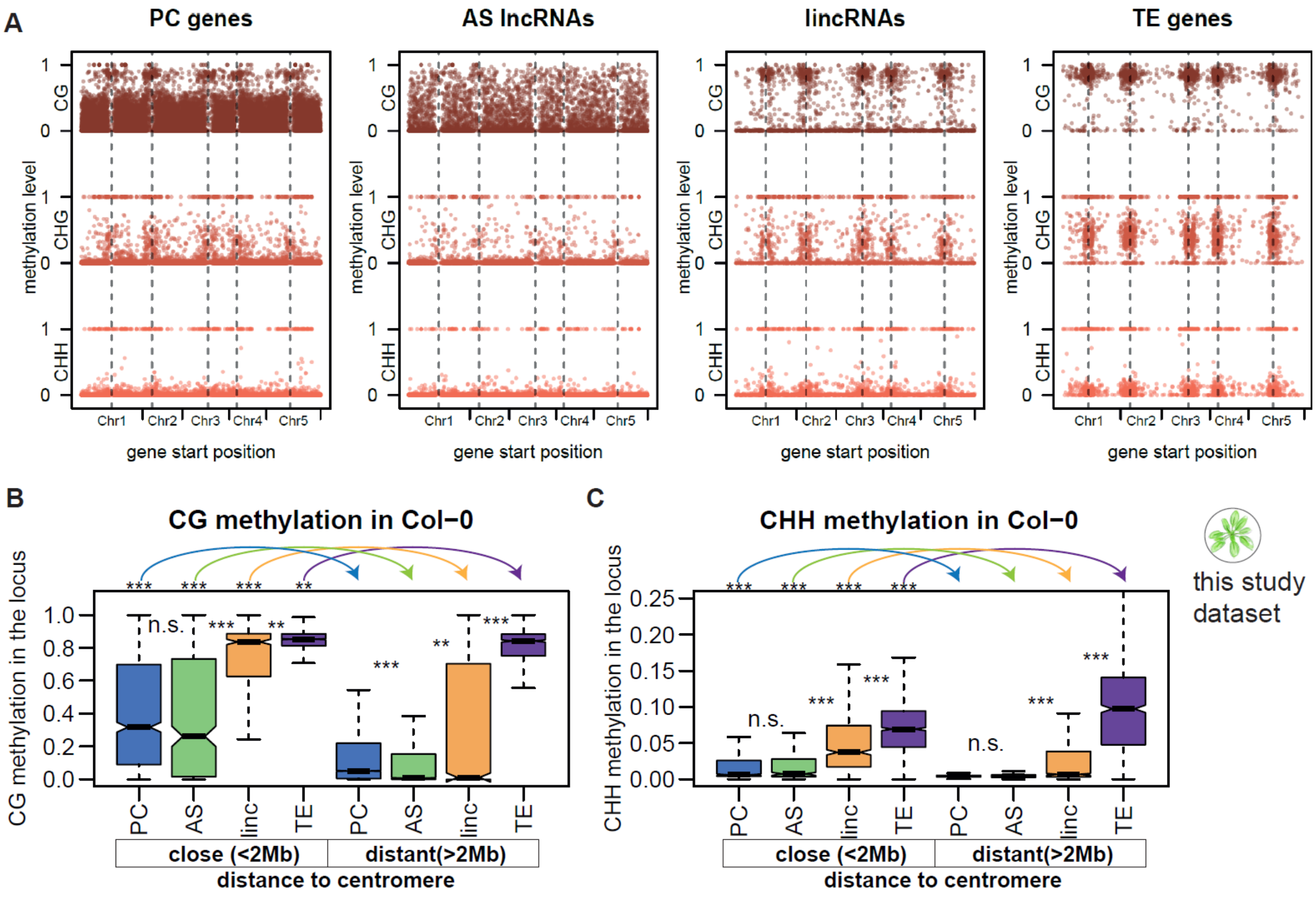


### Supplemental Figure S13. DNA methylation vs distance to the centromere

**A.** Scatterplots showing CG (top, brown), CHG (middle, terracotta) and CHH (bottom, red) methylation levels for genes with respect to their genomic position. Dashed vertical black lines indicate centromeres. **B-C**. CG (**B**) and CHH (**C**) methylation level in Col-0 rosettes for genes that are close to the centromere (the start of the gene is closer than 2 Mb to the centromere) and distant from the centromere (the start of the gene is further than 2 Mb to the centromere).

Outliers in the boxplot are not plotted. *P*-values were calculated using Mann-Whitney tests on equalized sample sizes: *** $P < 10^{-10}$, ** $P < 10^{-5}$, * $P < 0.01$, n.s.: $P > 0.01$.

The centromere positions used: Chr1: 15,000,000, Chr2: 4,700,000, Chr3: 13,000,000, Chr4: 3,800,000, Chr5: 11,900,000.

The methylation data displayed is from Col-0 rosettes from this study. The methylation levels from four replicates for Col-0 were averaged.

**Supplemental Figure S14** (Supports Figure 3)

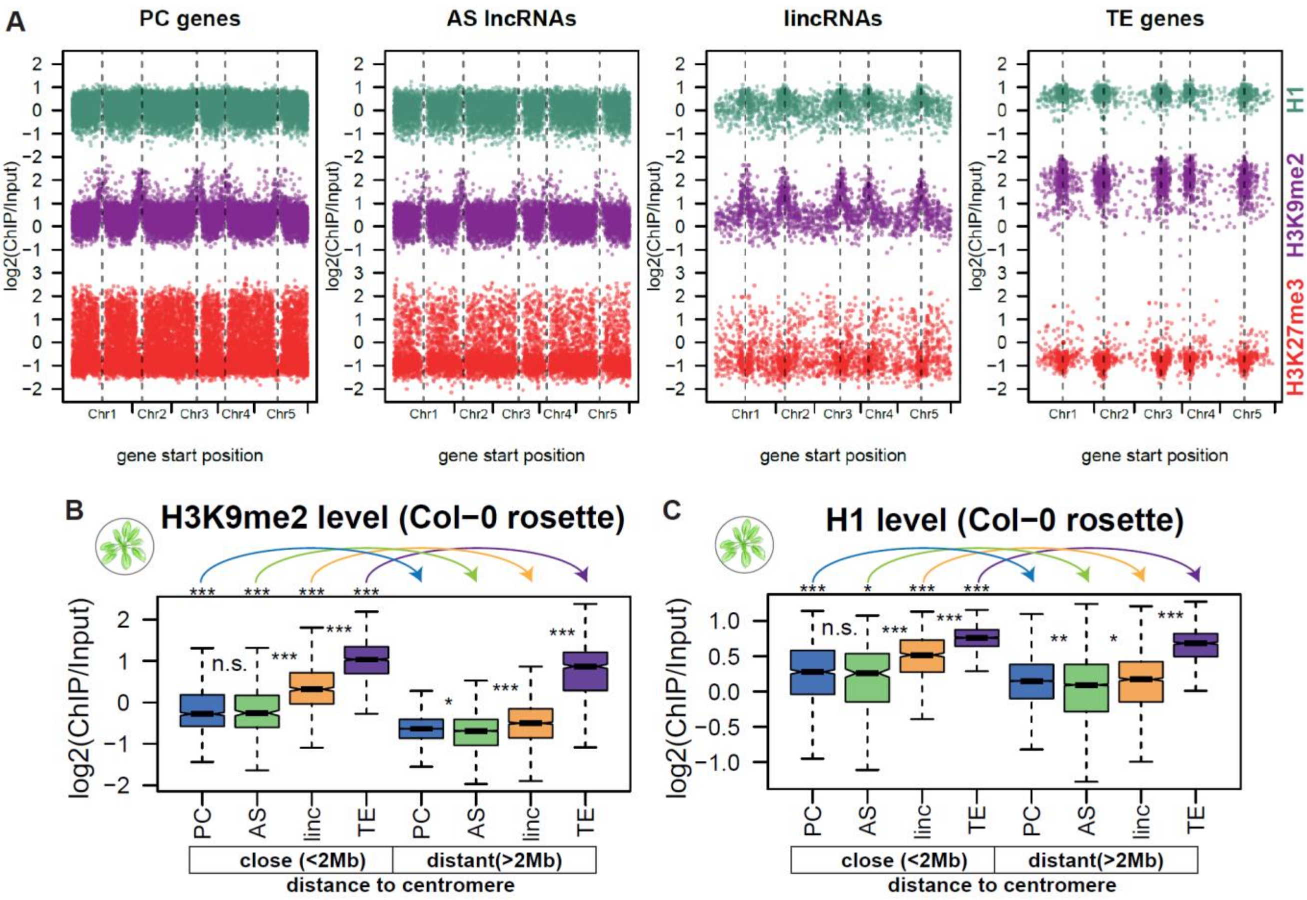


**Supplemental Figure S14. Heterochromatic histone marks vs distance to the centromere**

**A.** Scatterplots showing histone H1 (top, turquoise), H3K9me2 (middle, purple) and H3K27me3 (bottom, red) levels (normalized to input, averaged between the two replicates) for genes with respect to their genomic position. Dashed vertical black lines indicate centromeres. **B-C**. H3K9me2 (**B**) and histone H1 (**C**) levels in Col-0 rosettes for genes that are close to the centromere (the start of the gene is closer than 2 Mb to the centromere) and distant from the centromere (the start of the gene is further than 2 Mb to the centromere).

Outliers in the boxplot are not plotted. *P*-values were calculated using Mann-Whitney tests on equalized sample sizes: *** $P < 10^{-10}$, ** $P < 10^{-5}$, * $P < 0.01$, n.s.: $P > 0.01$.

The centromere positions used: Chr1: 15,000,000, Chr2: 4,700,000, Chr3: 13,000,000, Chr4: 3,800,000, Chr5: 11,900,000.

**Supplemental Figure S15** (Supports Figure 3)

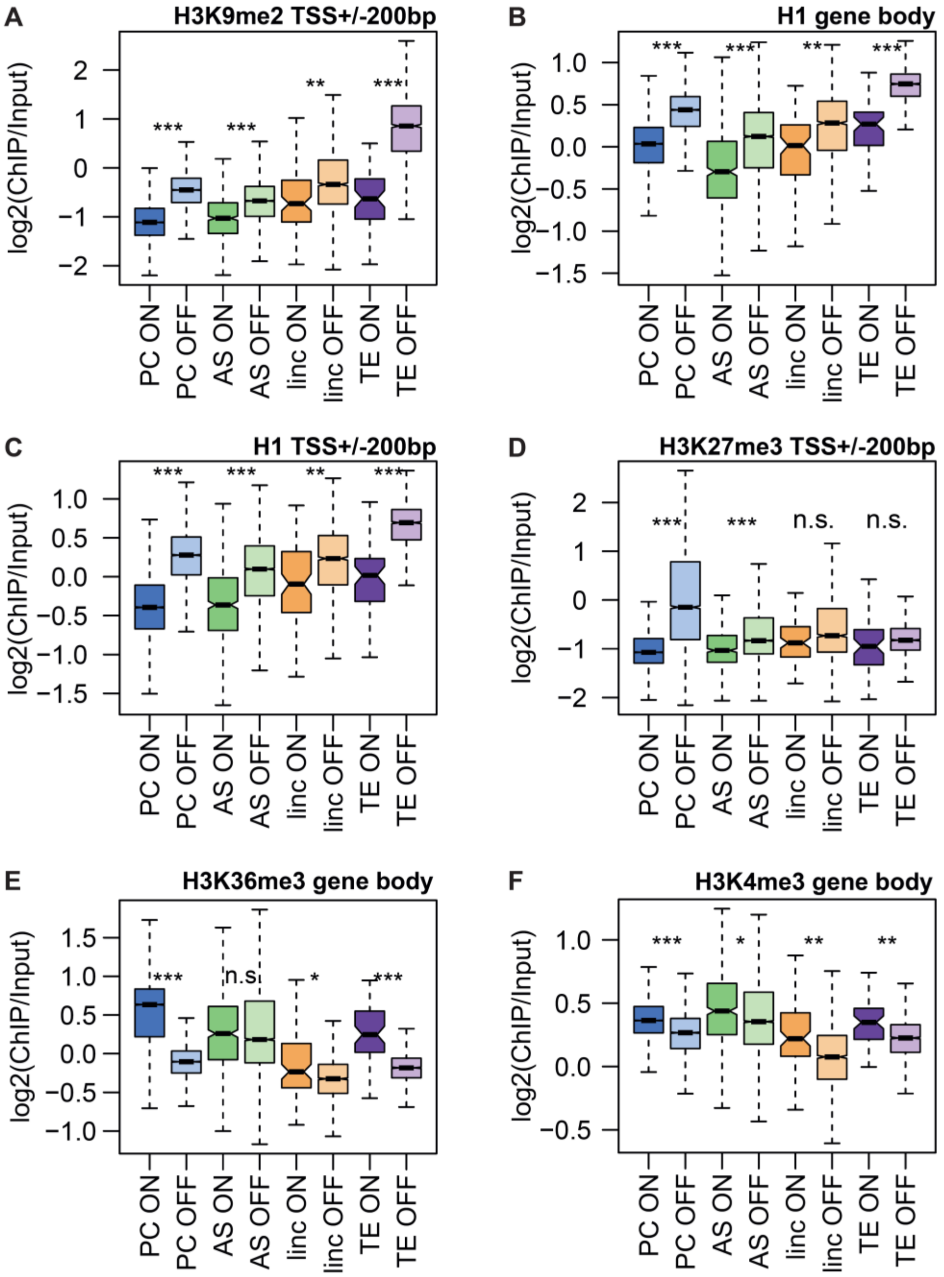


**Supplemental Figure S15. Histone modifications of silent and expressed genes: supplement**

**A.** H3K9me2 input-normalized coverage of the promoter (TSS+/-200bp) for expressed and silent genes. **B.** H1 input-normalized coverage of the gene for expressed and silent genes. **C.** H1 input-normalized coverage of the promoter (TSS+/-200bp) for expressed and silent genes. **D.** H3K27me3 input-normalized coverage of the promoter (TSS+/-200bp) for expressed and silent genes. **E.** H3K36me3 input-normalized coverage of the gene for expressed and silent genes. **F.** H3K4me3 input-normalized coverage of the gene for expressed and silent genes. Expressed and silent genes were defined from the expression of the matching samples of 14-leaf rosettes from Col-0 accession: ON: TPM>0.5, OFF: TPM<0.5. The expression was calculated as an average from 4 replicates. ChIP-seq coverage was averaged from 2 replicates. Outliers in the boxplots are not plotted. *P*-values were calculated using Mann-Whitney tests (equalized sample sizes for PC genes and AS lncRNAs): *** $P < 10^{-10}$, ** $P < 10^{-5}$, * $P < 0.01$, n.s.: $P > 0.01$.

**Supplemental Figure S16** (Supports Figure 3)

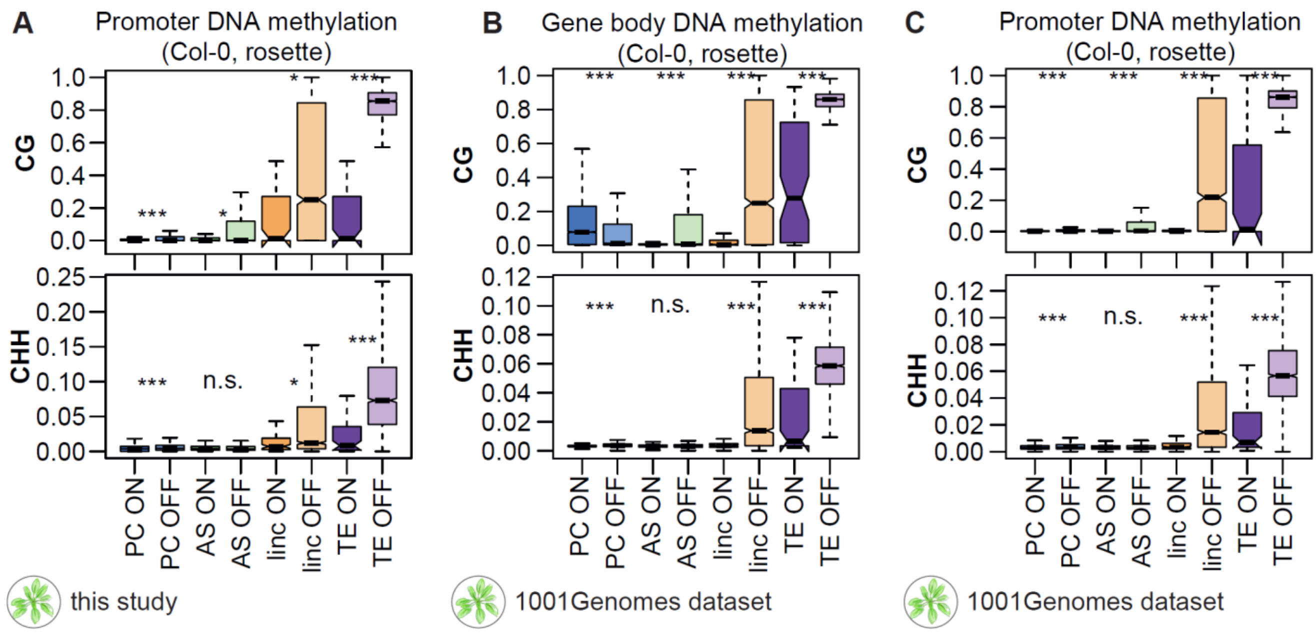


### Supplemental Figure S16. DNA methylation of silent and expressed genes: supplement

CG (top) and CHH (bottom) DNA methylation levels of expressed (ON, TPM>0.5) and silent (OFF, TPM<0.5) genes at their (**A**) promoters (TSS ± 200 bp), (**B**) gene bodies and (**C**) promoters. **A**: Expression and methylation data from Col-0 rosettes obtained in this study. **B** and **C**: Expression and methylation data from Col-0 rosettes from the 1001 Genomes Project dataset (Kawakatsu et al, 2016). Outliers are not plotted. *P*-values were calculated using Mann-Whitney tests (equalized sample sizes for PC genes and AS lncRNAs): *** $P < 10^{-10}$, ** $P < 10^{-5}$, * $P < 0.01$, n.s.: $P > 0.01$.

**Supplemental Figure S17** (Supports Figure 3)

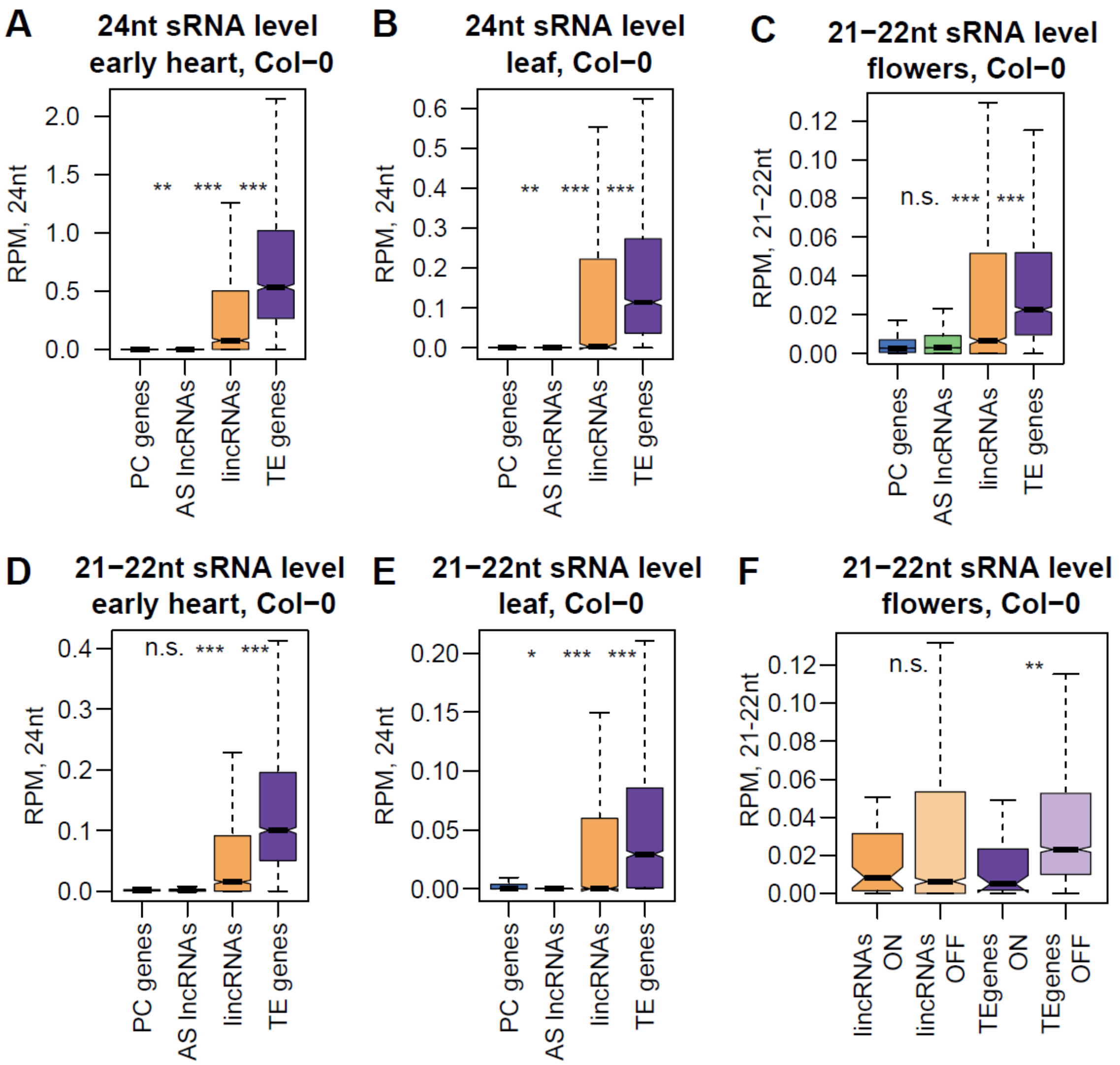


### Supplemental Figure S17. Coverage of 24-nt and 21–22-nt sRNA in early embryo and leaves

Coverage of 24-nt sRNA of PC gene, AS lncRNA, lincRNA and TE loci in **(A)** early heart (early embryonic stage) (left) and **(B)** leaves (right) in wild-type Col-0 accession (sRNA-seq data from (Papareddy et al. 2020)). The average of three replicates is plotted.
Coverage of 21–22-nt sRNA of PC gene, AS lncRNA, lincRNA and TE loci in Col-0 **(C)** flowers (this study, one replicate), **(D)** Col-0 early heart (early embryonic stage) and **(E)** leaves accession (sRNA-seq data from (Papareddy et al. 2020); the average of three replicates is plotted).
**F.** Coverage of 21–22-nt small RNA in Col-0 flowers plotted separately for expressed (ON, TPM>0.5) and silent (OFF, TPM<0.5) genes. Expression was calculated in the corresponding 9-leaf rosette samples.
Outliers are not plotted. *P*-values were calculated using Mann-Whitney tests on equalized sample sizes: *** $P < 10^{-10}$, ** $P < 10^{-5}$, * $P < 0.01$, n.s.: $P > 0.01$.

### **Supplemental Figure S18** (Supports Figure 3)

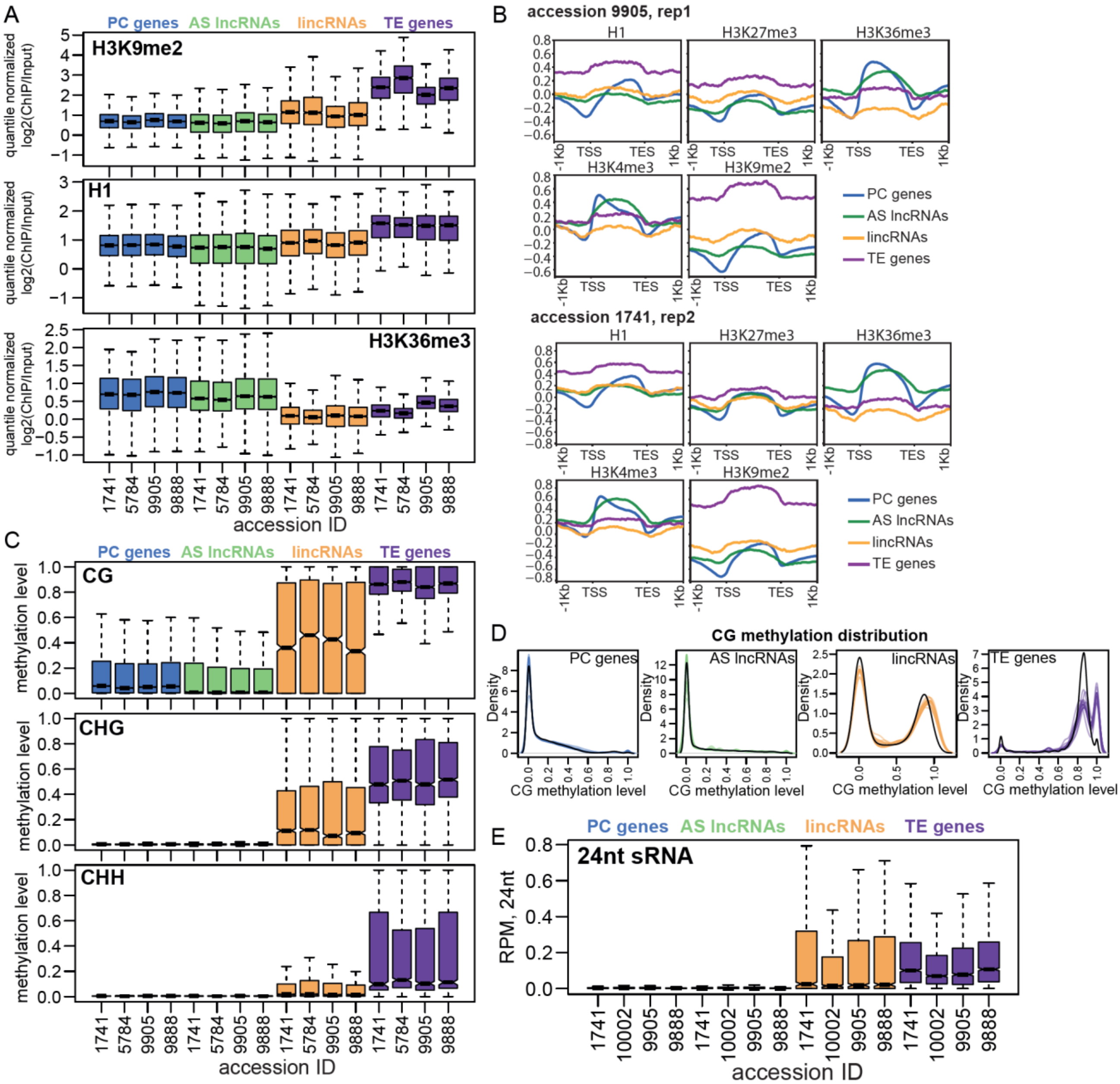


### **Supplemental Figure S18. Epigenetic patterns in non-reference accessions**

**A.** H3K9me2, histone H1 and H3K36me3 modifications in rosettes of four non-reference accessions. The histone modification levels were first log2-normalized by input, then normalized by scaling values from each accession to fit within the same range by applying the function: (x - quantile(x,.20)) / (quantile(x,.80) - quantile(x,.20)) in R. Quantile-normalized values were then averaged between the replicates for each accession. **B.** Averaged profiles (deeptools-built) of log2 input-normalized ChIP-seq signal for five histone marks over four gene types. Data from rosettes from acc. 9905 (replicate 1) and acc 1741 (replicate 1) are plotted. **C.** CG, CHG and CHH DNA methylation levels in rosettes of four non-reference accessions. Methylation level was calculated as the ratio between the number of methylated and unmethylated reads over all Cs in the respective context (CG or CHH) in the locus and averaged between 2–4 replicates. **D.** Density of CG methylation level in 13 nonreference accessions (pale lines) and in Col-0 (black line). **E.** Coverage of 24-nt small RNAs in the gene body for flowers from four non-reference accessions, calculated as the number of 24-nt reads mapping to the locus divided by the total number of reads and the locus length.

**Supplemental Figure S19** (Supports Figure 4)

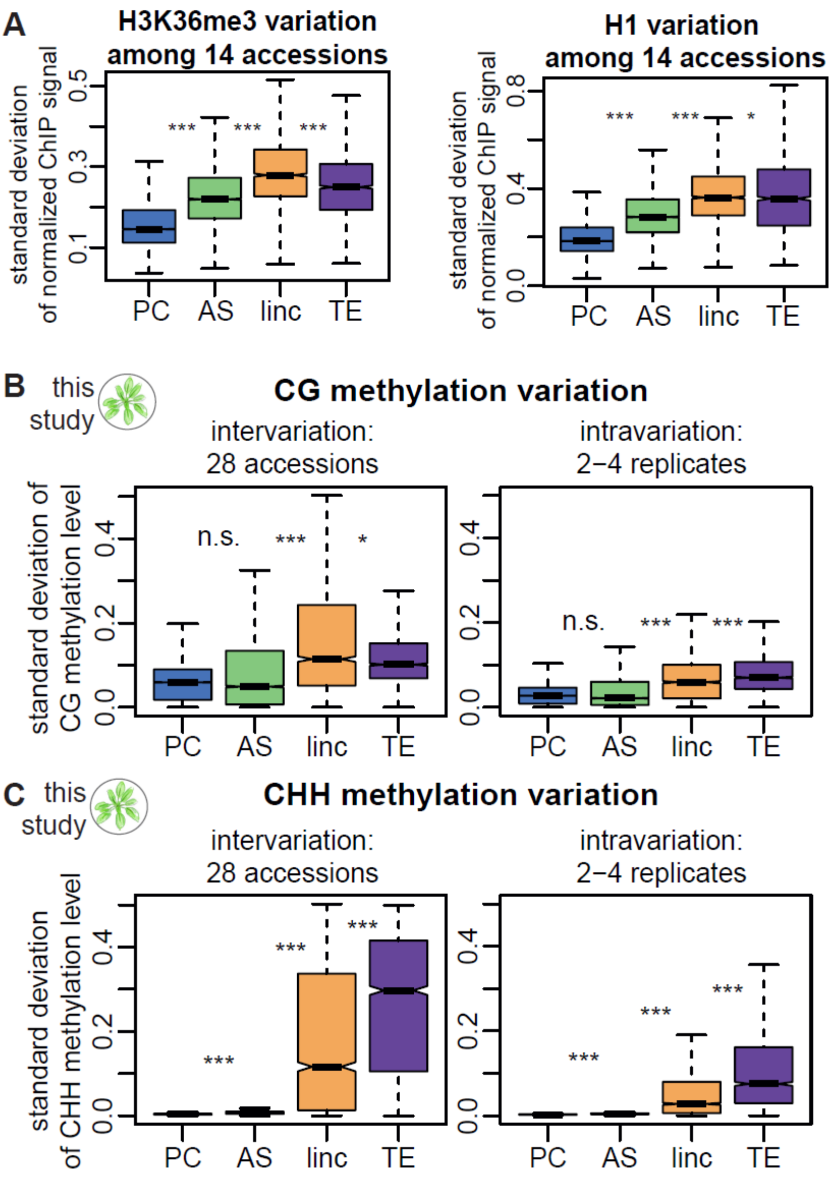


**Supplemental Figure S19. Epigenetic variation supplement 1**

**A.** Variation in H3K36me3 (left) and H1 (right) levels in mature leaves (14-leaf rosette) for the four gene categories from our cumulative annotation. The histone modification levels were first log2-normalized by input, then normalized by scaling values from each accession to fit within the same range by applying the function: (x - quantile(x,.20)) / (quantile(x,.80) - quantile(x,.20)) in R. The standard deviation between accessions was calculated across quantile-normalized values averaged across replicates for each accession. **B-C.** Left, Standard deviation of CG (**B**) and CHH (**C**) methylation levels across 28 accessions. Methylation level for each accession was calculated as the ratio between the number of methylated and unmethylated reads over all Cs in the respective context (CG or CHH) in the locus and averaged between 2–4 replicates. Right, Standard deviation of CG (**B**) and CHH (**C**) methylation levels across 2–4 replicates: average across 28 accessions is plotted. *P*-values were calculated using Mann-Whitney test on equalized sample sizes: *** $P < 10^{-10}$, ** $P < 10^{-5}$, * $P < 0.01$, n.s.: $P > 0.01$.

**Supplemental Figure S20** (Supports Figure 4)

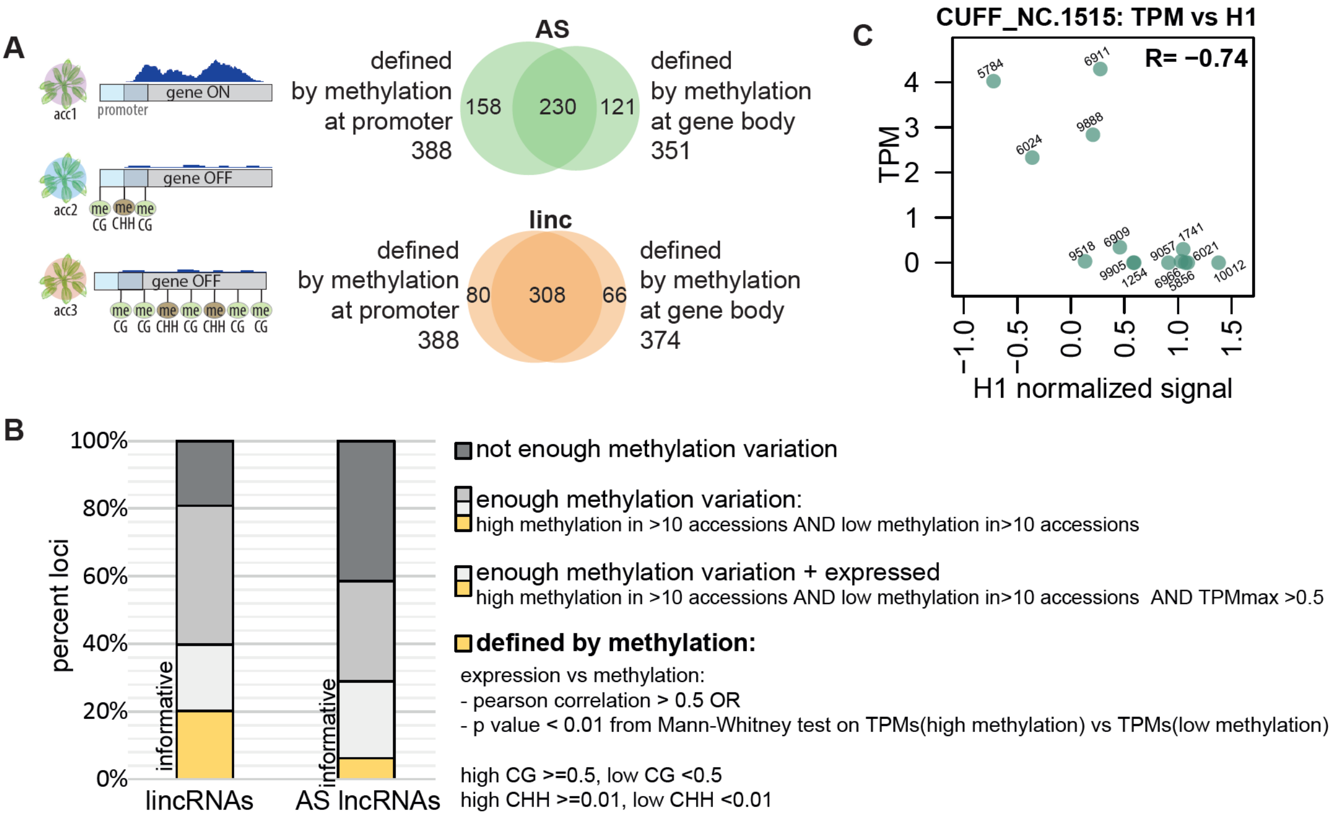


**Supplemental Figure S20.** Epigenetic variation supplement 2

**A.** Summary of lncRNAs for which expression can be explained by methylation (see Methods). The Venn diagrams show the overlap between loci for AS lncRNAs (green) and lincRNAs (orange) that were found to be defined by CG or CHH methylation level at their promoter (TSS ± 200 bp) or gene body. **B.** Expression in rosettes as a function of H1 levels in rosettes for lincRNA CUFF_NC.1515 in 13 accessions. **C.** Distribution of the informativeness of lincRNA and AS lncRNA loci and the corresponding criteria.

**Supplemental Figure S21** (Supports Figure 5)

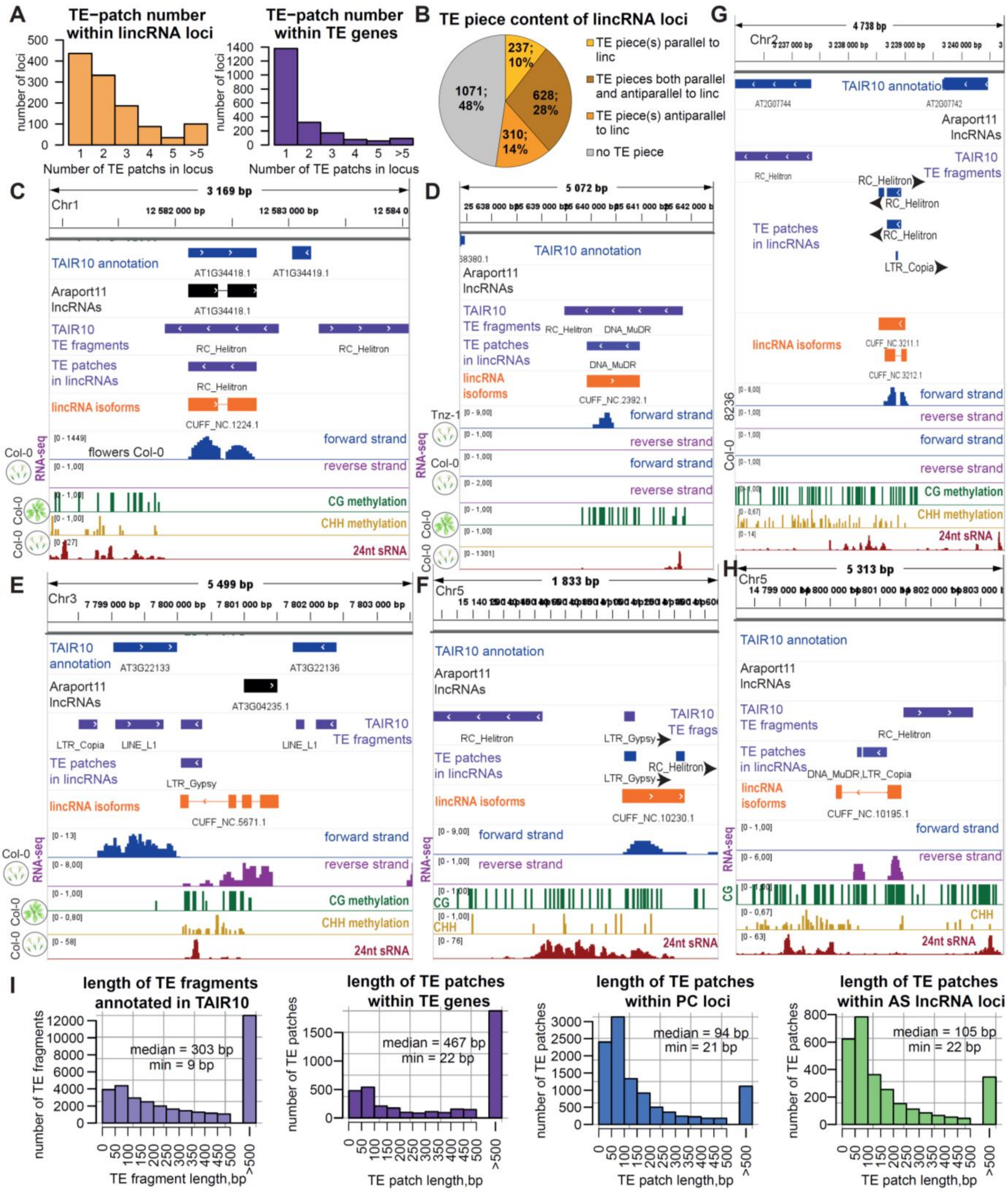


**Supplemental Figure S21. TE patches in lincRNAs**

**A.** Distribution of the number of TE patches inside lincRNA loci (left) and TE genes (right). Patches that are sense or antisense to the lincRNA direction were both counted. **B.** Distribution of lincRNAs with regard to their TE content and the direction of the TE pieces/patches. Parallel is sense to the lincRNA direction, antiparallel is antisense to the lincRNA direction. **C, D.** IGV screenshots showing two examples of lincRNAs that are fully overlapped by a TAIR10 TE fragment in the antisense direction. **E, F.** IGV screenshots showing two examples of lincRNAs that overlapped a TAIR10 TE fragment in the sense direction. **G, H.** IGV screenshots showing two examples of lincRNAs that do not overlap any TAIR10 TE fragments but do contain TE patches. **I.** Length distribution of (from left to right) TAIR10-annotated TE fragments, TE patches within TE genes, TE patches within PC genes and TE patches within AS lncRNAs. (PC genes, TE genes and AS lncRNAs as annotated by our cumulative transcriptome annotation).

### **Supplemental Figure S22** (Supports Figure 5)

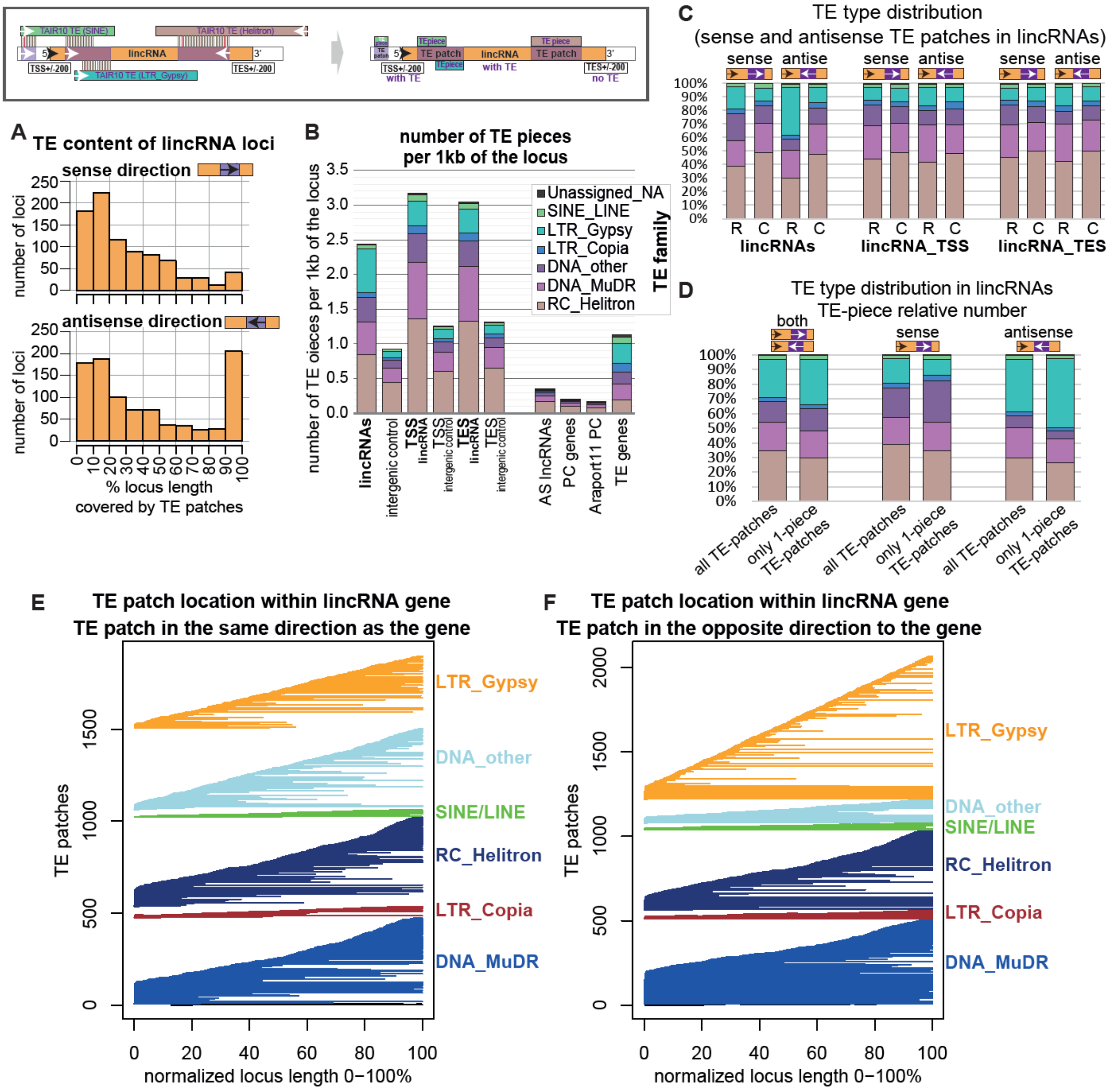


**Supplemental Figure S22. Sense and antisense TE pieces in lincRNAs: content, family, position**

**A.** Distribution of the relative length of the lincRNA loci with sense (top) or antisense (bottom) TE patches. **B.** Number of TE-pieces of different TE families per 1 kb of different types of loci. For lincRNA loci and border regions controls were performed by randomly shuffling the regions within intergenic space of the TAIR10 genome (the shuffling was done three times and the results were averaged between these three replicates). **C.** Proportion of sense and antisense TE-pieces of different TE families within lincRNA gene body, promoter (TSS ± 200 bp) and downstream border (TES ± 200 bp). “R” stands for real loci and “C” stands for controls. **D.** As we plotted the distribution of pieces in (**B**) and (**C**), and TE-patches can have many TE-pieces of the same type that are identical to each other thus bloating the numbers, we controlled for it by plotting the TE-family distribution for all lincRNAs with TE-patches and lincRNAs with TE patches that only contained one TE piece. While there is difference, the main trends are preserved. **E, F.** Position of sense (**E**) and antisense (**F**) TE patches of different TE-families within lincRNA loci.

**Supplemental Figure S23** (Supports Figure 5)

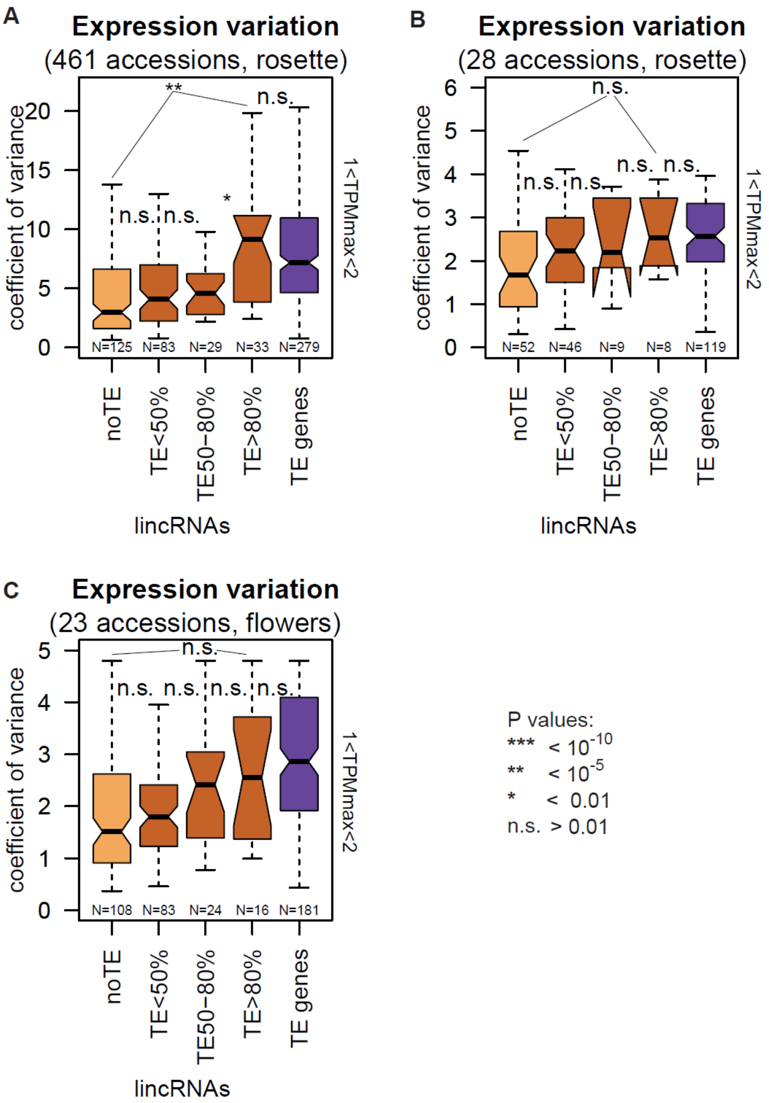


**Supplemental Figure S23. Expression variation vs TE content: expression control**

Expression variability for lincRNAs as a function of TE content, with TE genes for comparison. (**A**) across 461 accessions: rosette dataset from the 1001 Genomes Project dataset (Kawakatsu et al), (**B**) across 28 accessions: rosette dataset from this study, (**C**) across 23 accessions: flower dataset from this study. *P*-values were calculated using Mann-Whitney tests: *** $P < 10^{-10}$, ** $P < 10^{-5}$, * $P$ <0.01, n.s.: $P$ >0.01. Outliers in the boxplots are not shown.

**Supplemental Figure S24** (Supports Figure 5)

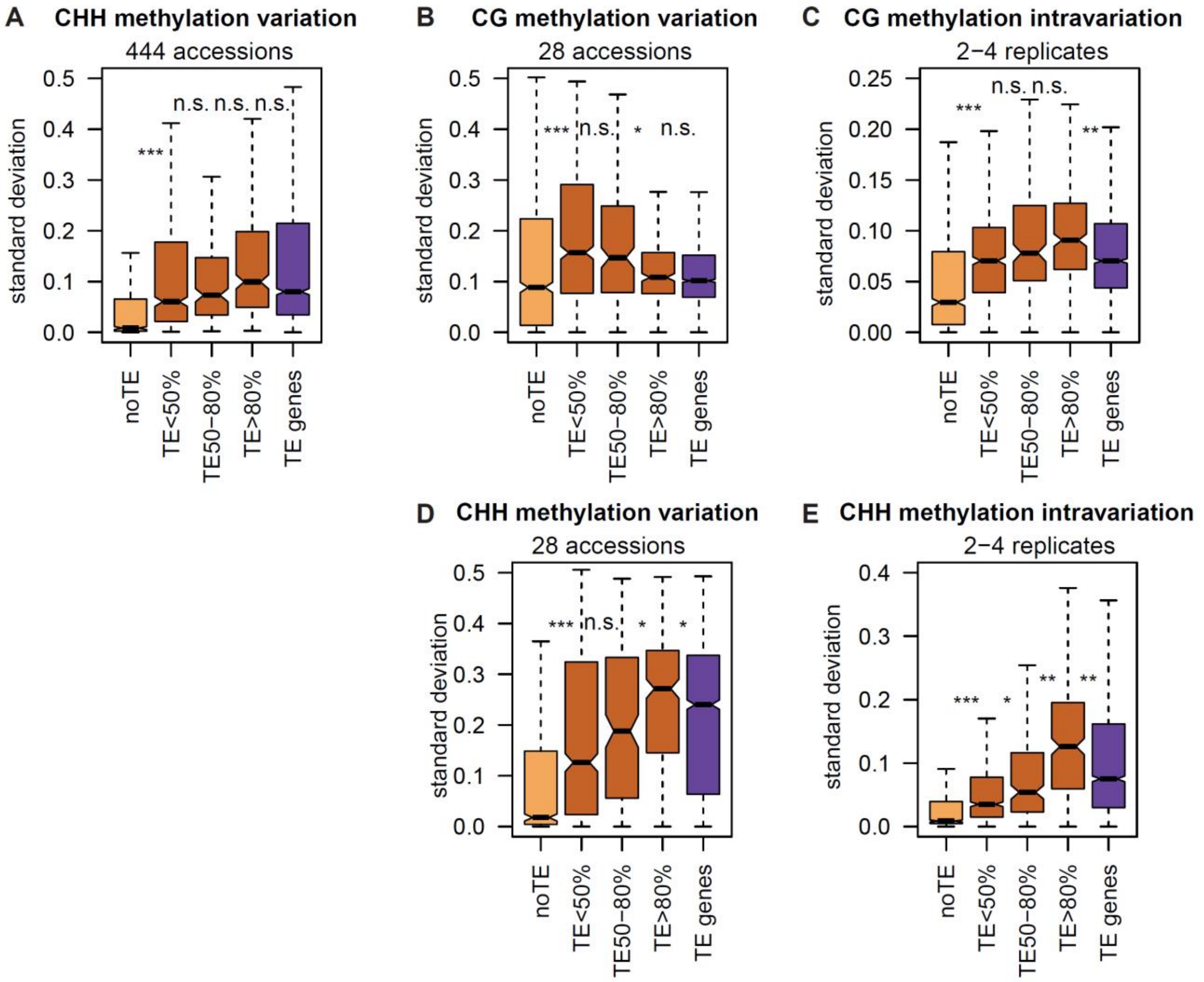


### Supplemental Figure S24. lincRNA methylation variation vs. TE content: supplement

Standard deviation of CG methylation levels across 444 accessions (Kawakatsu et al. 2016) (**A**), CG methylation levels across 28 accessions (values for each accession are averaged from 2-4 replicates) (**B**), CG methylation levels across 2-4 replicates (averaged across 28 accessions) (**C**), CHH methylation levels across 28 accessions (values for each accession are averaged from 2–4 replicates) (**D**), CHH methylation levels across 2–4 replicates (averaged across 28 accessions) (**E**) for lincRNAs as a function of TE content, with TE genes for comparison. *P*-values were calculated using Mann-Whitney tests: *** $P < 10^{-10}$, ** $P < 10^{-5}$, * $P < 0.01$, n.s.: $P > 0.01$. Outliers are not shown.

**Supplemental Figure S25** (Supports Figure 5)

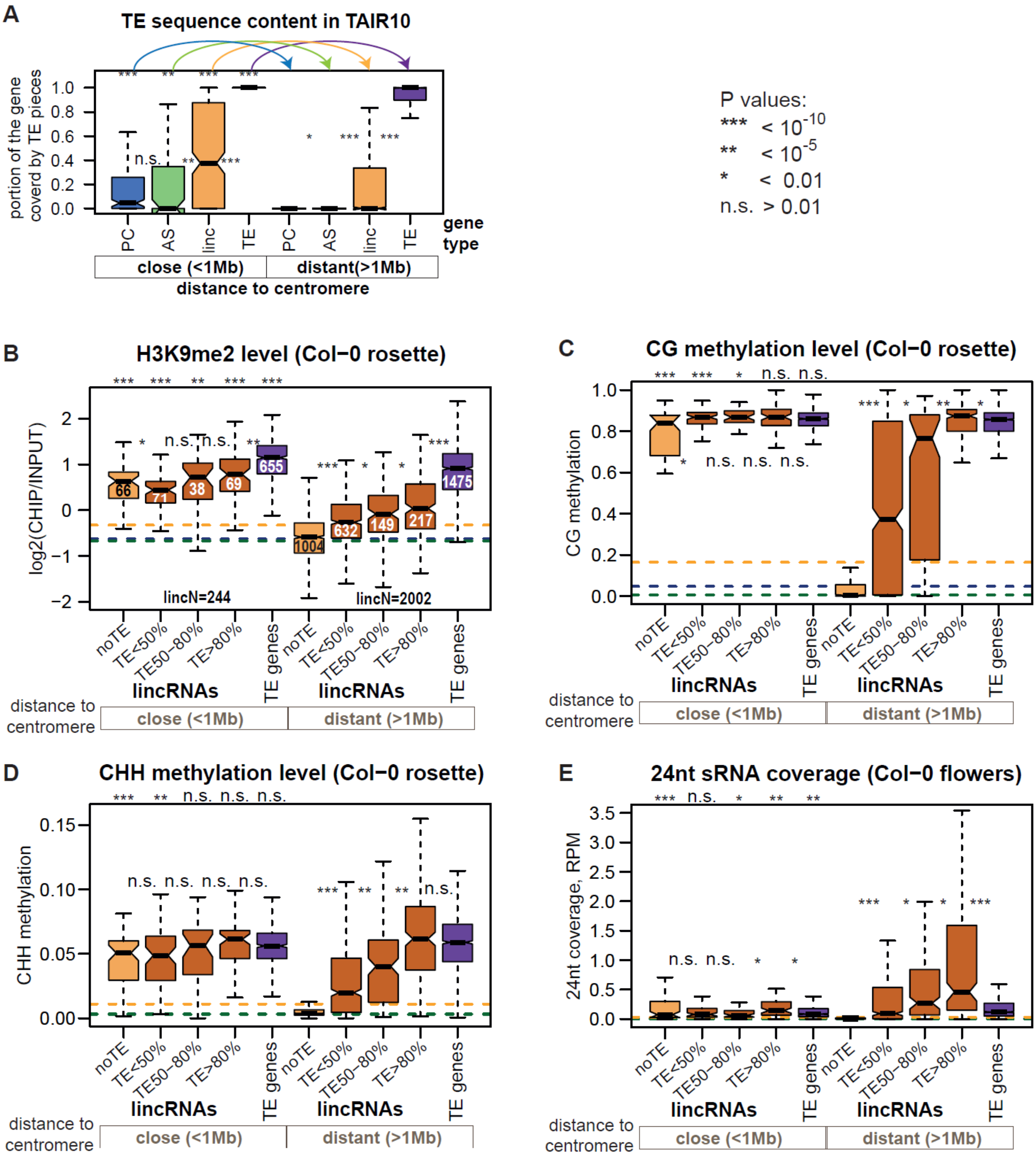


### Supplemental Figure S25. TE pieces affect epigenetics when controlled for chromosomal location

TE content (**A**), H3K9me2 level in rosettes (**B**), CG methylation level in rosettes (**C**), CG methylation level in rosettes (**D**), and 24-nt sRNA coverage in flowers (**E**) for different types of genes, split into those close and distant from the centromeres. For all boxplots: the stars and colored arrows on top of the plot indicate comparisons between “close” and “distant” genes of the same type. *P*-values were calculated using Mann-Whitney tests: *** $P < 10^{-10}$, ** $P < 10^{-5}$, * $P$ <0.01, n.s.: $P$ >0.01. Outliers in the boxplots are not shown. The centromere positions used: Chr1: 15,000,000, Chr2: 4,700,000, Chr3: 13,000,000, Chr4: 3,800,000, Chr5: 11,900,000. The methylation data displayed are from Col-0 rosettes from (Kawakatsu et al. 2016).

**Supplemental Figure S26** (Supports Figure 6)

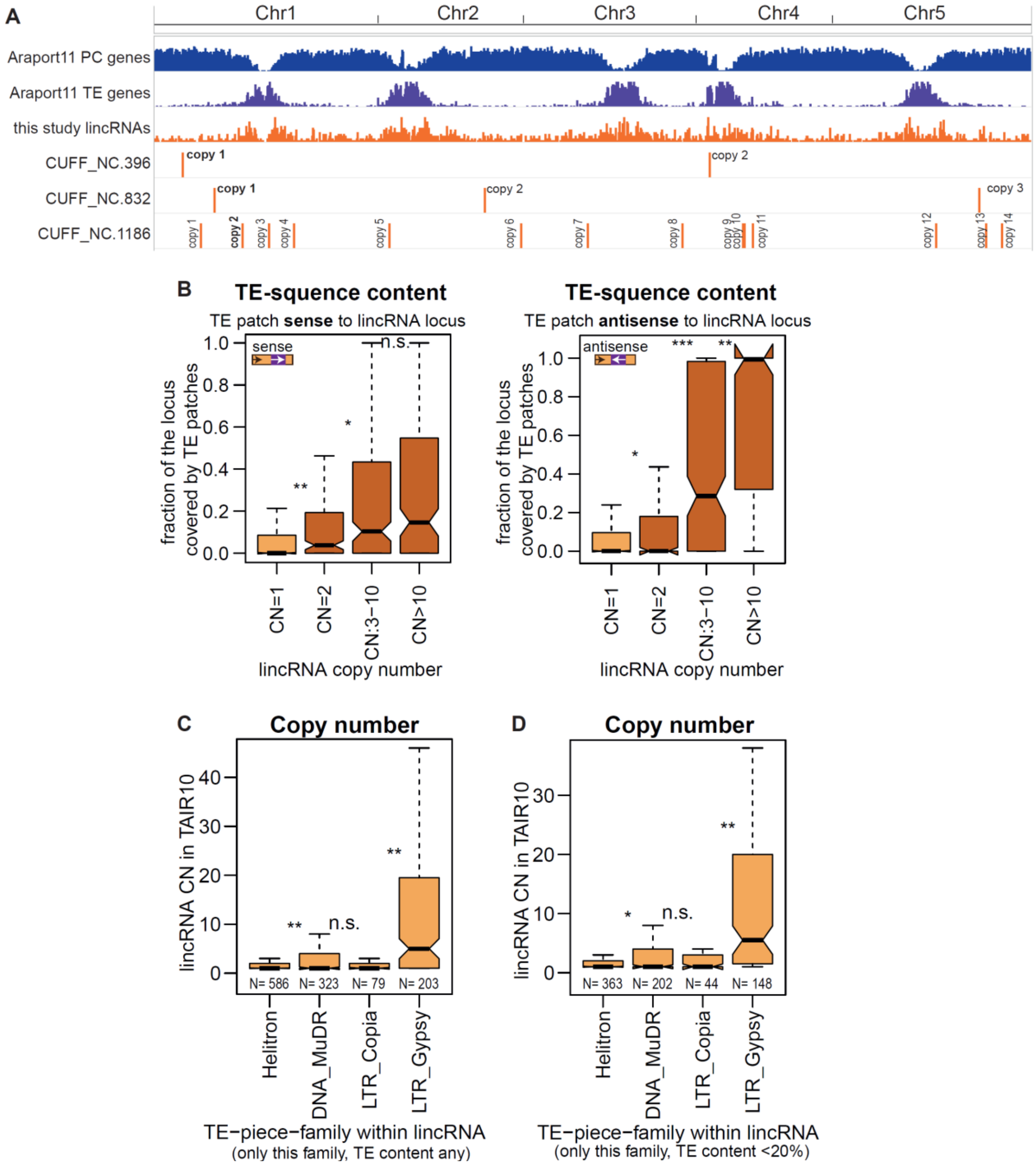


### Supplemental Figure S26. Copy number supplement

**A.** Genomic location of three examples of lincRNAs with multiple copies. Copy 1 indicates the position of the originally identified lincRNA locus, while other copies indicate the results of our BLAST-based copy search. **B.** Fraction of the lincRNA locus occupied by TE patches in sense (left) and antisense (right) direction for lincRNA loci with different copy number in the TAIR10 genome. **C-D.** Copy number of lincRNAs with TE pieces from four different types for lincRNAs with any TE content (C) and lincRNAs with no more than 20% of the locus occupied by TE pieces (D). Only lincRNAs with TE-pieces of one type are plotted. Numbers below boxes indicate the number of lincRNA loci in each category. TE content was calculated irrespective of the direction of TE pieces. *P*-values were calculated using Mann-Whitney tests: *** $P < 10^{-10}$, ** $P < 10^{-5}$, * $P < 0.01$, n.s.: $P > 0.01$. Outliers in the boxplots are not shown.

### **Supplemental Figure S27** (Supports Figure 6)

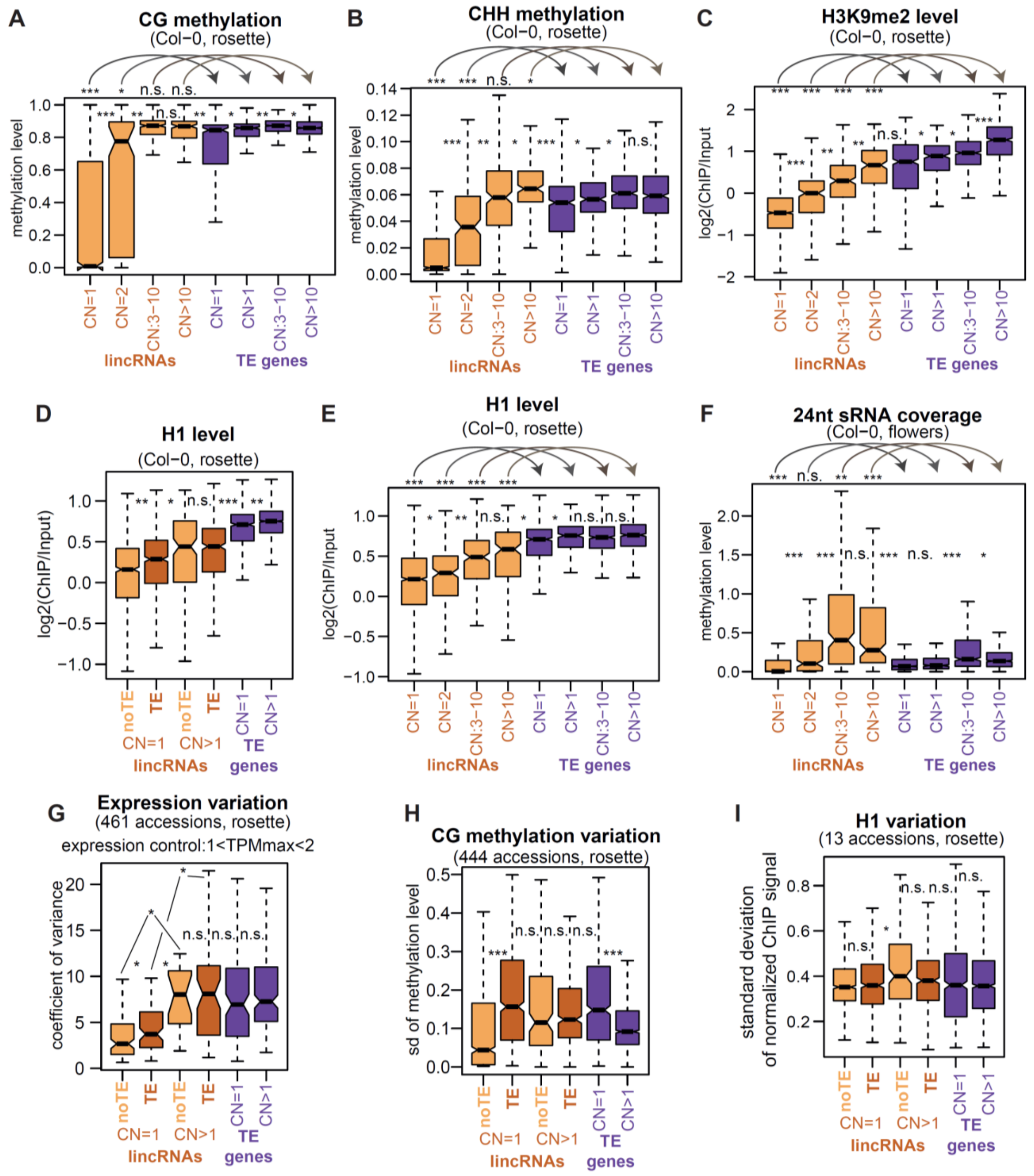


**Supplemental Figure S27. Copy number affects lincRNA epigenetic pattern: supplement**

**A, B, C, E, F.** Boxplots showing the CG methylation (**A**), CHH methylation (**B**), H3K9me2 (**C**) and H1 (**E**) levels in Col-0 rosettes, and 24-nt sRNA coverage in Col-0 flowers (**F**) for lincRNAs and TE genes with different copy number in the TAIR10 genome. **D, G, H, I.** The boxplots show H1 level in Col-0 rosettes (**D**), expression variation across 461 accessions (1<TPMmax<2) (Kawakatsu et al. 2016) (**G**), CG methylation (Kawakatsu et al. 2016) (**H**) and H1 level (**I**) variability for the four types of lincRNAs and the two types of TE genes. *P*-values in the boxplots are calculated using a Mann-Whitney test: *** $P < 10^{-10}$, ** $P < 10^{-5}$, * $P < 0.01$, n.s.: $P > 0.01$. Outliers in the boxplots are not plotted**.**

**Supplemental Figure S28** (Supports Figure 7)

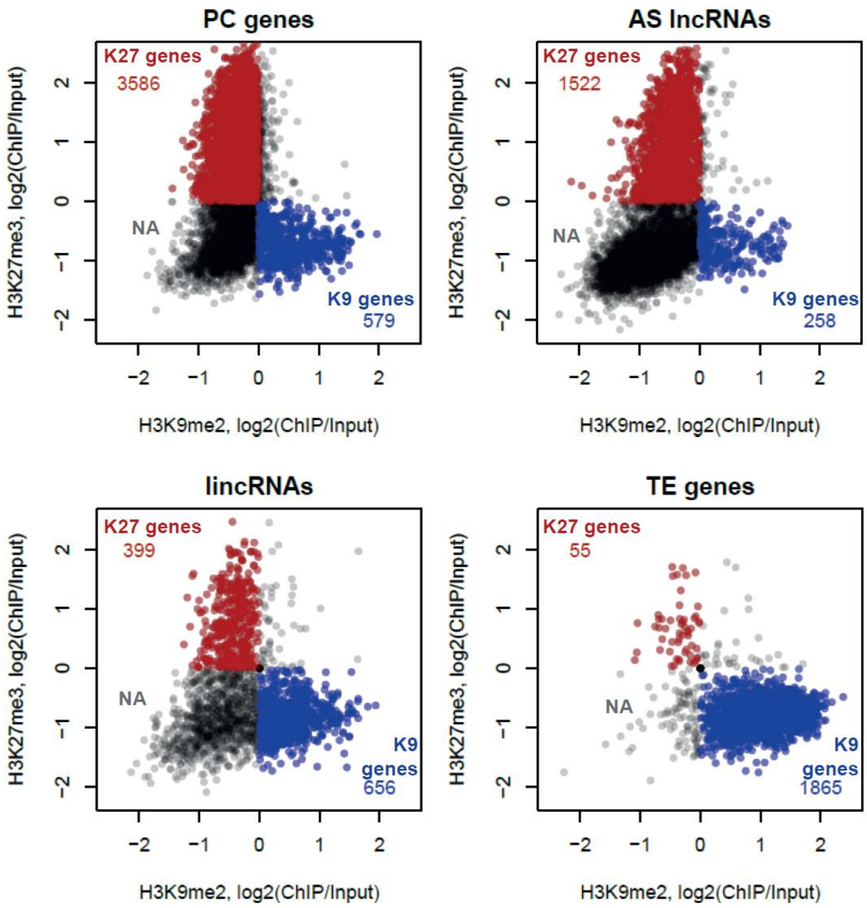


**Supplemental Figure S28. H3K27me3 and H3K9me2 dichotomy**

Scatterplots showing the H3K27me3 (y-axis) as a function of H3K9me2 (x-axis) levels in mature leaves (14-leaf rosette) of Col-0 for PC genes, AS lncRNAs, lincRNAs and TE genes from our cumulative annotation. The histone modification level was calculated over the entire gene body and was normalized to the input; the levels from two ChIP-seq replicates were averaged. K27 genes (red) are defined as those with the H3K27me3 level >0 and H3K9me2 level <0. K9 genes (blue) are defined as those with the H3K27me3 level >0 and H3K9mn2 level <0. NA genes (grey) are the rest of genes.

**Supplemental Figure S29** (Supports Figure 7)

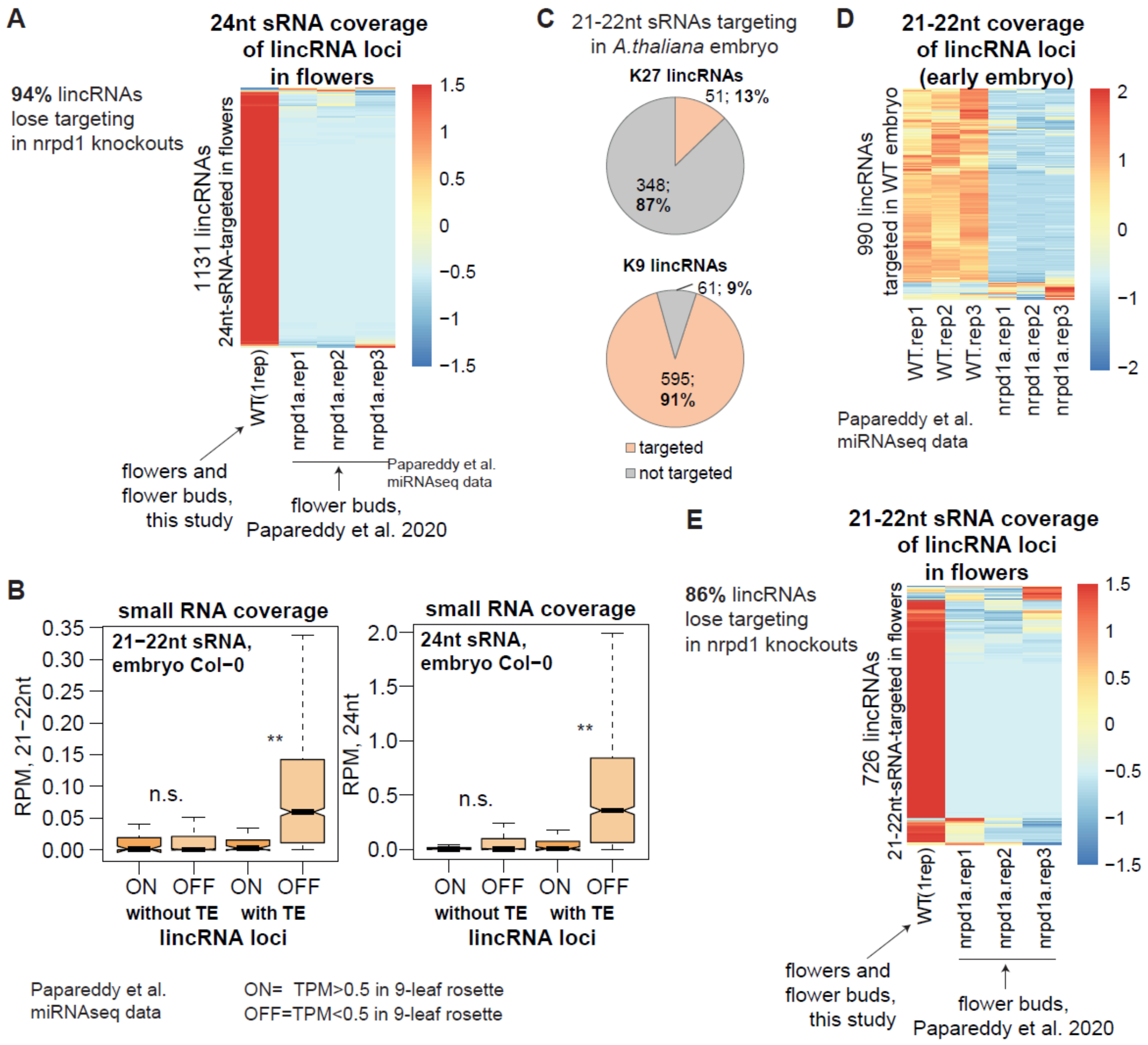


### Supplemental Figure S29. Loss of sRNA targeting *nrpd1a* mutants

**A.** Coverage of 24-nt sRNA in Arabidopsis flowers in the wild type (WT, Col-0) and in Pol IV-deficient mutants (*nrpd1a*, Col-0 background) (Papareddy et al. 2020). 1,131 lincRNAs that are targeted (RPM>0.03) by 24-nt sRNAs in the WT flowers are plotted. As WT flower bud data from (Papareddy et al. 2020) were not available, we used our own Col-0 data from flower tissue that contained flowers and flower buds. **B.** Normalized coverage by 21–22-nt (left) and 24-nt (right) small RNAs of ON (TPM>0.5) and OFF (TPM<0.5) lincRNA loci with and without TE pieces (Methods). Coverage was calculated as average of three replicates of WT “early heart” (Papareddy et al. 2020). *P*-values calculated using Mann-Whitney test: $^{**}P<10^{-5}$, $^{*}P<0.01$, n.s. $P>0.01$. Outliers not plotted**.** **C.** Relative number of K27 and K9 lincRNAs targeted by 21–22-nt sRNAs (RPM>0.03) in Arabidopsis embryos (“early heart” stage) (Papareddy et al., 2020). The small RNA coverage is averaged across three replicates. **D.** Coverage of 21–22-nt sRNA in Arabidopsis embryos (“early heart” stage) in the wild type (WT, Col-0) and in Pol IV-deficient mutants (*nrpd1a*, Col-0 background) (Papareddy et al., 2020). 990 lincRNAs that are targeted (RPM>0.03, average of three replicates) by 21–22-nt sRNAs in the WT are plotted. **E.** 21–22-nt sRNA coverage in Arabidopsis flowers in the wild type (WT, Col-0) and in Pol IV-deficient mutants (*nrpd1a*, Col-0 background) (Papareddy et al. 2020). 726 lincRNAs that are targeted (RPM>0.03) by 21–22-nt sRNAs in the WT flowers are plotted.

**Supplemental Figure S30** (Supports Figure 7)

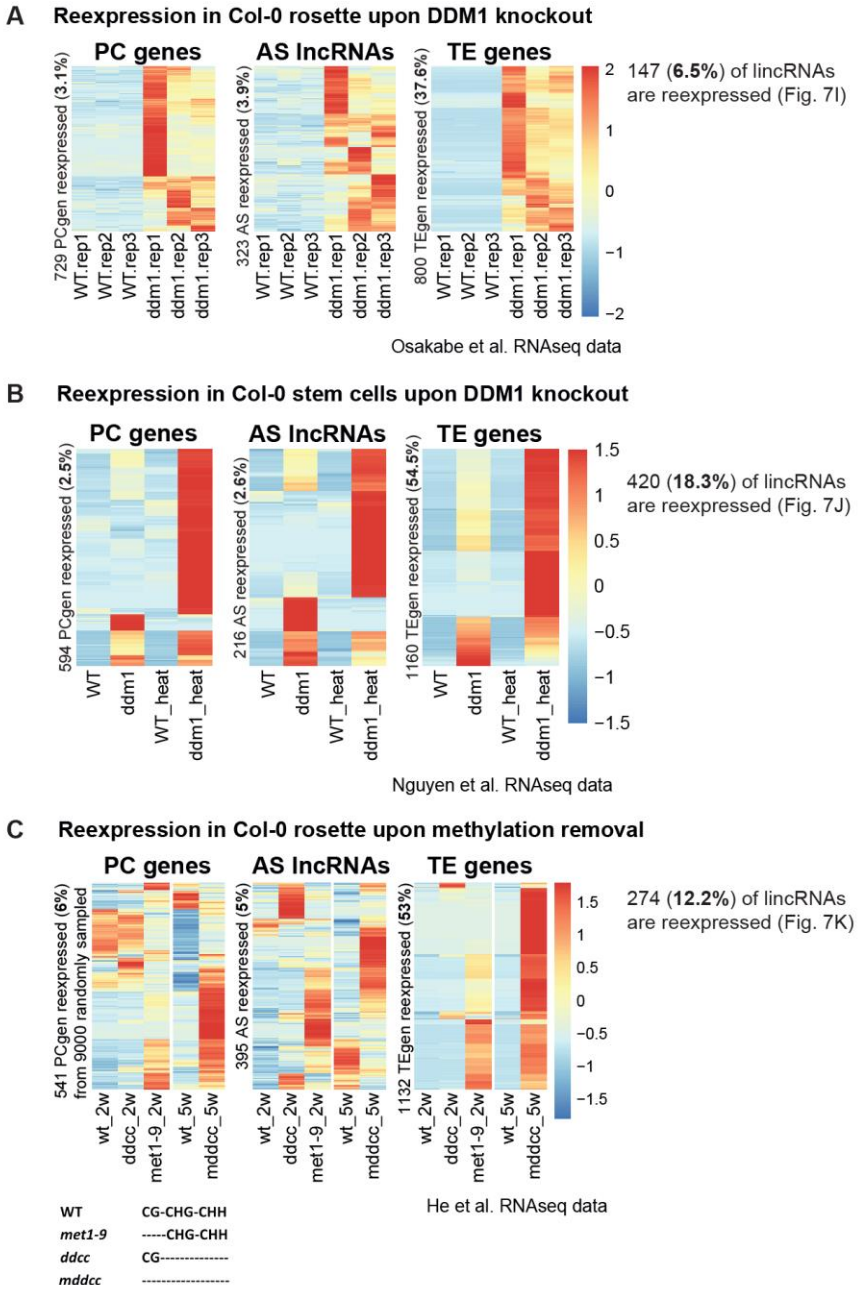


### Supplemental Figure S30. TE silencing mutants supplement

**A.** Expression level of PC gene, AS lncRNA and TE loci re-expressed in rosettes of the *ddm1* mutant in the Col-0 background (Osakabe et al. 2021). The re-expression was defined as no expression in the WT (TPM<0.5, average from three replicates) and expression in the mutant (TPM>0.5, MUT>3*WT, average from three replicates) (same for **B**). **B.** Expression level of PC gene, AS lncRNA and TE loci re-expressed in the *ddm1* mutant stem cells (Nguyen et al. 2023) **C.** Expression level of lincRNAs re-expressed in the rosettes of mutants of DNA methylases (He et al. 2022). A locus was considered re-expressed if it was re-expressed in one of the three mutants compared to the matching WT control. *ddcc* and *met1-9* mutants were 2-week-old seedlings and matched to the 2-week-old WT, and the *mddcc* mutant was matched to the 5-week-old WT. ***ddcc***: quadruple mutant for *DRM1*, *DRM2*, *CMT2*, and *CMT3*. ***mddcc***: quintuple mutant for *MET1*, *DRM1*, *DRM2*, *CMT2*, and *CMT3*. Heatmaps were built using “pheatmap” in R with scaling by row. No column clustering, row clustering trees not displayed.

**Supplemental Figure S31** (supports Figure 7)

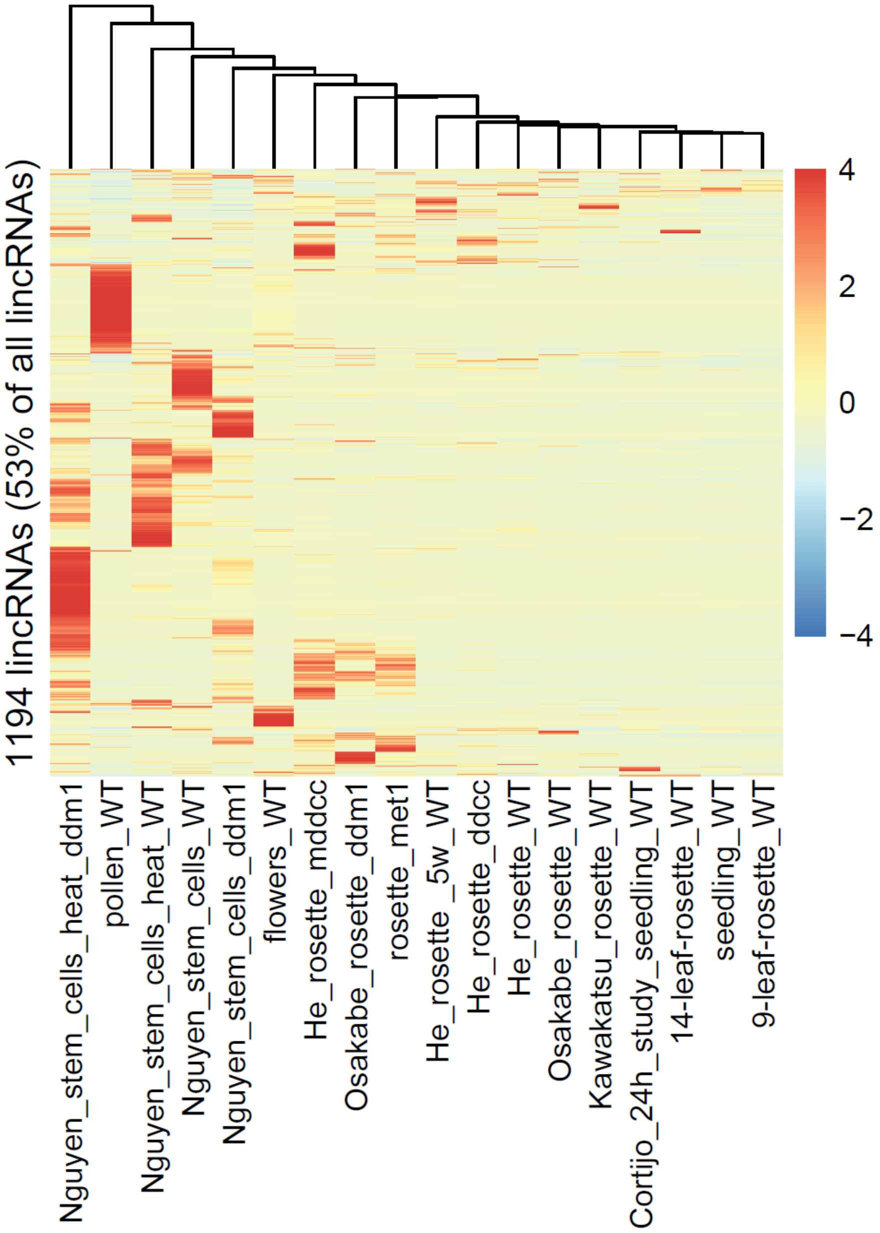


**Supplemental Figure S31. The scope of lincRNA expression potential in Col-0**

Heatmap showing the expression of lincRNA loci across the datasets used in our study. The heatmap was built using “pheatmap” in R with scaling by row. Rows clustered using “median” method in hclust, clustering tree not displayed. Only lincRNAs expressed in at least one sample (TPM>0.5) are plotted.

### **Supplemental Figure S32** (supports Figure 8)

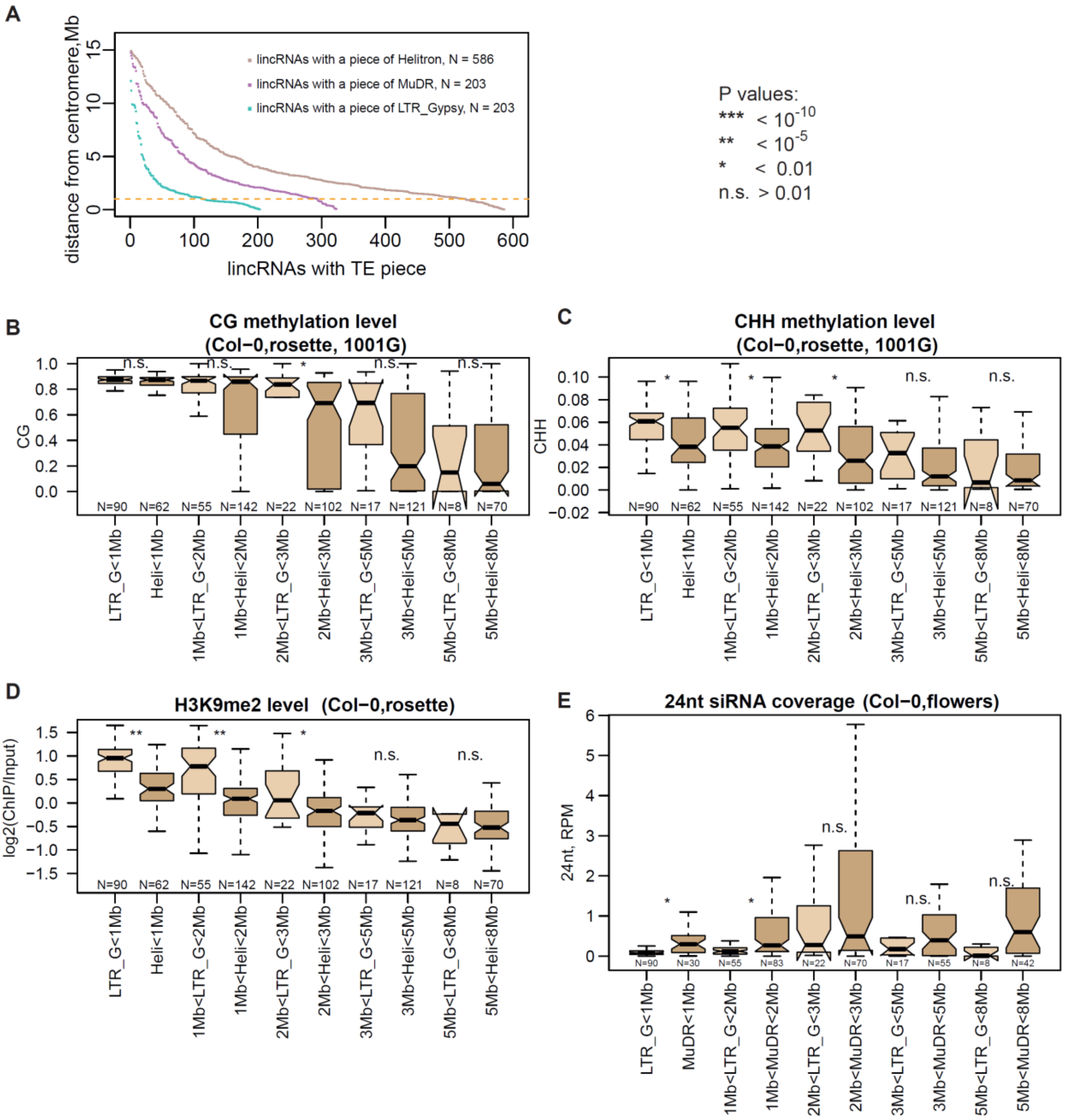


**Supplemental Figure S32. Genomic position and epigenetic pattern of lincRNAs with pieces of Class I and II TEs**

**A.** Distribution of lincRNAs with TE pieces from different TE families. Distance from the centromere vs the number of lincRNAs is plotted. LincRNAs are sorted by distance to the centromere. Only lincRNAs with TE pieces from only one TE type are plotted. **B, C, D.** CG methylation level in rosettes (**B**), CG methylation level in rosettes (**C**), H3K9me2 level in rosettes (**D**), for lincRNAs with either LTR_*Gypsy* or *Helitron* TEpieces split into five bins based on their distance to the centromeres. **E.** Coverage of 24-nt sRNA in flowers for lincRNAs with either LTR_*Gypsy* or DNA_*MuDR* TE-pieces split into five bins based on their distance to the centromeres. Only lincRNAs with TE-pieces of one type are plotted. Numbers below boxes indicate the number of lincRNA loci in each category. *P*-values were calculated using Mann-Whitney tests: *** $P < 10^{-10}$, ** $P < 10^{-5}$, * $P < 0.01$, n.s.: $P > 0.01$. Outliers in the boxplots are not shown. The centromere positions used: Chr1: 15,000,000; Chr2: 4,700,000, Chr3: 13,000,000, Chr4: 3,800,000, Chr5: 11,900,000. The methylation data displayed are from Col-0 rosettes from (Kawakatsu et al. 2016).

### **Supplemental Figure S33** (Supports Figure 8)

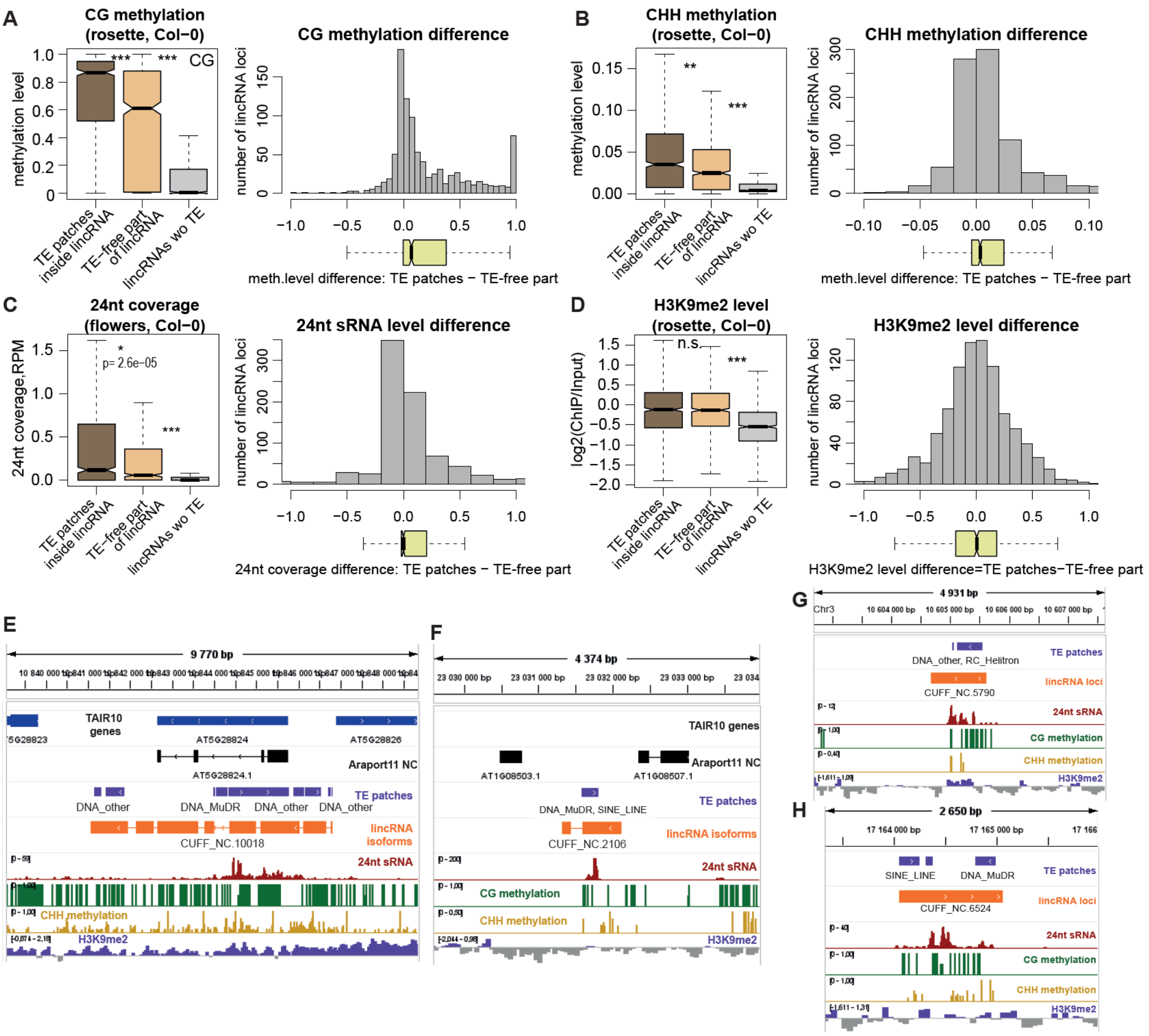


### **Supplemental Figure S33. Spreading of silencing from TE patches**

Boxplots showing CG methylation level (**A**), CG methylation level (**B**), 24-nt sRNA coverage (**C**) and H3K9me2 level (**D**) for TE patches within lincRNAs, TE-patch-free parts of TE-containing lincRNA loci and lincRNA loci without TE patches. The histograms show the distribution across all TE-containing lincRNAs of the following value: the epigenetic mark level at TE-patches minus the level at TE-free parts of the lincRNA locus. The boxplots below histograms show skewing towards positive values for CG methylation, CHH methylation and 24-nt sRNA coverage indicating that for many TE-containing lincRNA loci the levels of methylation and sRNA targeting are higher at the TE patch and lower outside of the patch suggesting the spreading of silencing. *P*-values were calculated using Mann-Whitney tests: *** $P < 10^{-10}$, ** $P < 10^{-5}$, * $P < 0.01$, n.s.: $P > 0.01$. Outliers in the boxplots are not shown. **E, F.** IGV screenshots showing examples of lincRNAs with TE patches that have higher level of CG methylation and 24-nt sRNA coverage over the TE patch than over the rest of the lincRNA locus. **G, H** IGV screenshots showing additional examples of lincRNAs with TE patches and epigenetic spreading outside the TE patches.

**Supplemental Figure S34** (Supports Figure 8)

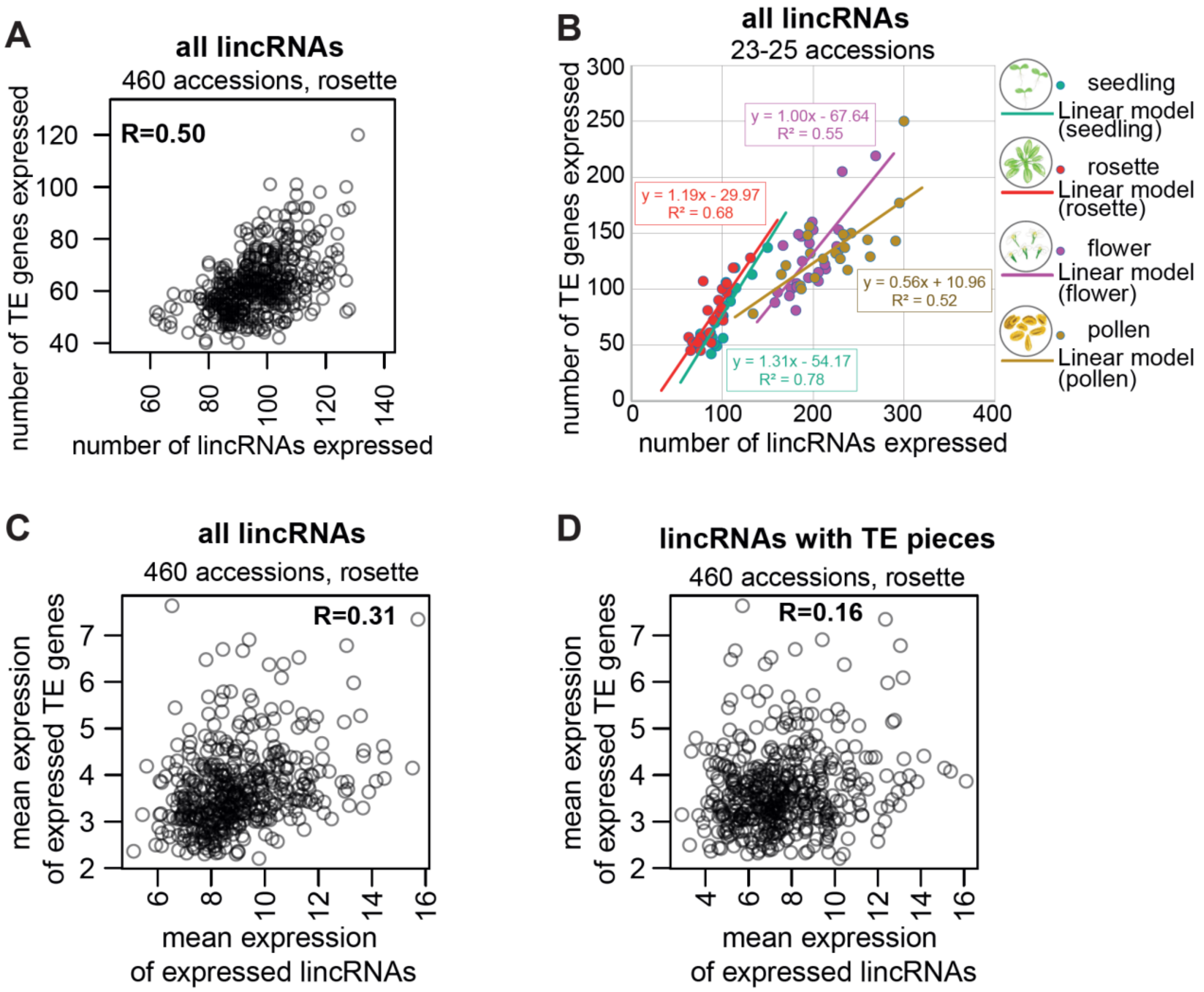


**Supplemental Figure S34. Variability of the number of TE genes and lincRNAs expressed**

**A.** Scatterplot showing the number of TE genes expressed in rosettes of 460 different accessions (Kawakatsu et al. 2016) as a function of the number of lincRNAs expressed in the same accession. (Accession 10010 that expressed twice as many TE genes was not included) **B.** Scatterplot showing the number of TE genes expressed in seedlings, 9-leaf rosettes, flowers, and pollen of 23–25 different accessions as a function of the number of lincRNAs expressed in the same accession and tissue. **C, D.** Scatterplot showing the average expression level of all expressed (TPM>0.5) TE genes expressed in rosettes of 460 different accessions (Kawakatsu et al. 2016) as a function of the average expression level of all lincRNAs (**C**) or all lincRNAs with TE pieces (**D**) expressed (TPM>0.5) in the same accession.

**Supplemental Figure S35** (Supports Figure 8)

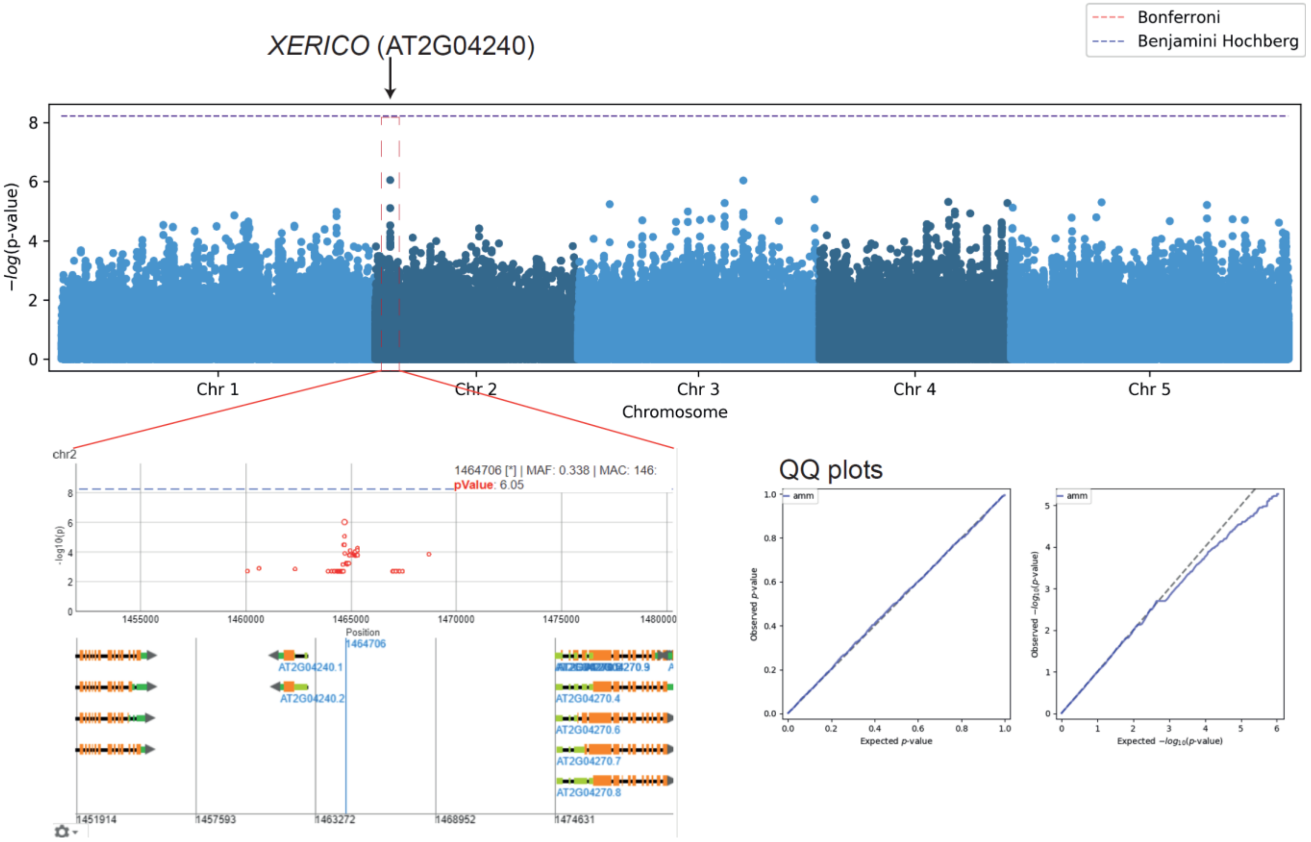


### Supplemental Figure S35. GWAS on expressed TE gene number

The Manhattan plot of the GWAS results on the Imputed Full-sequence dataset from the Arabidopsis 1001 Genomes Project population (461 accessions) using the number of TE genes expressed in each accession as phenotype. GWAS was performed and data visualized on the GWA-Portal (https://gwas.gmi.oeaw.ac.at/). Estimated pseudo-heritability: 0.87. Protocol used: AMM.

### **Supplemental Figure S36** (Supports Discussion)

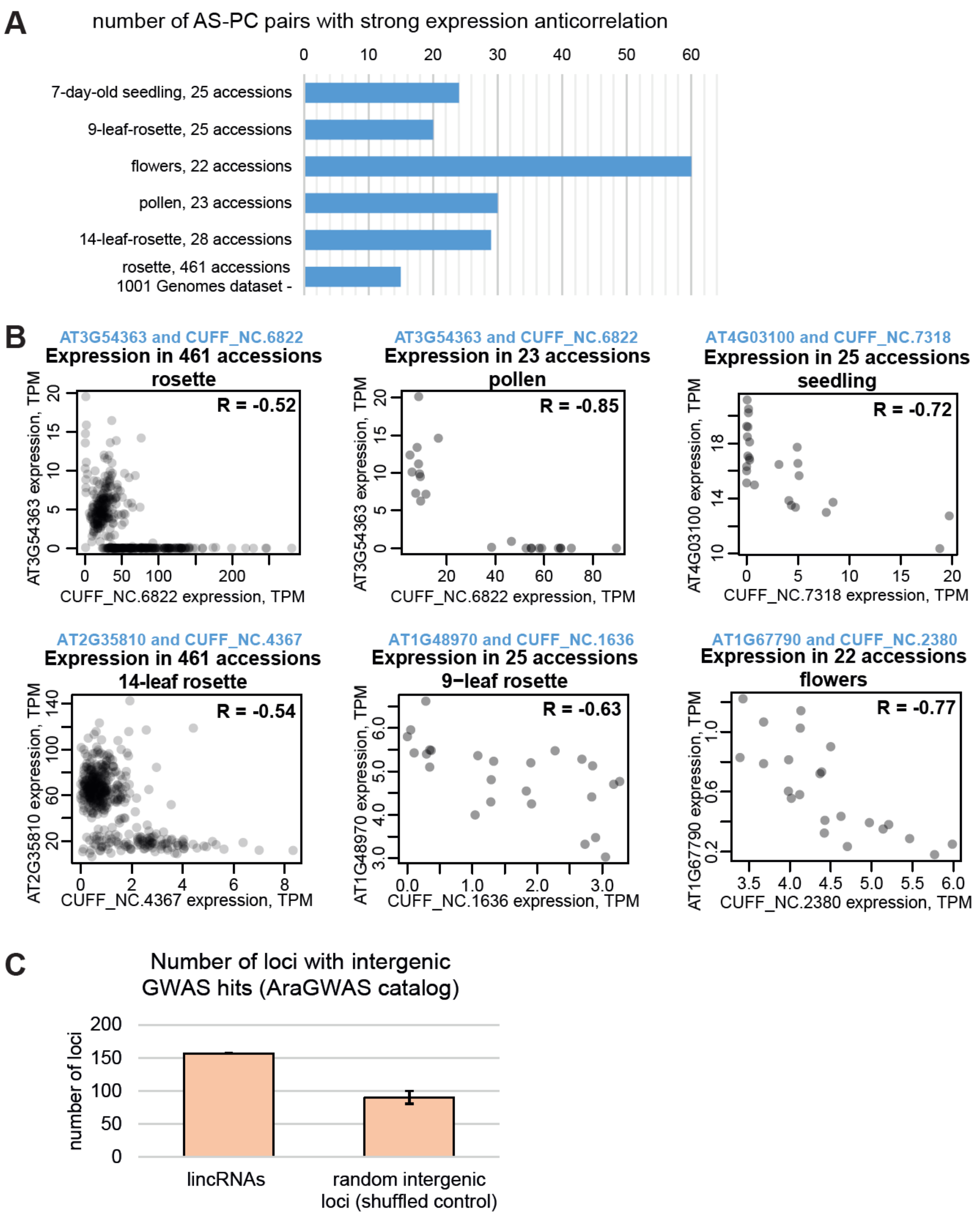


### **Supplemental Figure S36. lncRNA candidates**

**A.** The number of AS-lncRNA—PC-gene pairs that show negative correlation of their expression levels across different Arabidopsis accessions. A Pearson's correlation lower than –0.4 was considered strong. See the list of AS lncRNAs and the corresponding Araport11 PC genes in Supplemental Table S9. **B.** Examples of some candidate AS-PC pairs with strong negative correlation across accessions. Scatterplot showing the expression of the PC genes as a function of the expression level of its antisense lncRNA: each circle is one accession. Pearson correlation coefficient R is displayed. **C.** Number of lincRNA loci containing significant GWAS associations from the AraGWAS database (https://aragwas.1001genomes.org/) (Supplemental Table S10). Random intergenic controls were obtained by randomly shuffling the lincRNA loci from our annotation 8 times using bedtools shuffle -noOverlapping -seed $N -chromFirst -chrom -excl $denovo_pc -excl $PC_araport -excl $PC_tair -excl $TE_genes -i $denovo_linc -g TAIR10.chr_length_wo_ChrM_ChrC.txt > shuff.$N.bed. Error bars show standard deviation between 8 replicates.

### Supplemental Figure S37 (Supports Discussion)

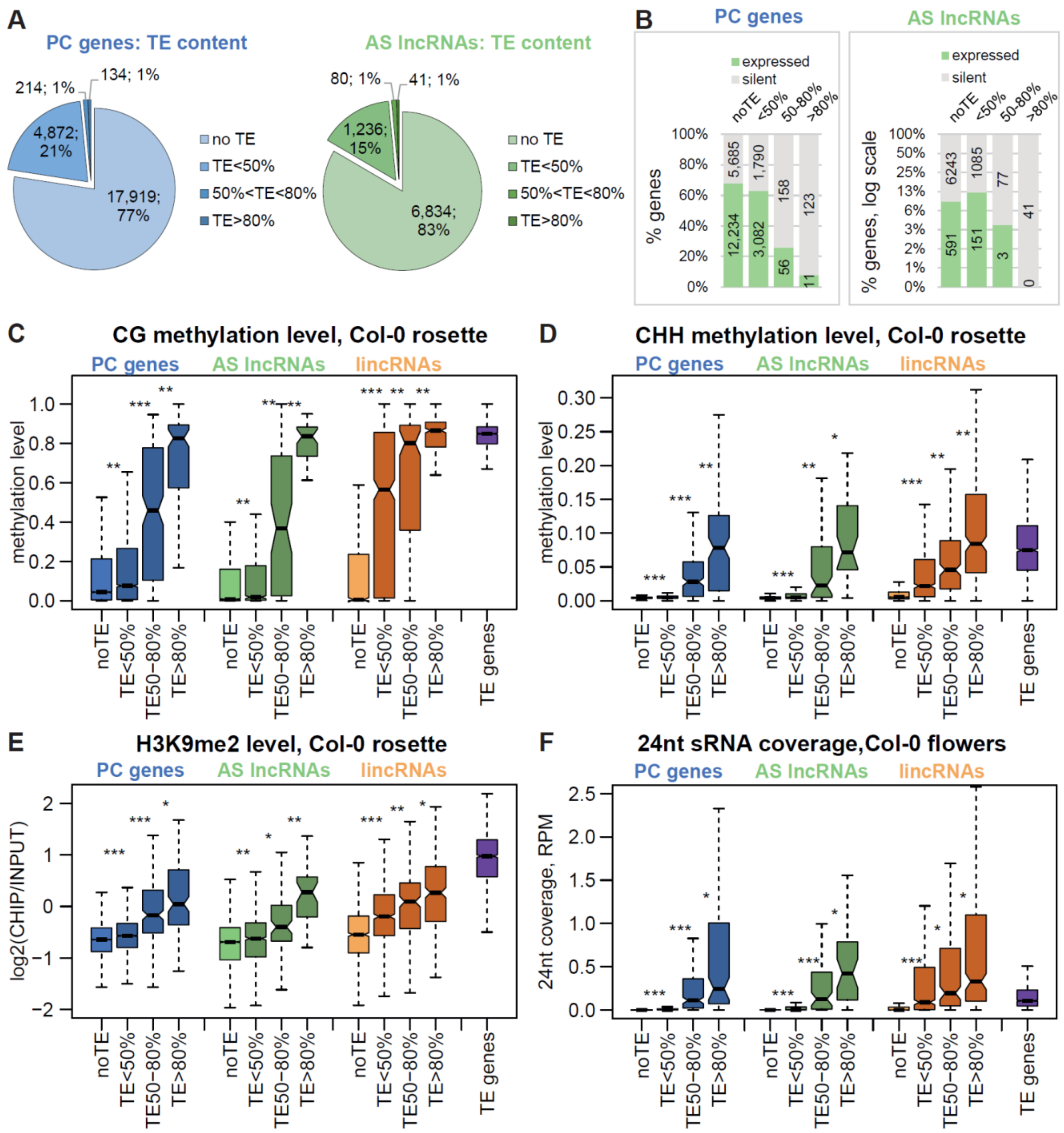


### Supplemental Figure S37. TE pieces affect expression as epigenetic patterns of AS lncRNAs and PC genes

**A.** Distribution of PC genes (left) and AS lncRNA loci (right) across four categories according to their TE content (TE-patches in any relative direction). **B.** Proportion of expressed PC genes (left) and AS lncRNAs (right) as a function of their TE content. The y-axis is displayed in log2 scale for AS lncRNAs. **C, D, E, F.** Levels of CG methylation (**C**), CHH methylation (**D**), H3K9me2 (**E**), and 24-nt sRNAs (**F**) for PC genes, AS lncRNAs and lincRNAs as a function of their TE content, with TE genes for comparison. *P*-values in the boxplots are calculated using Mann-Whitney test: *** $P < 10^{-10}$, ** $P < 10^{-5}$, * $P < 0.01$, n.s.: $P > 0.01$. Outliers in the boxplots are not plotted.

**Supplemental Figure S38** (Supports Discussion)

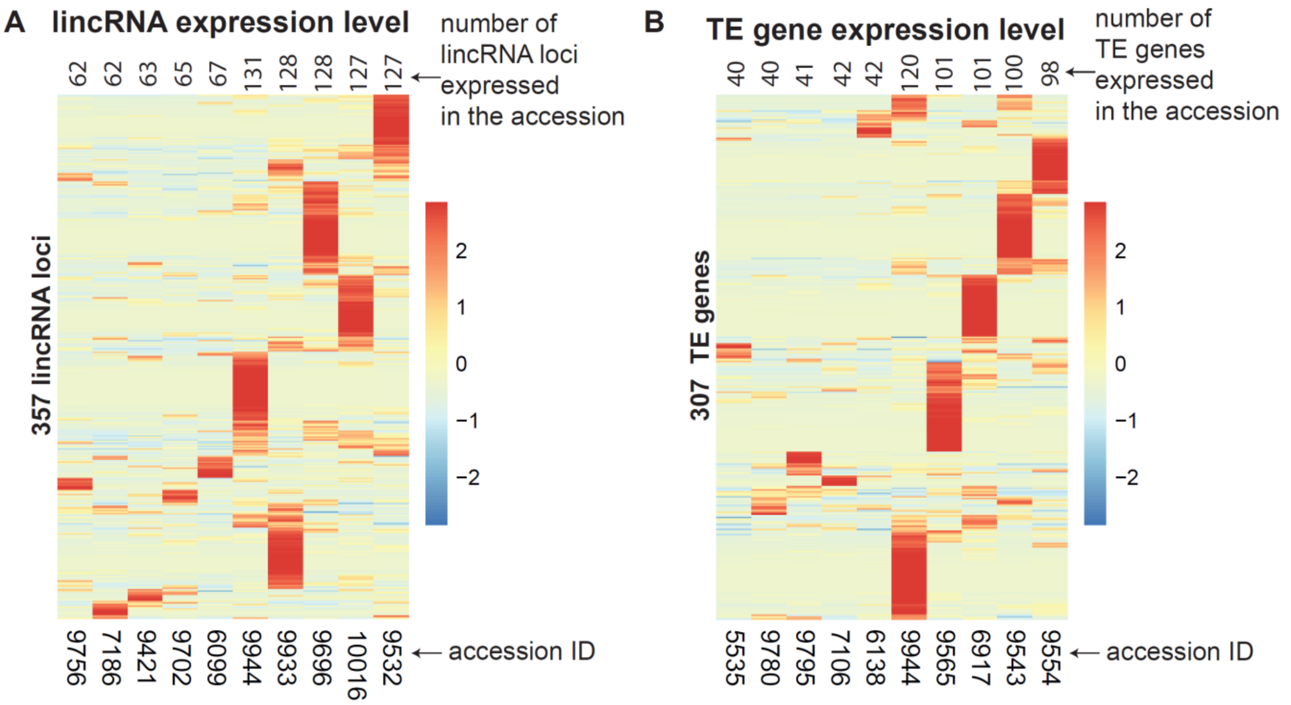


**Supplemental Figure S38. High expression variability despite similar expressed gene numbers**

Gene expression levels for (**A**) lincRNA loci and (**B**) TE genes in rosettes (Kawakatsu et al. 2016) for the five accessions with the smallest number of genes expressed and five accessions with the highest number of genes expressed (accession 10010 that expressed twice as many TE genes was not included). Heatmaps were built using “pheatmap” in R with scaling by row. Only genes expressed in at least one sample are plotted. Clustering trees for rows not shown.
